# Trophic diversity and evolution in Enantiornithes: a synthesis including new insights from Bohaiornithidae

**DOI:** 10.1101/2023.07.18.549506

**Authors:** Case Vincent Miller, Jen A. Bright, Xiaoli Wang, Xiaoting Zheng, Michael Pittman

## Abstract

The “opposite birds” Enantiornithines were the dominant birds of the Mesozoic, but our understanding of their ecology is still tenuous. In particular, diets of enantiornithine species have remained speculative until recently. While this new work has been effective at determining diet within groups of enantiornithines, diet data thus far has been too sparse to comment on larger trends in the diversity and evolution of enantiornithine ecology. We introduce new data on the enantiornithine family Bohaiornithidae, famous for their large size and strong teeth and claws. In tandem with previously-published data on the earlier-diverging pengornithids and later-diverging longipterygids, we comment on the breadth of enantiornithine ecology and potential patterns in which it evolved. Body mass, jaw mechanical advantage, finite element analysis of the jaw, and traditional morphometrics of the claws and skull are compared between bohaiornithids and living birds. The sample size for living bird body mass is over ten times larger than previous studies on longipterygid and pengornithid diet, with implications in interpreting their results. We find bohaiornithids to be ecologically diverse: *Bohaiornis* and *Parabohaiornis* are similar to living plant-eating birds; *Longusunguis* resembles raptorial carnivores; *Zhouornis* is similar to both fruit-eating birds and generalist feeders; and *Shenqiornis* and *Sulcavis* plausibly ate fish, plants, or a mix of both. This ecological diversity is wider than any other enantiornithine family studied previously, which may be driven by strengthening of the jaw relative to other early birds. This strong jaw would allow bohaiornithids to eat harder foods than other birds at the time, but their jaws were weaker than most “strong-jawed” living birds. With these reconstructions of diet in Bohaiornithidae, there is quantitative support for enantiornithines inhabiting nearly every trophic level. By combining these reconstructions with past dietary predictions for Longipterygidae and Pengornithidae, we predict the ancestral enantiornithine bird to have been a generalist which ate a wide variety of foods. This would suggest that the ecological diversity of enantiornithine birds represents specialisation in taking foods their ancestors were already eating, rather than many dramatic changes in diet. However, more quantitative data from across the enantiornithine tree is needed to refine this prediction. By the Early Cretaceous, enantiornithine birds had diversified into a variety of ecological niches in a similar way to crown birds after the K-Pg extinction, adding to the body of evidence that traits unique to crown birds (e.g. a toothless beak or cranial kinesis) cannot completely explain their ecological success.

## Introduction

The diet of crown birds is extremely broad, and dietary evolution within the crown is a complex mosaic [1]. However, dietary evolution in birds outside the crown is poorly understood, largely because the diet of most Mesozoic birds remains speculative [2]. Some recent studies have begun to elucidate this matter [3–7]. The most progress has been made among enantiornithines [8–10], the most abundant and speciose birds in the Cretaceous [11]. This progress only amounts to an examination of 12 of the over 100 described enantiornithine species [11], though. This has limited any large-scale examinations of the overall trophic diversity of enantiornithines and the patterns in which they diversified. Ideally, the next step to answering these questions is to examine a large group of enantiornithines which are phylogenetically intermediate to the previously-studied families. The enantiornithine family Bohaiornithidae (Fig. 1 centre) fits both of these requirements, and thus serves as an ideal stepping stone to a large-scale understanding of enantiornithine ecology.

**Figure 1.**
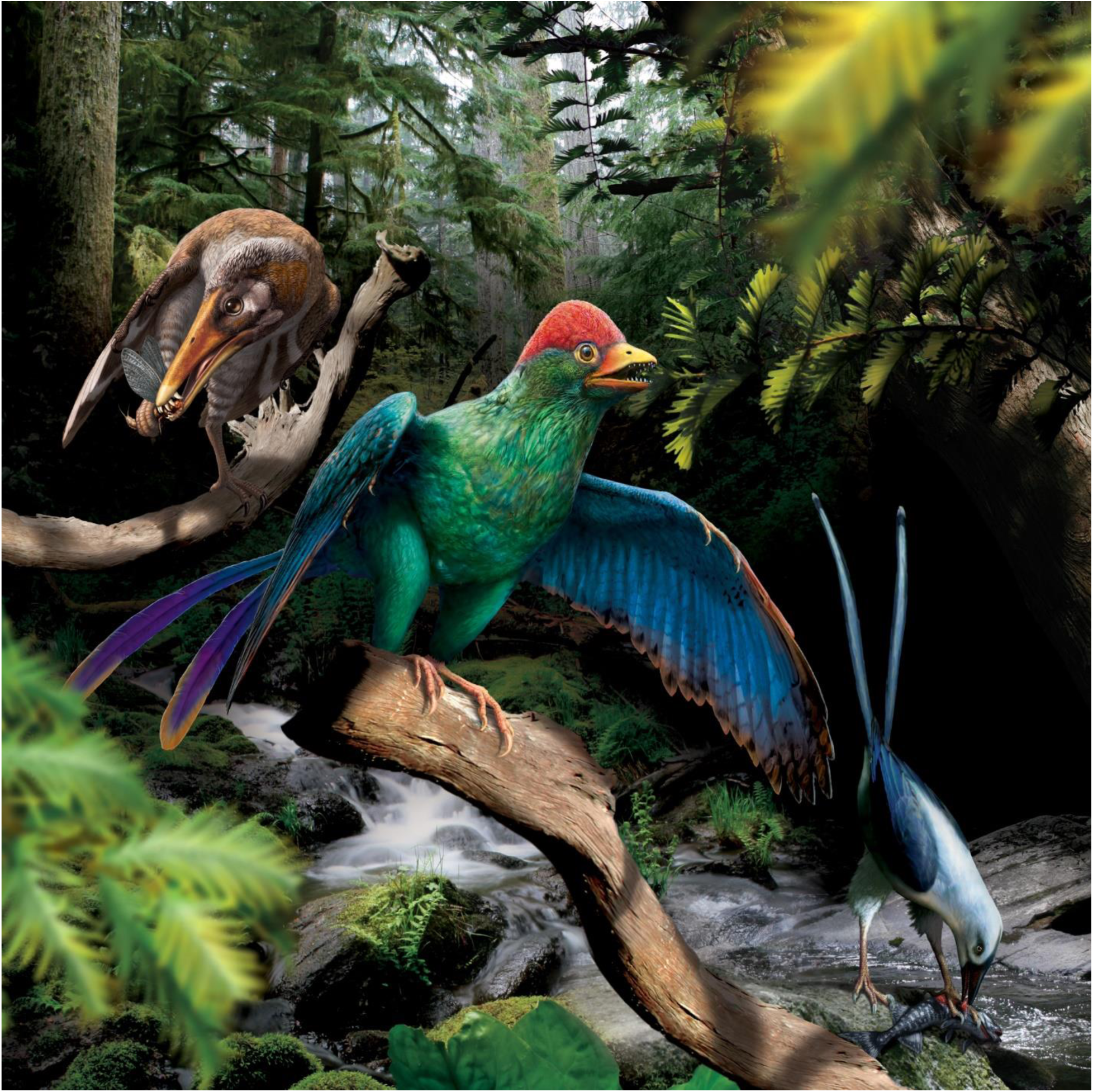
Life reconstruction of enantiornithine birds feeding. *Longipteryx* (left), *Bohaiornis* (centre), and *Pengornis* (right) are pictured in the Early Cretaceous forests of northeastern China, roughly 120 million years ago. *Bohaiornis* is depicted feeding on cypress (Cupressaceae, [150]) leaves after the findings in this work. *Longipteryx* is depicted feeding on the mayfly *Epicharmeropsis hexavenulosus* [151] after [8]. *Pengornis* is depicted feeding on the fish *Lycoptera davidi* [152] after [9].

Bohaiornithids are iconic among enantiornithines for their robust teeth, large claws, large size [12], and iridescent colouration [13]. Their robust teeth have led the clade to be interpreted as durophagous [14, 15, 16 pg. 83, 17], while their large claws and large body size lead to raptorial interpretations [12, 16 pg. 270, 18]. Bohaiornithidae has a troubled taxonomic history, with authors proposing the clade actually represents an evolutionary grade [19, 20] or at least a group in need of phylogenetic redefinition [21]. But across the literature the original six taxa referred to Bohaiornithidae (*Bohaiornis*, *Longusunguis*, *Parabohaiornis*, *Shenqiornis*, *Sulcavis*, and *Zhouornis* [12]; Fig. 2) have consistently resolved as closely-related taxa (Table S1), so for the purposes of this work we will refer to this group as a clade (see Methods for further justification). With the recently-described *Beiguornis* [22], Bohaiornithidae is the most speciose family of enantiornithine birds. Given their unusual morphology and apparent ecological success, Bohaiornithidae presents an ideal study topic for Mesozoic bird diet.

**Figure 2.**
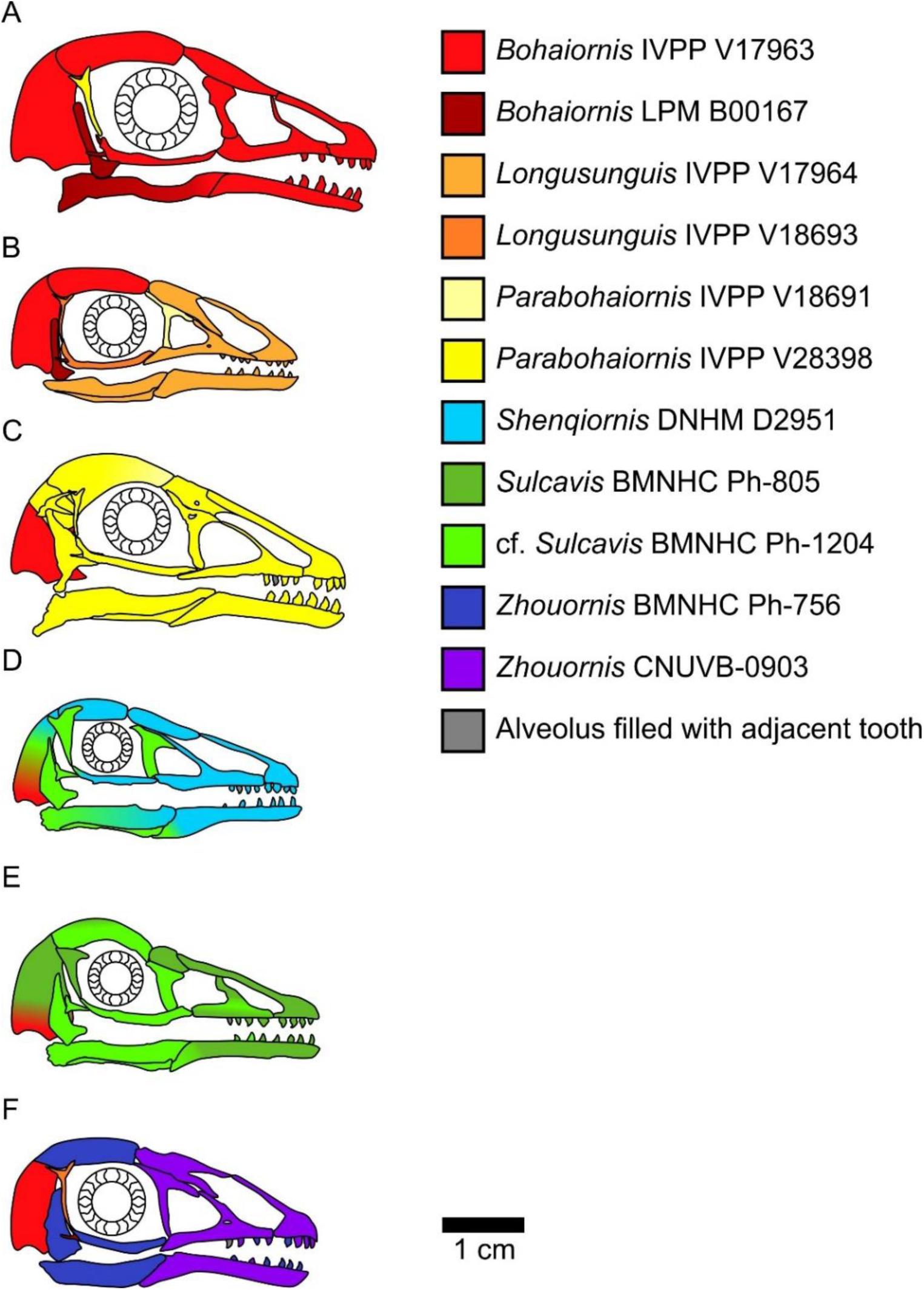
Bohaiornithid skull reconstructions used for MA and FEA calculations in this study. Reconstructions are of *Bohaiornis* (A), *Longusunguis* (B), *Parabohaiornis* (C), *Shenqiornis* (D), *Sulcavis* (E), and *Zhouornis* (F). Different colours indicate elements taken from different individual specimens. All sclerotic rings are based on *Longipteryx* specimen BMNHC Ph-930B. See the Methods section for more details on reconstruction. Scale for each reconstruction is based on the individual which makes up the largest portion of the reconstruction.

To investigate bohaiornithid diet, we utilise four quantitative diet proxies: body mass, mechanical advantage (MA) and related functional indices of the jaws, finite element analysis (FEA) modelling the jaws during a bite, and traditional morphometric (TM) analysis of claw shape and size. Size has a strong effect on birds’ diets [23–25], so estimating the mass of extinct birds [26] helps narrow dietary possibilities. Functional indices are ratios of measures of an animal that inform the mechanical efficiency of body parts to exert or withstand certain forces. Most commonly, the functional index used in ecology is MA of the jaw, looking at trade-offs between bite speed and force [27–29]. Herein three versions of MA are collected alongside three other functional indices which have previously discriminated animal diet [2, 30]. Finite element analysis is an engineering tool used to model forces acting on irregular structures [31]. Here it is used to model bird jaws during a bite. If a model experiences less strain under the same relative load as another model, its shape can withstand a greater force before failure. By maintaining a constant relative load for FEA models, models can be compared in terms of relative strength [32, 33]. TM describes the shape of animal parts with measurements relevant to their ecological role [34]. Here, measurements are taken of the curvature and relative size of pedal unguals which can differentiate between non-raptorial and raptorial birds [35–38] and, potentially, between specific types of raptorial behaviour [8, 36, 39].

These proxies are of little use without reference values, so comparative data is taken from nearly 200 extant birds [8, 9] including tinamous, flamingos, turacos, strisores, and songbirds among others. These birds are also ecologically diverse, with diet categories based on the EltonTraits 1.0 database of bird diet [40]. Table S2 provides cut-offs for diet assignment. Claw shape is not expected to correlate with diet but the use of talons, following the classification of [9] separating raptorial birds into raptors taking small prey (which can be completely encircled in the pes) and those hunting large prey (which cannot be encircled by the pes) based on feeding records in the Birds of the World database [41]. With the recent publication of the bird database AVONET [42], we are able to expand our dataset comparing mass and diet to 8,758 birds of the 9,994 recognized species of birds.

We expand the framework used in our past works [8, 9] by including traditional morphometrics of the skull. This line of evidence was recently used successfully in reconstructing the diet of longipterygid enantiornithines [10], so we expect it to be useful here as well. We incorporate the extant data from [10], collect new data for bohaiornithids, and analyse the data with a modified version of their methodology. Notably, we use alternate sources of body mass and diet data and adjust the data in an attempt to remove the effects of body size.

After each proxy is analysed, they are synthesised into a set of likely diets agreed upon by different evidence. Combining multiple lines of evidence allows for more precise and confident diet assignments than any single line can provide. Using this framework [2] we quantitatively test hypotheses of durophagy and raptorial behaviour in bohaiornithids.

Once dietary predictions are made for Bohaiornithidae, we can begin to examine large-scale trends in enantiornithine ecology via ancestral state reconstruction. Each of the diet proxies above can be predicted for the common ancestor of Enantiornithes and placed into the same framework as any individual species, to create a diet hypothesis for the common ancestor.

However, one would reasonably question if this sample size is large enough to intuit an ancestral diet reconstruction. To better visualize the uncertainty of our relatively small sample size, we also include a larger tree with qualitative diet assignments. This tree assumes published diet hypotheses are all correct as a test of what sampling density is necessary to produce confident ancestral state reconstructions of diet.

### Abbreviations

ACH, relative average cranial height; AMA, anterior jaw-closing mechanical advantage; AHM, relative average mandible height; AO, relative articular offset; BMNH, Beijing Museum of Natural History, Beijing, China; CI, confidence interval; CNUVB, Capital Normal University, Beijing, China; CUGB, China University of Geosciences, Beijing, China; DNHM, Dalian Natural History Museum, Dalian, China; FDA, flexible discriminate analysis; FEA, finite element analysis; HSD, honest significant differences; IVPP, Institute of Vertebrate Paleontology and Paleoanthropology, Beijing, China; LPM, Liaoning Paleontology Museum, Shenyang, China; MA, mechanical advantage; MCH, relative maximum cranial height; MHGU, The Museum of Hebei GEO University, Shijiazhuang, China; MMH, relative maximum mandibular height; PCA, principle component analysis; pFDA, phylogenetic flexible discriminant analysis; PMA, posterior jaw-closing mechanical advantage; TM, traditional morphometrics; MWAM, mesh-weighted arithmetic mean; µε, microstrain.

## Results

### Body Mass

Bird masses separated into 4 significantly different combinations of diet categories using phylogenetic honest significant differences (HSD) [8, 43]: a. nectarivores; b. granivores and invertivores; c. frugivores and generalists; and d. folivores and tetrapod hunters. Piscivores and scavengers were not significantly different from groups b or c. Optimizing the Youden index [44], cut-off points between these diet groups are as follows: between a and b, 9 g (95% confidence interval [CI] 8–10 g); between b and c, 56 g (95% CI 41–63 g); and between c and d, 265 g (95% CI 162–303 g). Mass of birds by diet and cut-off points between them are visualised in Fig. 2.

Estimated bohaiornithid body masses are provided in Table 1. Masses range from 91 g to 905 g, x̄ = 287 g, though the largest estimate (*Zhouornis* CNUVB-903) is an outlier. With the larger *Zhouornis* excluded, masses range from 91 to 406 g, x̄ = 248 g.

**Table 1.**
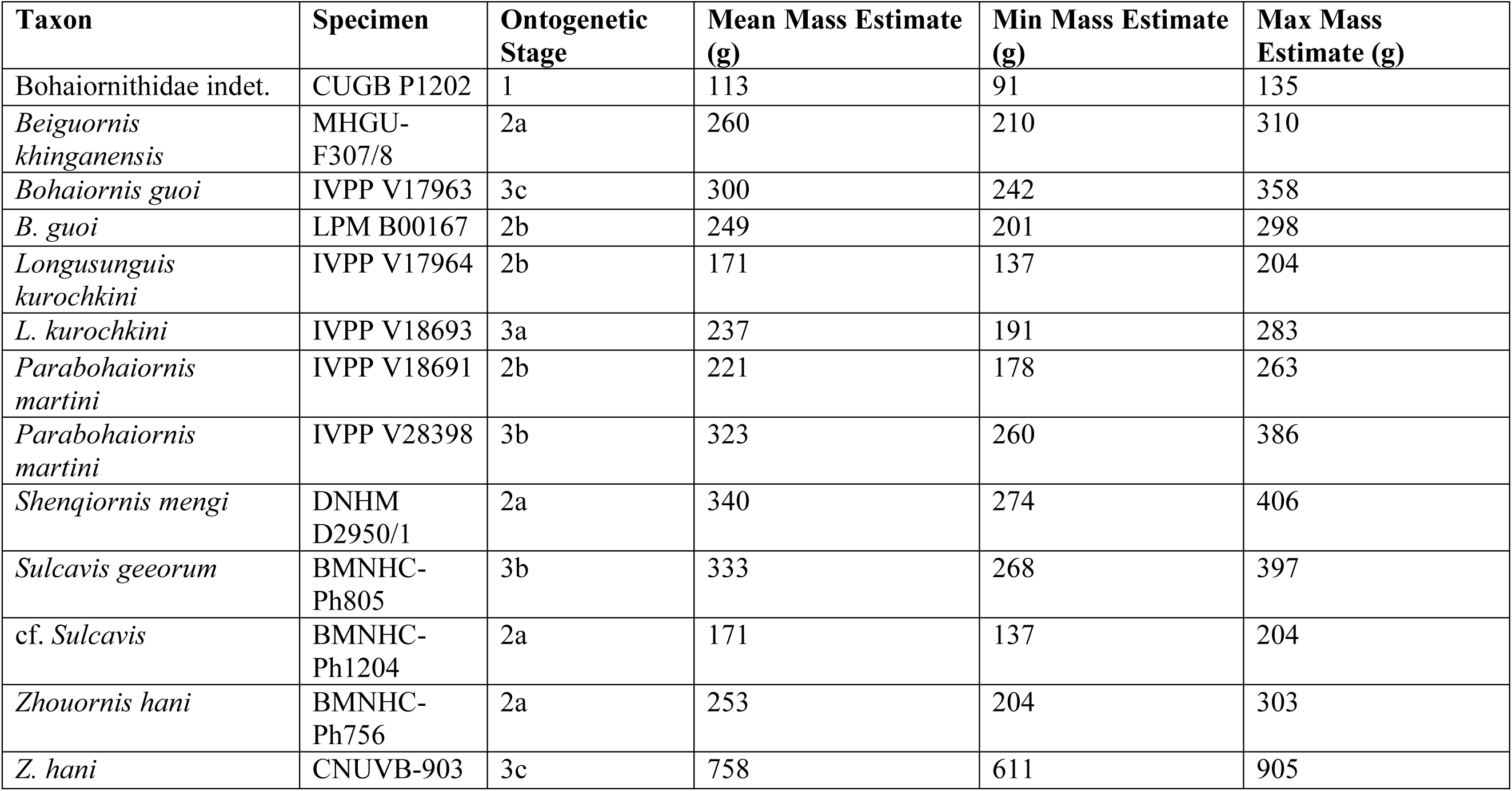
Masses for bohaiornithid taxa based on the regression equations of [26]. Most masses were previously reported in [2], though masses for the juvenile bohaiornithid CUGB P1202 [13], *Beiguornis khinganensis* MHGU-F307/8 [22], and cf. *Sulcavis* BMNHC-Ph1204 [21] are newly-calculated in this study from literature images.

| **Taxon** | **Specimen** | **Ontogenetic Stage** | **Mean Mass Estimate (g)** | **Min Mass Estimate (g)** | **Max Mass Estimate (g)** |
| --- | --- | --- | --- | --- | --- |
| Bohaiornithidae indet. | CUGB P1202 | 1 | 113 | 91 | 135 |
| *Beiguornis khinganensis* | MHGU-F307/8 | 2a | 260 | 210 | 310 |
| *Bohaiornis guoi* | IVPP V17963 | 3c | 300 | 242 | 358 |
| *B. guoi* | LPM B00167 | 2b | 249 | 201 | 298 |
| *Longsunguis kurochkini* | IVPP V17964 | 2b | 171 | 137 | 204 |
| *L. kurochkini* | IVPP V18693 | 3a | 237 | 191 | 283 |
| *Parabohaiornis martini* | IVPP V18691 | 2b | 221 | 178 | 263 |
| *Parabohaiornis martini* | IVPP V28398 | 3b | 323 | 260 | 386 |
| *Shenqiornis mengi* | DNHM D2950/1 | 2a | 340 | 274 | 406 |
| *Sulcavis georum* | BMNHC-Ph805 | 3b | 333 | 268 | 397 |
| cf. *Sulcavis* | BMNHC-Ph1204 | 2a | 171 | 137 | 204 |
| *Zhouornis hani* | BMNHC-Ph756 | 2a | 253 | 204 | 303 |
| *Z. hani* | CNUVB-903 | 3c | 758 | 611 | 905 |

### Mechanical Advantage and Functional Indices

Graphs of MA results are available in Fig. S1 (univariate) and Fig. 4 (multivariate), with 3D graphs in supplemental files. Posterior predictions of bohaiornithid diet from flexible discriminate analysis (FDA) are provided in Table 2. Extant MA and functional index results are unchanged from [9].

**Table 2.** Posterior probabilities predicting bohaiornithid diet by FDA from MA and functional indices of extant bird jaws. Values with green backgrounds are more likely, values with red backgrounds are less likely. All bohaiornithids have high affinity with generalists and low affinity with husking granivores, with other affinities varying by taxon. Diet abbreviations: GranivoreH, Husking Granivore; GranivoreS, Swallowing Granivore; Tetra Hunt, Tetrapod Hunter.

| Taxon | Folivore | Frugivore | Generalist | GranivoreH | GranivoreS | Invertivore | Nectarivore | Piscivore | Scavenger | Tetra Hunt |
| --- | --- | --- | --- | --- | --- | --- | --- | --- | --- | --- |
| *Bohaiornis* | 9.99E-01 | 6.43E-06 | 2.92E-04 | 5.97E-13 | 2.15E-04 | 5.43E-06 | 1.62E-05 | 2.00E-07 | 1.74E-10 | 7.78E-08 |
| *Longusunguis* | 1.53E-02 | 1.81E-01 | 5.39E-01 | 5.48E-07 | 4.38E-03 | 1.57E-01 | 3.05E-02 | 6.91E-02 | 9.85E-05 | 3.95E-03 |
| *Parabohaiornis* | 2.44E-01 | 6.72E-04 | 2.34E-01 | 2.49E-07 | 3.64E-01 | 1.05E-02 | 2.42E-04 | 1.49E-02 | 1.32E-01 | 7.69E-05 |
| *Shenqiornis* | 7.32E-03 | 1.12E-02 | 6.45E-01 | 1.81E-06 | 2.00E-03 | 5.15E-02 | 8.12E-03 | 2.65E-01 | 9.50E-03 | 5.24E-04 |
| *Sulcavis* | 7.28E-02 | 1.58E-02 | 8.22E-01 | 5.46E-07 | 4.12E-03 | 2.43E-02 | 3.39E-03 | 5.68E-02 | 5.74E-04 | 6.25E-04 |
| *Zhouornis* | 3.20E-02 | 1.61E-01 | 3.11E-01 | 9.82E-07 | 2.30E-02 | 2.69E-01 | 5.42E-02 | 7.86E-02 | 9.64E-04 | 6.99E-02 |

Bohaiornithids generally have low anterior jaw-closing mechanical advantage (AMA), posterior jaw-closing mechanical advantage (PMA), relative articular offset (AO), and relative maximum mandibular height (MMH) relative to living birds (Fig. S1A–D,G,H,J). Most bohaiornithids also have a low jaw-opening mechanical advantage (OMA), but *Bohaiornis* has a high OMA. Other functional indices are intermediate across bohaiornithids.

In principle component analysis (PCA), *Longusunguis*, *Shenqiornis*, and *Sulcavis*, all plot near one another in a region inhabited by invertivores and generalists. This is driven by their low jaw-closing MA (Fig. S2A). *Parabohaiornis*, with a slightly higher jaw-closing MA, plots nearby but closer to frugivores. *Bohaiornis* plots in an unoccupied region nearest frugivores and tetrapod hunters, separating from other bohaiornithids by its high upper jaw OMA (Fig. S2A). *Zhouornis* plots between *Bohaiornis* and other bohaiornithids near invertivores and granivores due to its high average cranial height (ACH).

In flexible discriminant analysis (FDA), bohaiornithids other than *Bohaiornis* plot together in an indeterminate region inhabited by all diets but folivores and husking granivores. This mirrors their high affinity with generalist feeders (Table 2). *Bohaiornis* plots far from other bohaiornithids and the extant-inhabited area of the functional phylomorphospace along discriminant axis 2 (Fig. S2B). The reason for this is unclear. The only functional index in which *Bohaiornis* differs from other bohaiornithids is OMA (Fig. S1), which is loaded primarily on discriminant axis 1 (Fig. S2B). Phylogenetic flexible discriminate analysis (pFDA) does not provide meaningful results when applied to functional index data, likely due to its poor explanatory power of the extant data [9].

### Finite Element Analysis

Graphs of FEA results are available in Fig. 5 (mesh-weighted arithmetic mean [MWAM] strain [45]) and Fig. 6 (multivariate strain intervals [46]), with 3D graphs in supplemental files. Posterior predictions of bohaiornithid diet from FDA are provided in Table 3. Extant FEA results are unchanged from [9].

**Table 3.** Posterior probabilities predicting bohaiornithid diet by FDA from FEA following the intervals method [46]. Values with green backgrounds are more likely, values with red backgrounds are less likely. Bohaiornithid affinities varying considerably between taxa, only universally not resembling tetrapod hunters. Diet abbreviations: GranivoreH, Husking Granivore; GranivoreS, Swallowing Granivore; Tetra Hunt, Tetrapod Hunter

| Taxon | Folivore | Frugivore | Generalist | GranivoreH | GranivoreS | Invertivore | Nectarivore | Piscivore | Scavenger | Tetra Hunt |
| --- | --- | --- | --- | --- | --- | --- | --- | --- | --- | --- |
| *Bohaiornis* | 3.44E-10 | 9.33E-12 | 1.73E-07 | 2.25E-23 | 1.00E+00 | 7.09E-07 | 4.67E-19 | 1.43E-10 | 8.82E-06 | 3.37E-19 |
| *Longusunguis* | 3.05E-01 | 1.15E-13 | 6.95E-01 | 1.56E-26 | 2.26E-15 | 3.58E-09 | 4.35E-26 | 1.51E-11 | 3.47E-29 | 1.07E-20 |
| *Parabohaiornis* | 8.49E-02 | 3.30E-15 | 2.08E-13 | 3.50E-24 | 5.52E-01 | 1.18E-04 | 4.67E-04 | 3.63E-01 | 3.12E-13 | 1.13E-06 |
| *Shenqiornis* | 2.28E-08 | 1.45E-23 | 9.39E-09 | 4.31E-11 | 2.28E-13 | 2.47E-05 | 3.61E-02 | 9.58E-01 | 6.11E-03 | 2.81E-20 |
| *Sulcavis* | 6.82E-06 | 1.91E-16 | 1.19E-03 | 9.94E-01 | 5.99E-18 | 5.65E-09 | 4.71E-03 | 1.22E-07 | 4.68E-35 | 1.55E-22 |
| *Zhouornis* | 5.69E-50 | 1.00E+00 | 5.55E-20 | 4.89E-13 | 4.06E-12 | 2.08E-18 | 8.99E-13 | 4.74E-20 | 1.13E-53 | 2.52E-17 |

MWAM strain of bohaiornithids is low (89–156 µε, x̄ = 128 µε), with *Bohaiornis* and *Zhouornis* experiencing less strain under loading that any extant carnivorous bird in this study (min = 105 µε).

In PCA, *Bohaiornis* inhabits a region of the strain-space that is only inhabited by extant herbivores and *Zhouornis* inhabits a space that is exclusively non-carnivorous. *Longusunguis*, *Parabohaiornis*, and *Sulcavis* inhabit a region where all diets overlap. *Shenqiornis* inhabits a nearby region with slightly less occupation by invertivores, piscivores, and frugivores.

Bohaiornithids tend to display heterogeneous strain, i.e. more model area under high or low strain rather than under intermediate strain.

In FDA, bohaiornithids tend to plot in uninhabited areas of the strain-space (Fig. 6B). Thus, the high confidence of posterior predictions from FEA (Table 3) reflect the nearest extant diet group. As previously noted for this dataset [8], groups with similar distance from the origin are not meaningfully different in jaw strength, so all bohaiornithids, not just specific taxa, should be interpreted as plotting in the same functional space as folivores, frugivores, nectarivores, piscivores, and scavengers. The exception is *Longusunguis*, the bohaiornithid with the second weakest jaw whose affinities are strongest with generalist feeders. pFDA does not provide meaningful results when applied to FEA intervals data, likely due to the low phylogenetic signal in these data [9].

### Traditional Morphometrics

#### Pes

Graphs of pedal TM results are available in Fig. 7, with character weights in Fig. l and 3D graphs in supplemental files. Posterior predictions of bohaiornithid pedal ecology from FDA and pFDA are provided in Table 4. Extant pedal TM results are unchanged from [9]. Bohaiornithids generally spread across the phylomorphospace. In PCA, *Sulcavis* plots among large raptors and both *Parabohaiornis* individuals plot among non-raptorial perching birds (though the latter separate into an adjacent uninhabited space with PC3 included, see supplemental files). *Longusunguis* IVPP V18693, both specimens of *Bohaiornis*, and both specimens of *Zhouornis* plot in an indeterminate region of primarily small raptors and non-raptorial perching birds. *Longusunguis* in particular also plots near tinamous, which are ground birds.

**Figure 3.**
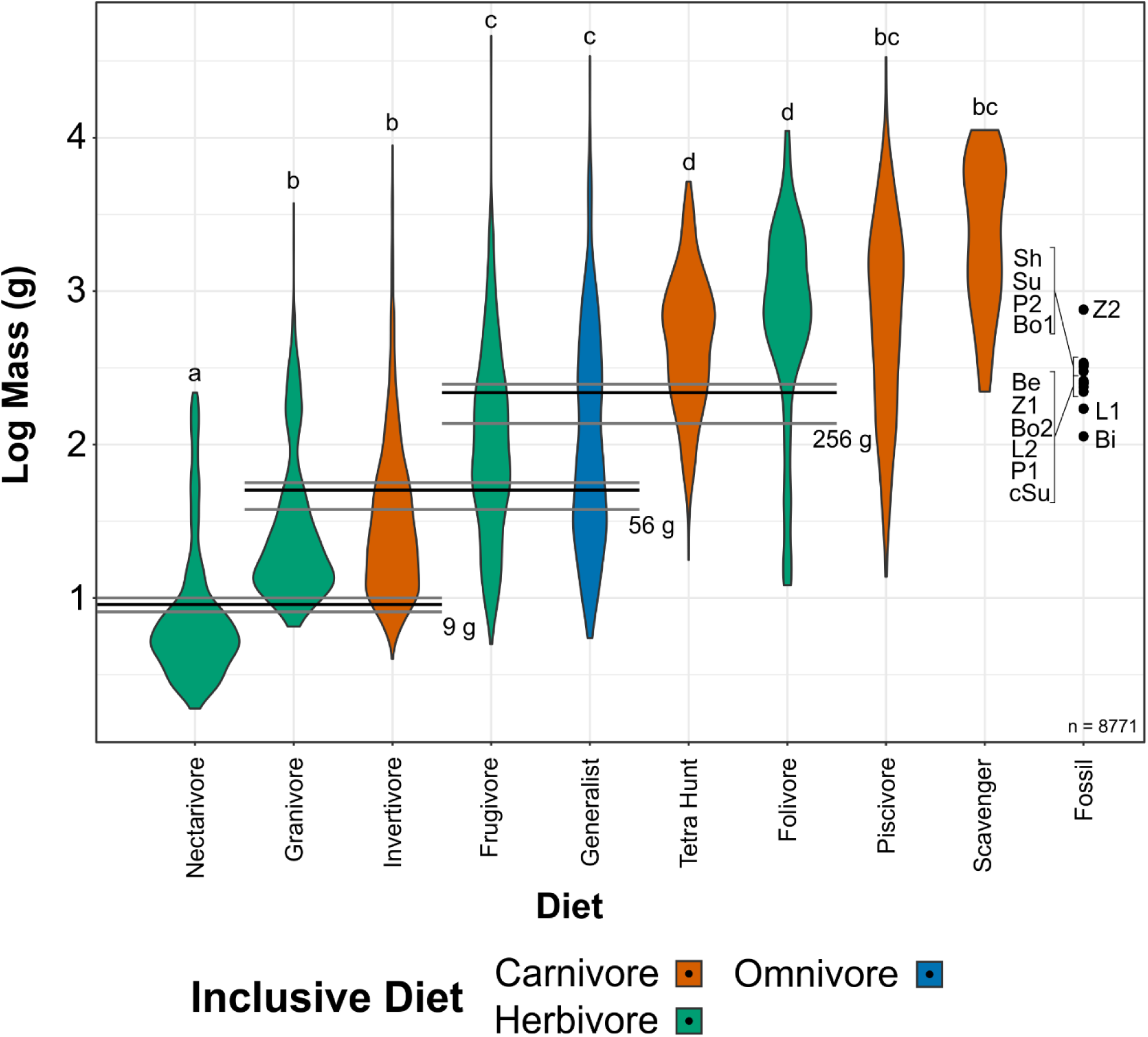
Violin plots of bird mass by diet, arranged in order of ascending mean mass. Masses were tested for significant differences via phylogenetic HSD. Diets marked with the same letter are not significantly different from one another. Cut-off points between significantly different mass groups (black lines, with 95% CIs as grey lines) were calculated by optimizing the Youden index and plotted. Note that, unlike in other diet treatments herein, granivores are not separated into husking and swallowing granivores. Mean bohaiornithid mass estimates are plotted for comparison, see Table 1. Diet abbreviations: Tetra Hunt, Tetrapod Hunter. Fossil taxon abbreviations: Be, *Beiguornis khinganensis* MHGU-F307/8; Bi, Bohaiornithidae indet. CUGB P1202; Bo1, *Bohaiornis* LPM B00167; Bo2, *Bohaiornis* IVPP V17963; L1, *Longusunguis* IVPP V17964; L2, *Longusunguis* IVPP V18693; P1, *Parabohaiornis* IVPP V18691; P2, *Parabohaiornis* IVPP V28398; Sh, *Shenqiornis* DNHM D2950/1; Su, *Sulcavis* BMNH Ph-805; cSu, cf. *Sulcavis* BMNHC-Ph1204; Z1, *Zhouornis* CNUVB-0903, Z2, *Zhouornis* BMNHC Ph 756.

**Table 4.** Posterior probabilities predicting bohaiornithid pedal ecology by FDA and pFDA from TM of extant bird claws. Values with green backgrounds are more likely, values with red backgrounds are less likely. Bohaiornithid affinities vary considerably by taxon and between FDA and pFDA. Diet abbreviations: GranivoreH, Husking Granivore; GranivoreS, Swallowing Granivore; Tetra Hunt, Tetrapod Hunter. Category abbreviations: Large Raptor, raptor taking prey which does not fit in the foot; Small Raptor, raptor taking prey which can fit in the foot.

| Test | Taxon | Ground | Perch | Large Raptor | Small Raptor | Shrike |
| --- | --- | --- | --- | --- | --- | --- |
| FDA | *Bohaiornis*<br>LPM B00167 | 2.45E-02 | 4.03E-01 | 2.69E-01 | 3.03E-01 | 4.60E-04 |
|  | *Bohaiornis*<br>IVPP V17963 | 1.92E-02 | 2.52E-01 | 2.31E-04 | 7.23E-01 | 5.78E-03 |
|  | *Longusunguis* | 3.65E-02 | 4.93E-01 | 3.76E-01 | 9.45E-02 | 4.42E-07 |
|  | *Parabohaiornis*<br>IVPP V18690 | 2.94E-05 | 8.57E-01 | 1.49E-03 | 5.57E-02 | 8.63E-02 |
|  | *Parabohaiornis*<br>IVPP V18691 | 2.41E-05 | 9.66E-01 | 2.66E-02 | 4.41E-03 | 2.85E-03 |
|  | *Sulcavis* | 2.03E-01 | 1.80E-02 | 2.17E-01 | 5.62E-01 | 2.93E-09 |
|  | *Zhouornis* CNUVB-0903 | 1.22E-01 | 3.62E-01 | 6.10E-02 | 4.07E-01 | 4.83E-02 |
|  | *Zhouornis* BMNHC Ph 756 | 5.46E-01 | 1.41E-02 | 1.72E-02 | 4.22E-01 | 1.33E-04 |
| pFDA | *Bohaiornis*<br>LPM B00167 | 0.00E+00 | 6.09E-02 | 1.29E-01 | 8.10E-01 | 4.57E-05 |
|  | *Bohaiornis*<br>IVPP V17963 | 0.00E+00 | 6.21E-07 | 4.58E-06 | 4.87E-12 | 1.00E+00 |
|  | *Longusunguis* | 0.00E+00 | 2.66E-03 | 9.94E-01 | 2.82E-03 | 6.46E-04 |
|  | *Parabohaiornis*<br>IVPP V18690 | 0.00E+00 | 1.98E-04 | 2.10E-01 | 7.90E-01 | 3.27E-12 |
|  | *Parabohaiornis*<br>IVPP V18691 | 0.00E+00 | 2.68E-02 | 3.05E-01 | 6.68E-01 | 3.28E-07 |
|  | *Sulcavis* | 0.00E+00 | 2.87E-04 | 3.92E-05 | 3.27E-05 | 1.00E+00 |
|  | *Zhouornis* CNUVB-0903 | 0.00E+00 | 4.94E-01 | 2.84E-01 | 2.19E-01 | 3.00E-03 |
|  | *Zhouornis* BMNHC Ph 756 | 0.00E+00 | 3.10E-01 | 5.23E-03 | 3.96E-03 | 6.80E-01 |

In FDA, *Parabohaiornis* shows a consistent affinity with non-raptorial perching birds (Table 4). Both specimens of *Bohaiornis* and the younger specimen of *Zhouornis* show affinity with small raptors as well as non-raptorial perching birds, with the older *Zhouornis* showing more affinity with ground birds than perching birds (Table 4). *Longusunguis* and *Sulcavis* both specimens show affinity with large raptors, with *Longusunguis* also similar to non-raptorial perchers and *Sulcavis* to small raptors (Table 4).

While pFDA results differ from [9] due to the use of a new phylogeny and additional extant data (see Methods), the differences are minimal. The new optimal λ of the extant data is 0.33 (optimal λ was 0.31 in [9]), and axis loadings are nearly identical (Fig. S3C). In pFDA, *Bohaiornis* LPM B00167 shows affinity with small raptors, *Longusunguis* with large raptors, and *Zhouornis* CNUVB-903 with non-raptorial perching birds (Table 4). The remaining bohaiornithid fossils plot outside the phylomorphospace occupied by extant taxa (Fig. 7C).

#### Skull

A graph of skull TM as described in its original application to enantiornithines [10] is available in Fig. S4A. However, as seen in Fig. S4B and discussed at length in the Methods, skull length appears to be a size proxy that is redundant given the mass data examined in this work. Instead, we compare relative rostrum length as used in [10] to relative skull length (i.e. skull length normalised to body size; Fig. 8). Bohaiornithids display distinctly short skulls (relative skull length < 1.6), with *Zhouornis* displaying the shortest skull relative to body mass of any taxon sampled. Their relative rostrum lengths fall in the middle of the extant range.

### Ancestral State Reconstruction

Qualitative ancestral state reconstruction of enantiornithine diet is presented in Fig. 9. Numerical likelihoods of each node are available in the supplemental information. Quantitative ancestral states are provided for body mass (Fig. S5), MA and functional indices (Fig. S6), FEA MWAM strain (Fig. S7), and pedal TM (Fig. S8). Due to software limitations, these presentations have polytomies non-randomly resolved and thus are only one possible reconstruction. Table 5 provides more valid values of each quantitative trait for the common ancestor of Enantiornithes by averaging results from 10,000 random polytomy resolutions. To facilitate interpreting multivariate traits of the common ancestor of Enantiornithes, FDA posterior probabilities are provided in Table 6 for the common ancestor MA data in Table 5, and a PCA graph of the common ancestor pedal TM data is provided in Fig. S9.

**Table 5.**
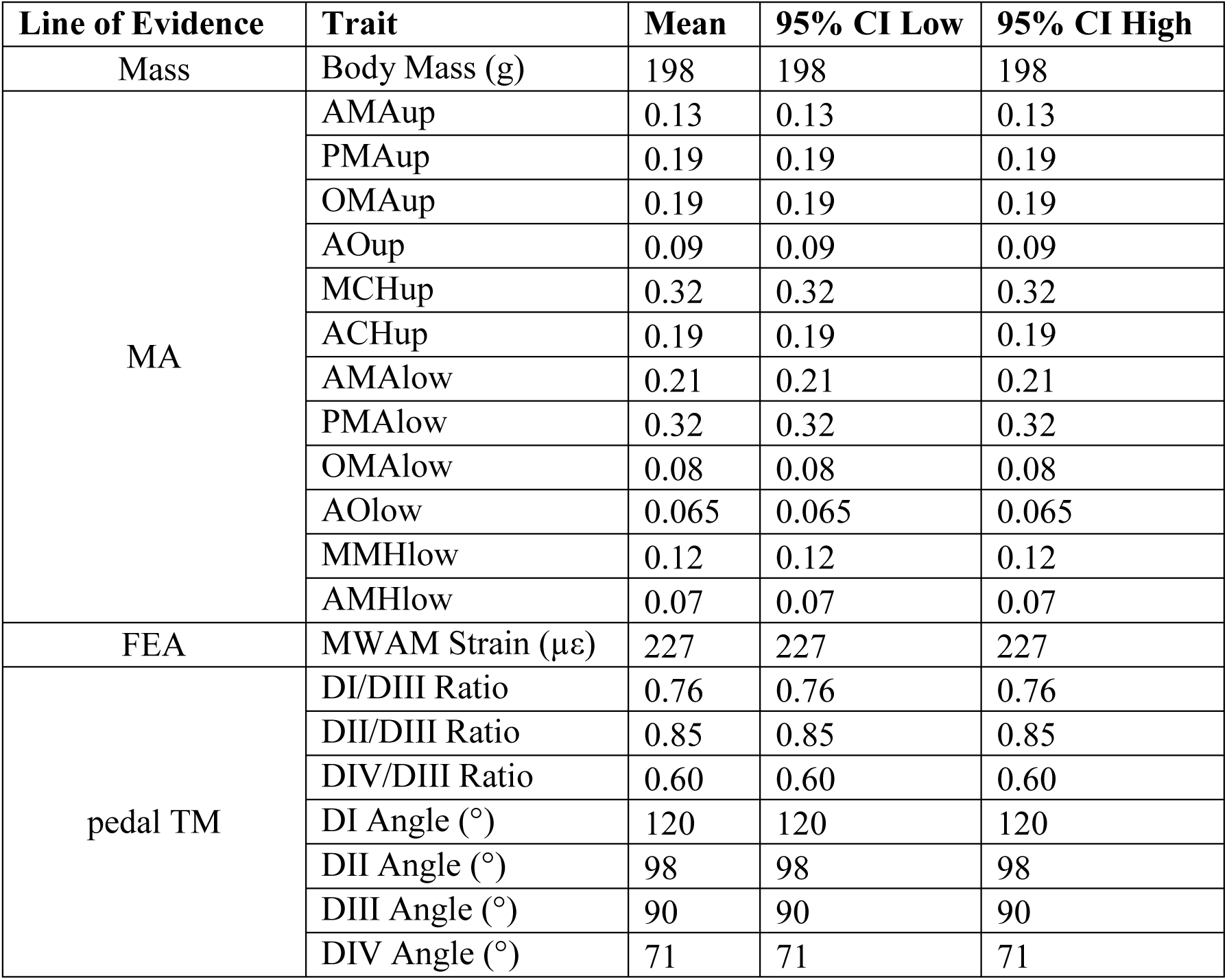
Quantitative ancestral states of the common ancestor of Enantiornithines. Mean values are the average of 10,000 random tree permutations where polytomies are randomly resolved. 95% confidence intervals are given as the 2.5% and 97.5% quantiles of the permutations, though upper and lower bounds for all values are identical to seven significant figures. As these values are based on multiple possible topologies of the enantiornithine tree, we consider them more valid than those visualised in Figs. 12–16.

**Table 6.** Posterior probability of diet in the common ancestor of Enantiornithes based on MA and functional indices. Generalist feeding is the most likely, followed by piscivory and invertivory. Diet abbreviations: GranivoreH, Husking Granivore; GranivoreS, Swallowing Granivore; Tetra Hunt, Tetrapod Hunter.

| Folivore | Frugivore | Generalist | GranivoreH | GranivoreS | Invertivore | Nectarivore | Piscivore | Scavenger | Tetra Hunt |
| --- | --- | --- | --- | --- | --- | --- | --- | --- | --- |
| 2.53E-03 | 5.58E-03 | 6.51E-01 | 2.17E-07 | 6.44E-03 | 7.57E-02 | 3.99E-03 | 2.12E-01 | 4.28E-02 | 4.41E-04 |

## Discussion

### Body Mass

Bohaiornithid masses are most consistent with folivory, frugivory, generalist feeding, piscivory, scavenging, or tetrapod hunting. Mean estimates of mass for most bohaiornithid specimens (Table 1) are near the 265 g split between frugivores + generalists + piscivores + scavengers (group c) and folivores + tetrapod hunters (group d) (Fig. 3), and the uncertainties of both the split and the masses cause all but two bohaiornithid specimens to fall within both groups c and d. The exceptions are the indefinite bohaiornithid CUGB P1202, firmly within group c but also highly immature, and *Zhouornis* CNUVB-903 whose estimated mass is firmly within group d. The largest specimens of *Bohaiornis*, *Shenqiornis*, *Sulcavis*, and *Zhouornis* have mean mass estimates which are greater than 265 g, which we interpret as stronger affinity with folivory and tetrapod hunting than other bohaiornithids.

We note that every bohaiornithid is less massive than any avian obligate scavenger, and facultative scavenging in birds as small as even the largest bohaiornithid, *Zhouornis* CNUVB- 903, is facilitated by anthropogenic waste [47, 48] and thus may not represent a natural state.

Coupled with past work proposing very large body size is a prerequisite for obligate scavenging [49], we consider scavenging less likely in bohaiornithids than the other diets mentioned above.

### Mechanical Advantage and Functional Indices

As previously observed [8, 9], skull functional indices only effectively separate folivores, husking granivores, and scavengers in the functional morphospace (Fig. 4). This means dietary resolution is poor from this line of evidence. In addition to previously-proposed explanations of MA adaptations being only impactful in particular lineages [23] or ecologies [8], we note here the need to investigate how cranial kinesis in extant birds affects measures of their skull function. Extension of the premaxilla can increase outlever distance and modify both upper jaw anterior jaw-closing mechanical advantage (AMA) and jaw-opening mechanical advantage (OMA), though the extent of excursion of the premaxilla in life and whether it occurs during biting is unrecorded for the vast majority of taxa. Future investigation into this unknown and incorporation of the data into MA analyses may yield better explanatory power for ecological aspects.

**Figure 4.**
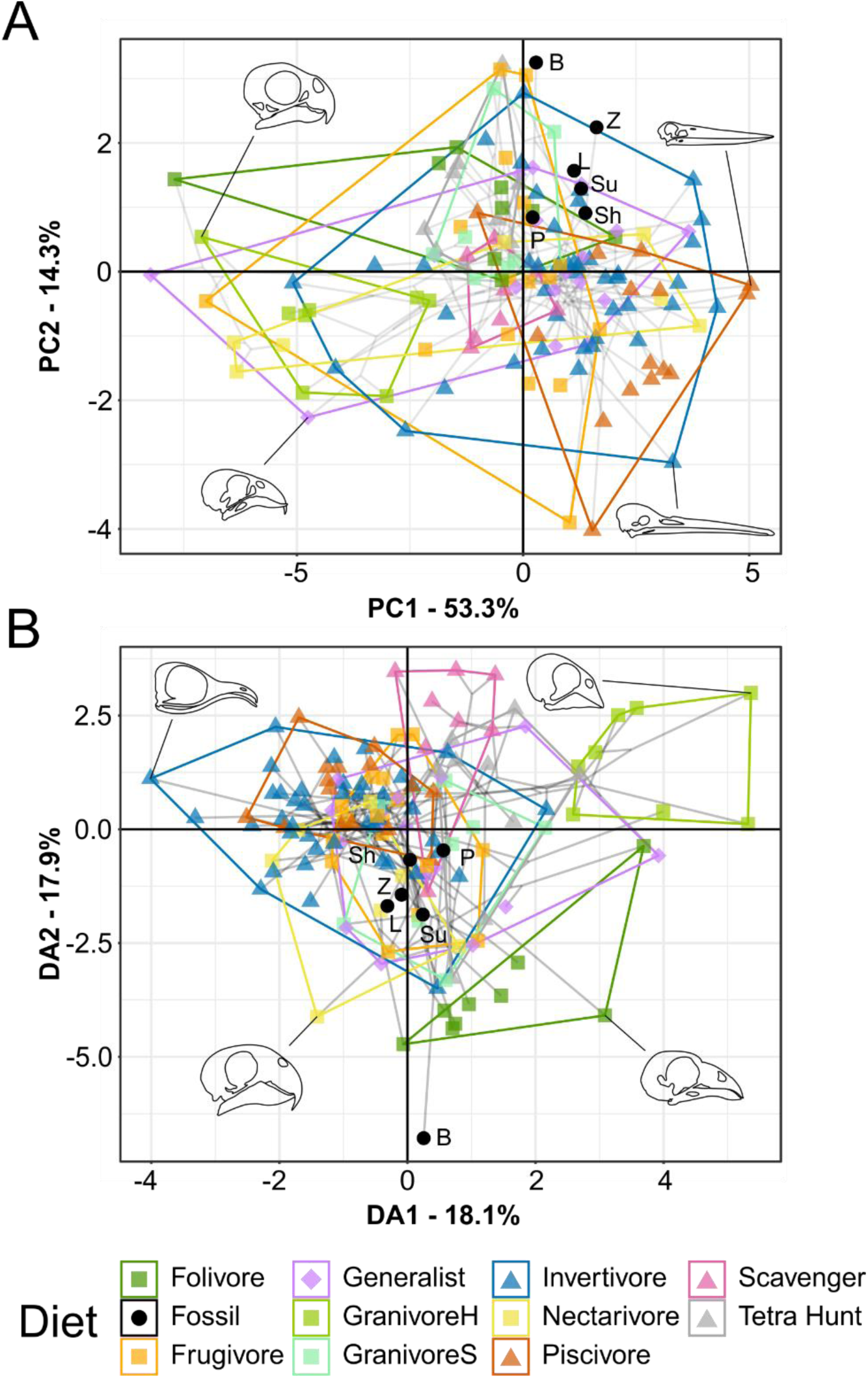
Functional phylomorphospace of MA and functional index data, grouped by diet. Grey lines indicate phylogenetic relationships. Line drawings of skulls for selected taxa are provided for reference. Data are presented with PCA (A) and FDA (B). See Fig. S2 for character weights and Table 2 for FDA posterior predictions. Diet abbreviations: GranivoreH, Husking Granivore; GranivoreS, Swallowing Granivore; Tetra Hunt, Tetrapod Hunter. Fossil taxon abbreviations: B, *Bohaiornis*; L, *Longusunguis*; P, *Parabohaiornis*; Sh, *Shenqiornis*; Su, *Sulcavis*; Z, *Zhouornis*.

#### Species-level interpretations

Bohaiornithids separate into two distinct functional guilds: *Bohaiornis* and *Parabohaiornis* in one with distinctly high OMA; and *Longusunguis*, *Shenqiornis*, *Sulcavis*, and *Zhouornis* with OMA values more in line with the average extant bird. The posteroventral of the cranium is a landmark location for OMA and this region is reconstructed in every other taxon with material from *Bohaiornis* specimen IVPP V17963 (Fig. 2), so it is possible that the OMA is overestimated in other bohaiornithid taxa. Given the similarity of the posterior deflection of the posterodorsal cranium in *Shenqiornis*, *Sulcavis*, and especially *Parabohaiornis* to the deflection seen in *Bohaiornis* we consider OMA values for non-*Bohaiornis* bohaiornithids to be reasonable estimates.

*Bohaiornis* has unusually high OMA (0.16 lower and 0.33 upper; Fig. S1E–F). Folivores have a diagnostically high OMA (x̄ = 0.10 lower, 0.27 upper; Fig. S1E–F), making this the strongest affinity for this taxon (Table 2). Notably, though, it has a posterior jaw-closing mechanical advantage (PMA) below any extant folivore (0.16 lower jaw PMA and 0.09 upper jaw PMA, vs a folivore minimum of 0.24 lower jaw PMA and 0.18 upper jaw PMA; Fig. S1C–D). Note that OMA is strongly affected by the length of the cranium, which may be under different developmental constraints in bohaiornithids (see Skull Traditional Morphometrics Discussion). However, if anything one would expect bohaiornithids to have shorter crania and thus lower OMA than an equivalent crown bird, making a distinctly high OMA even more impactful. *Parabohaiornis* has considerable affinity with swallowing granivores and folivores (Table 2), respectfully due to its relatively high AMA and PMA and lower jaw OMA (Fig. S1A– F).

The remaining bohaiornithids plot within the region of undifferentiated diets (Fig. 4) and have the strongest affinity with dietary generalists (Table 2), previously interpreted as a “default” prediction in longipterygids [8]. These taxa, however, do show greater affinity with frugivores and less with invertivores and piscivores compared to pengornithids (though *Shenqiornis* has nearly the same affinity with piscivores), bringing all three diets into roughly equal likelihood after generalist feeding. The exception is *Sulcavis*, whose slightly higher OMA increases affinity with folivores rather than frugivores. The posterior skull of *Shenqiornis* is largely reconstructed from cf. *Sulcavis* material (Fig. 2D), so its membership in this functional guild is less certain.

#### Evaluating bohaiornithid durophagy

Durophagy, as hypothesized in bohaiornithids [14, 15, 16 pg. 83, 17], is poorly-defined. Thus we cannot clearly define a “durophage” diet category to test this hypothesis. Rather, we can test it indirectly by comparing bohaiornithid jaw performance to that of husking granivores and parrots. Husking granivores are labelled as such in this work based on regular recorded behaviour of cracking hard seeds (Table S2), and parrots are the archetypical avian durophages with adaptations across the clade for withstanding high bite forces [50]. The jaw-closing mechanical advantage of *Parabohaiornis* is higher than the nectarivorous brown lory (*Chalcopsitta duivenbodei*; upper jaw AMA 0.12 in lory vs 0.15 in *Parabohaiornis*), but smaller than every other parrot and husking granivore. The average for upper jaw AMA for parrots (0.23) and husking granivores (0.24) is much higher. The maximum relative jaw height in bohaiornithids is also below the minimum in parrots, and the same is true for husking granivores in the lower jaw. The average relative maximum cranial height (MCH) of husking granivores and bohaiornithids is nearly the same (respectively 0.35 and 0.34). In short, the upper jaw in bohaiornithids is somewhat robust, but the upper and lower jaws lack force production adaptations seen in extant avian durophages.

### Finite Element Analysis

Bohaiornithid jaws are stronger than both longipterygids’ and pengornithids’. Under the same construction and loading conditions, longipterygid jaws ranged from 259 to 354 µε [8] and pengornithid jaws from 190 to 275 µε [9]. Bohaiornithids, in contrast, have jaw strain ranging from 89 to 156 µε (Fig. 5). While they do separate from these other enantiornithine families, the intermediate strength of their jaws compared to extant birds means they overlap with most diets in multivariate space (Fig. 6). Thus, dietary diagnosis from FEA is less precise in bohaiornithids than previously-studied enantiornithines.

**Figure 5.**
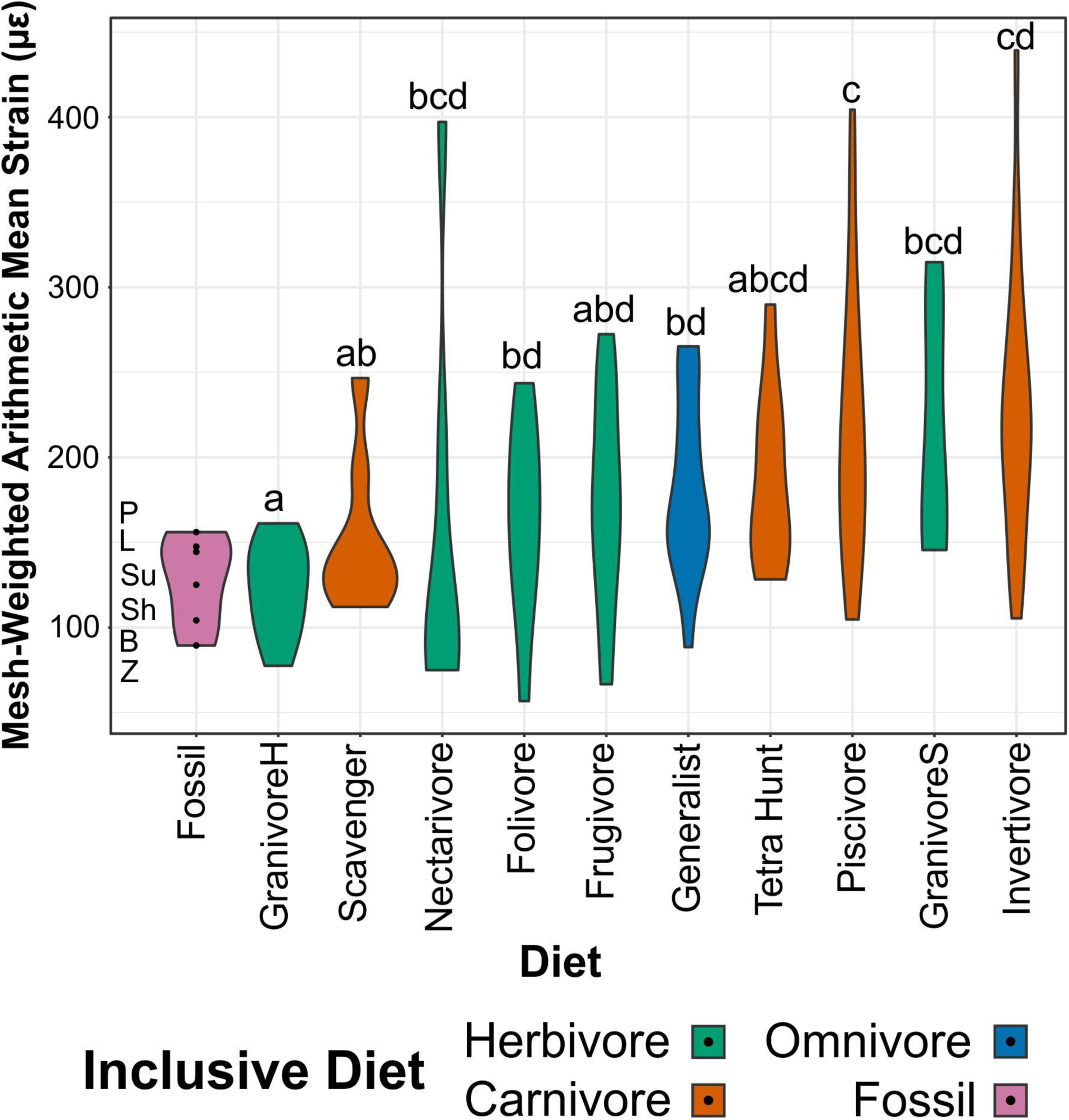
Violin plots of MWAM strain of FEA models, organized by diet. Extant diets ascend in average MWAM strain from left to right. MWAM strains were tested for significant differences via phylogenetic HSD. Diets marked with the same letter are not significantly different from one another. Diet abbreviations: GranivoreH, Husking Granivore; GranivoreS, Swallowing Granivore; Tetra Hunt, Tetrapod Hunter. Fossil taxon abbreviations: B, *Bohaiornis*; L, *Longusunguis*; P, *Parabohaiornis*; Sh, *Shenqiornis*; Su, *Sulcavis*; Z, *Zhouornis*.

**Figure 6.**
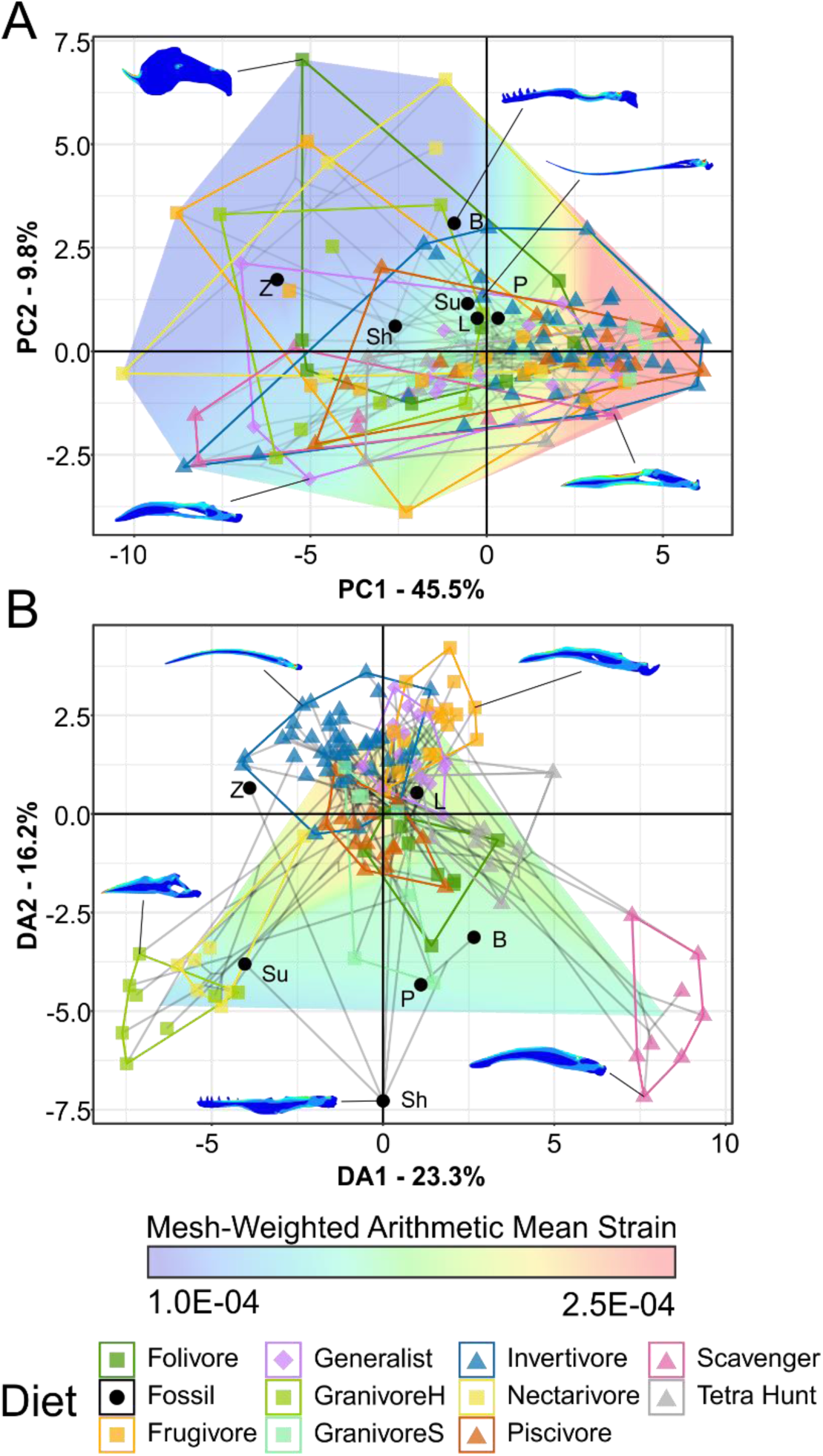
Phylogenetic strain-space of total maximum in-plane principal strain intervals for extant and fossil bird lower jaw finite element models, grouped by diet. MWAM strain is mapped overtop the data. Grey lines indicate phylogenetic relationships. Contour plots for selected taxa are provided for reference. Data are presented with PCA (A) and FDA (B). In PCA (A) overall strain increases along PC1, and strain heterogeneity (i.e. lower areas of intermediate strain) increases along PC2. In FDA (B), DA1 and DA2 have loadings of various similar low-strain intervals, with high-strain intervals clustering near the origin. See Table 3 for FDA posterior predictions. Diet abbreviations: GranivoreH, Husking Granivore; GranivoreS, Swallowing Granivore; Tetra Hunt, Tetrapod Hunter. Fossil taxon abbreviations: B, *Bohaiornis*; L, *Longusunguis*; P, *Parabohaiornis*; Sh, *Shenqiornis*; Su, *Sulcavis*; Z, *Zhouornis*.

#### Species-level interpretations

In lieu of a clear separation in multivariate space, we can still consider diets unlikely for a taxon if the mesh-weighted arithmetic mean (MWAM) strain of the jaw under loading falls outside the range of any birds with that diet. However, unlike in Longipterygidae in which the fossil taxa’s jaws are weaker than many extant diet groups [8], bohaiornithid jaws are stronger than most extant birds’. Rather than being unable to process a given food as in a weaker jaw, we interpret stronger jaws as “overbuilt” for a given diet. This implies the jaw evolved under pressures for more durable food, but does not necessarily prohibit consumption of more compliant foods.

*Bohaiornis*, *Shenqiornis*, *Sulcavis*, and *Zhouornis* experience less strain than any swallowing granivore in the extant dataset; *Bohaiornis*, *Shenqiornis*, and *Zhouornis* also experience less than any tetrapod hunter; and *Bohaiornis* and *Zhouornis* experience less than any carnivore (Fig. 5).

In multivariate space (Fig. 6A) it is apparent that bohaiornithid jaws tend to have a heterogeneous strain distribution. This contrasts with both scavengers and tetrapod hunters, which have a diagnostically homogenous strain distribution. To a lesser extent frugivores also tend to have homogenous strain, though the frugivorous African green pigeon *Treron calva* is notably the nearest neighbour of *Zhouornis*. *Shenqiornis* and *Sulcavis* notably do not plot near one another despite the angular and much of the surangular in the *Shenqiornis* reconstruction coming from cf. *Sulcavis* (Fig. 2D), owing largely to the more robust posterior dentary in *Shenqiornis*.

#### Evaluating bohaiornithid durophagy

Comparison to husking granivores and parrots warrants additional attention to evaluate hypotheses of durophagy. Bohaiornithids have overall low jaw strain under loading (89–156 µε, x̄ = 128 µε). This is on par with husking granivores (78–161 µε, x̄ = 123 µε), but weaker than parrots (57–95 µε, x̄ = 78 µε). Husking granivores plot across PC2 in Fig. 6, but parrots and bohaiornithids both experience mainly heterogeneous strain patterns. The locations of high and low strains differ between the two groups, though. Parrots tend to experience high strain at the rostral tip of the jaw, while the posterior end of the jaw is under minimal strain. Conversely bohaiornithids tend to experience high strains along the dorsal edge of the jaw, particularly near the dentary/surangular suture, and low strains along the ventral edge of the jaw. This may be partially explained by the lack of a keratinous beak in bohaiornithids, which acts as a strain sink in the rostral jaw of beaked animals [51]. Bohaiornithids have jaws of similar overall strength to husking granivores but weaker than parrots, and the patterns of strain they experience during a bite are not analogous to either group.

### Traditional Morphometrics

#### Pes

##### Evaluating bohaiornithid raptorial behaviour

As previously proposed [12], most bohaiornithid talons plot near some extant raptorial birds (Fig. 7) and have some affinity with them in flexible discriminant analysis (FDA) and phylogenetic FDA (pFDA; Table 4). However, most taxa also have some level of non-raptorial affinity.

**Figure 7.**
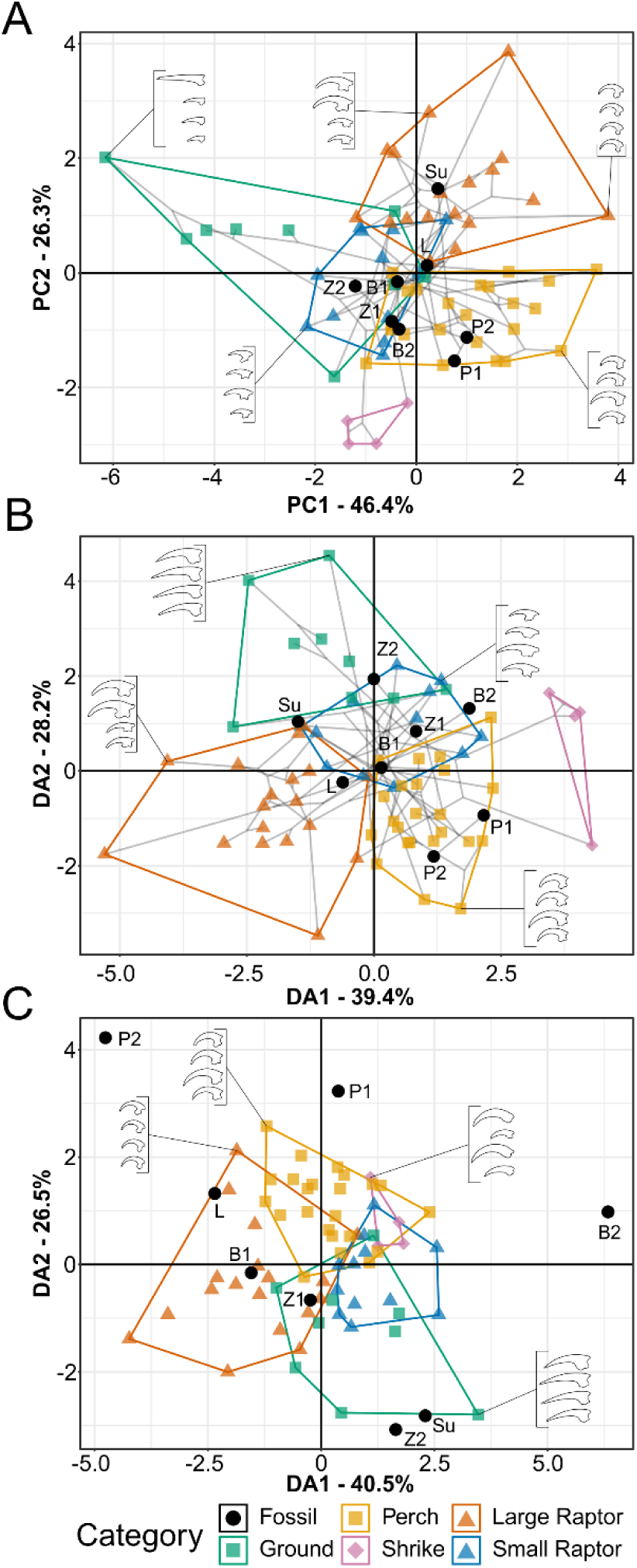
Phylomorphospace of extant and fossil bird claw shape from pedal TM, grouped by pedal ecological category. Grey lines indicate phylogenetic relationships. Line drawings of claws for selected taxa are provided for reference. Data are presented with PCA (A), FDA (B), and pFDA (C). See Fig. l for character weights and Table 4 for FDA and pFDA posterior predictions. Category abbreviations: Large Raptor, raptor taking prey which does not fit in the foot; Small Raptor, raptor taking prey which can fit in the foot. Fossil taxon abbreviations: B1, *Bohaiornis* LPM B00167; B2, *Bohaiornis* IVPP V17963; L, *Longusunguis* IVPP V18693; P1, *Parabohaiornis* IVPP V18690; P2, *Parabohaiornis* IVPP V18691; Su, *Sulcavis* BMNH Ph-805; Z1, *Zhouornis* CNUVB-0903, Z2, *Zhouornis* BMNHC Ph 756.

##### Species-level interpretations

*Longusunguis* is the bohaiornithid most consistently recovered here as raptorial. It plots within or near large raptors in every analysis (Fig. 7) and is consistently recovered as likely to be a large raptor (Table 4). The joints of digits I and II are somewhat hinged (= ginglymoid *sensu* [52]), though digits III and IV are unhinged [53] indicating grasping ability focussed in the first two digits as in many extant accipiters [36]. As noted by [12] the tarsometatarsus of *Longusunguis* and other bohaiornithids is relatively short and robust, typically interpreted as increasing grip strength at the cost of speed [36, 39, 54] and more common among raptorial avians in ambush predators [39].

*Sulcavis* has the strongest raptorial affinity in principle component analysis (PCA; Fig. 7A) and has some raptorial affinity in FDA and pFDA (Fig. 7B; Table 4). Unexpected for a raptorial bird, the specimen displays a relatively small and straight digit II ungual. Conversely, digit I is the most enlarged and recurved of any bohaiornithid and among the most enlarged and recurved in the dataset overall. Enlargement of the hallux without enlargement of any opposing digit is uncommon in extant birds, mostly occurring in larks (Alaudidae) in which the digit I is also nearly straight. Digit II is damaged in the holotype of *Sulcavis* [15], and it may be that our estimate of its extent does not reflect its actual enlargement and curvature. If this is the case then it likely used its feet raptorially as we hypothesise for *Longusunguis*, but if not then this may indicate some unique use for digit I other than raptorial grasping.

*Zhouornis* and *Bohaiornis* can reasonably be interpreted either as small raptors or non-raptorial perchers. Both have strong affinity with small raptors in FDA (Fig. 7B, Table 4) but also plot near ground birds and non-raptorial birds in PCA (Fig. 7A). Their claw curvature (average 91°) is dissimilar from ground birds, rendering that diagnosis less likely. Their phalanges are moderately hinged [18, 55–57] which indicates some level of grasping adaptation, useful both for raptors and non-raptorial perchers. The more mature specimen of each bird (B2 and Z1 in Fig. 7A; Fig. S10) has claws less similar to raptors and more similar to non-raptorial perchers. Digits I, II, and IV maintain similar proportions across ontogeny within each species, but in both species digit III increases in relative size in the more mature specimens (less confident in *Bohaiornis* as pes preservation is poor in IVPP V17963 [18]), explaining the shift towards non-raptorial birds.

*Parabohaiornis* consistently resolves as a non-raptorial perching bird across all analyses (Fig. 7, Table 4). The taxon has a relatively small digit I compared to other bohaiornithids, driving this finding. Published image resolution is too poor to tell how hinged their phalanges are [12].

#### Skull

Skull traditional morphometrics, while useful in determining longipterygid diet [10], appear less useful when examining bohaiornithids. Bohaiornithids have rostra to skull length ratios clustered around 0.8, a region inhabited by birds of all diets except nectarivores and granivores. Bohaiornithids also do not conform to the general correlation of increasing rostrum length and skull length. If anything, those with longer skulls have shorter rostra.

We offer two possible interpretations. Firstly, some bohaiornithids may have skull functions not discussed here. *Bohaiornis*, *Longusunguis*, and *Parabohaiornis*, do plot at the edge of the extant space near folivores (*Opisthocomus hoazin* and two phasianids), frugivores (two psittacoids), and a generalist (*Numida meleagris*). *Shenqiornis*, *Sulcavis*, and *Zhouornis*, however, plot outside the extant space and may represent an ecology dissimilar from any sampled extant bird.

Alternatively, this may indicate different developmental constraints on the skull in bohaiornithids (and possibly enantiornithines as a whole). One hypothesis proposes that crown birds ancestrally have relatively larger brains than their earlier-diverging counterparts [58], which may necessitate relatively longer skulls to house the larger brain. However, a subsequently-discovered enantiornithine basicranium displays many crown-bird-like characteristics and has been used to argue a crown-bird-like endocranium was present in the common ancestor of Ornithothoraces [59]. Given this uncertainty, we use skull TM only tentatively to interpret ecology in bohaiornithid taxa and generally recommend caution in interpreting non-crown bird ecology from measures including the cranium.

### Bohaiornithid Ecology and Evolution

By combining the individual lines of evidence discussed above, more precise diet diagnoses can be obtained. Table 7 summarizes these diagnoses.

**Table 7.** Summary table of interpretations of each line of evidence used herein. Body mass, MA, and FEA inform diet. Pedal TM informs the use of the pes or lack thereof in feeding. See relevant Discussion sections for additional details Bolded diets are agreed upon by all available diet proxies. Bolded pedal ecologies are either help discriminate diet (carnivory in *Longusunguis*) or are supported over other possibilities by diet information (non-raptorial perching in *Bohaiornis*). Given the uncertain application of skull TM to bohaiornithids, diets from this proxy are not bolded.

| Line of Evidence | Taxon | Likely Diets/Ecologies | Unlikely Diets/Ecologies |
| --- | --- | --- | --- |
| Body Mass | *Beiguornis* | Folivore, Frugivore, Generalist, Piscivore, Tetrapod Hunter | Granivore, Invertivore, Nectarivore |
|  | *Bohaiornis* | **Folivore**, Frugivore, Generalist, Piscivore, Tetrapod Hunter | Granivore, Invertivore, Nectarivore |
|  | *Longusunguis* | Folivore, Frugivore, **Generalist**, **Piscivore**, Tetrapod Hunter | Granivore, Invertivore, Nectarivore |
|  | *Parabohaiornis* | **Folivore**, Frugivore, **Generalist**, Piscivore, Tetrapod Hunter | Granivore, Invertivore, Nectarivore |
|  | *Shenqiornis* | **Folivore**, Frugivore, **Generalist**, **Piscivore**, Tetrapod Hunter | Granivore, Invertivore, Nectarivore |
|  | *Sulcavis* | **Folivore**, Frugivore, **Generalist**, **Piscivore**, Tetrapod Hunter | Granivore, Invertivore, Nectarivore |
|  | *Zhouornis* | Folivore, **Frugivore**, **Generalist**, Piscivore, Scavenger, Tetrapod Hunter | Granivore, Invertivore, Nectarivore |
| Mechanical Advantage | *Bohaiornis* | **Folivore**, Generalist, Swallowing Granivore | Husking Granivore, Scavenger |
|  | *Longusunguis* | Frugivore, **Generalist**, Invertivore, **Piscivore** | Husking Granivore, Scavenger |
|  | *Parabohaiornis* | **Folivore, Generalist**, Scavenger, Swallowing Granivore | Husking Granivore, Tetrapod Hunter |
|  | *Shenqiornis* | **Folivore**, Frugivore, **Generalist**, Invertivore, **Piscivore** | Husking Granivore, Tetrapod Hunter |
|  | *Sulcavis* | **Folivore, Generalist**, Invertivore, **Piscivore** | Husking Granivore, Scavenger |
|  | *Zhouornis* | **Frugivore, Generalist**, Invertivore, Piscivore | Husking Granivore, Scavenger |
| Finite Element Analysis | *Bohaiornis* | **Folivore**, Husking Granivore, Nectarivore | Swallowing Granivore, Invertivore, Piscivore, Scavenger, Tetrapod Hunter |
|  | *Longusunguis* | Folivore, Frugivore, **Generalist**, Husking Granivore, Swallowing Granivore, Invertivore, Nectarivore, **Piscivore** | Scavenger, Tetrapod Hunter |
|  | *Parabohaiornis* | **Folivore**, Frugivore, **Generalist**, Husking Granivore, Swallowing Granivore, Invertivore, Nectarivore, Piscivore | Scavenger, Tetrapod Hunter |
|  | *Shenqiornis* | **Folivore**, Frugivore, **Generalist**, Husking Granivore, Invertivore, Nectarivore, **Piscivore** | Swallowing Granivore, Scavenger, Tetrapod Hunter |
|  | *Sulcavis* | **Folivore**, Frugivore, **Generalist**, Husking Granivore, Invertivore, Nectarivore, **Piscivore** | Swallowing Granivore, Scavenger, Tetrapod Hunter |
|  | *Zhouornis* | **Frugivore, Generalist**, Husking Granivore, Nectarivore | Swallowing Granivore, Invertivore, Piscivore, Scavenger, Tetrapod Hunter |
| Pedal Traditional Morphometrics | *Bohaiornis* | **Non-Raptorial Perching**, Small Raptor | Ground Bird, Shrike-like |
|  | *Longusunguis* | **Large Raptor** | Ground Bird, Shrike-like |
|  | *Parabohaiornis* | Non-Raptorial Perching | Ground Bird |
|  | *Sulcavis* | Large Raptor, unique ecology not in dataset | Shrike-like |
|  | *Zhouornis* | Small Raptor, Non-Raptorial Perching | Ground Bird, Shrike-like |
| Skull Traditional Morphometrics | *Bohaiornis* | Folivore, Frugivore, Generalist | Granivore, Nectarivore |
|  | *Longusunguis* | Folivore, Frugivore, Generalist | Granivore, Nectarivore |
|  | *Parabohaiornis* | Folivore, Frugivore, Generalist | Granivore, Nectarivore |
|  | *Shenqiornis* | Unique ecology not in dataset | All sampled diets |
|  | *Sulcavis* | Unique ecology not in dataset | All sampled diets |
|  | *Zhouornis* | Unique ecology not in dataset | All sampled diets |

#### Species-level diet diagnoses

*Bohaiornis* has jaws highly reminiscent of avian folivores (note we use this term to refer to animals primarily consuming any non-reproductive plant tissues, not strictly leaves; Fig. 1 centre), its relative skull and rostrum size is highly similar to them, and its body mass that falls within the folivore range as well. As noted in the MA discussion, the most uniquely folivorous trait of this taxon is a high OMA. While the PMA of *Bohaiornis* is below that of any studied folivore, its jaw strength in FEA is also similar to that of folivores. The other groups it plots near in FEA function space (Fig. 6A), husking granivores and nectarivores, tend to be much less massive than *Bohaiornis*. An herbivorous diet would imply the claws were not used to kill prey, which is plausible for *Bohaiornis* (Fig. 7). Perching/arboreal lifestyles for living folivorous birds are uncommon, and these birds are typically weak fliers [60]. The hoatzin (*Opisthocomus hoazin*) and southern screamer (*Chauna torquata*) have both similar OMA (respectively 0.37 upper 0.07 lower and 0.28 upper 0.13 lower) and MWAM strain (both 121 µε) to *Bohaiornis* (upper OMA 0.33, lower OMA 0.16, MWAM strain 104 µε). Both of these extant folivores typically fly only short distances with continuous flapping [61, 62] which is also the flight style predicted for bohaiornithids [19]. These two taxa serve as the most likely extant analogues for *Bohaiornis*; either birds which spent most of their lives feeding and climbing in trees like the hoatzin [61] or more ground-based foragers who use trees as a refuge when resting or threatened [62]. *Bohaiornis* IVPP 17963 plots near the hoatzin in pedal TM (Fig. 7A), so the former lifestyle seems more likely for at least a mature *Bohaiornis*.

*Parabohaiornis* has strong herbivorous affinities, though they are less clear than those in *Bohaiornis*. Its skull is most similar to swallowing granivores in both MA (Table 2) and FEA (Table 3), though it is more massive than most extant granivorous birds (178–406 g, vs upper cutoff of 53 g). Sandgrouse (Pteroclidae) may serve as an extant analogy as swallowing granivores in the 200–300 g mass range [63]. The two sandgrouse in this study, *Pterocles exustus* and *Pterocles orientalis*, have overall similar MA (e.g. upper jaw AMA 0.18 and 0.19, 0.15 in *Parabohaiornis*) and skull robusticity (ACH 0.24 and 0.23, 0.25 in *Parabohaiornis*), but their jaws are weaker under loading in FEA (MWAM strain 311 and 273 µε, 156 µε in *Parabohaiornis*). Also of note, sandgrouse are generally terrestrial while the unguals of *Parabohaiornis* indicate an arboreal lifestyle. Alternatively, *Parabohaiornis* shows the next most affinity with folivores, and this diagnosis is more consistent with its large body mass and relatively short skull and intermediate-length rostrum. Like *Bohaiornis*, *Parabohaiornis* is adapted for non-raptorial perching and would likely be analogous to a hoatzin if folivorous. Generalist feeding is recovered as possible for Parabohaiornis in MA and FEA, but consistently less so than granivory and folivory so it is treated here as a “default prediction” [8] and not discussed at length.

The crop and gizzard (i.e. ventriculus or gastric mill) play major roles in folivory and granivory for extant avians, serving as a site to process the tough plant matter during digestion [60, 64]. Crops and gizzards have previously been inferred to be absent in Enantiornithes [4], which could potentially be strong evidence against folivory or granivory in *Bohaiornis* and *Parabohaiornis*. Two (non-exclusive) possibilities may still allow for effective herbivory in these taxa: oral processing and hindgut specialisation. Most groups of herbivorous lizards have minimal postcranial digestive specialisation, and instead are able to orally process plants to a level where nutrients can be efficiently extracted [65]. The lack of any accessory cusps as seen in many herbivorous lizard clades [66] is unsurprising in bohaiornithids, given the simplification of enamel at the Avialae node [67]. Better analogues are skinks (Scincidae) and tegus (Teiidae), which retain conical dentition which is simply more robust than their carnivorous and omnivorous relatives [66]. Bohaiornithids, notably, have some of the most robust teeth within Enantiornithes [14]. This may have been an adaptation or exaptation for processing plant matter. Additionally, several groups of extant folivorous birds have specialised hindguts to process plant matter. Both emus [68] and geese [69] have regions of their digestive tract specialized in fermenting fibrous foods, and anseriform [70, 71] and galliform [72–74] birds are known to increase their intestinal lengths when consuming high fibre diets, which allows for a higher percentage to be digested. While we lack any direct evidence of these adaptations in Bohaiornithidae, we contest there is evidence for specialised hindgut digestion of plant material in enantiornithines more broadly. Structures now recognised as plant propagules [75] have been identified in seven enantiornithine specimens [76]. Of these, four specimens are figured (STM 10-45, STM 29-8 [77], STM 10-12 [76], and STM 11-80 [78]), and in these figured specimens the propagules are preserved either in the lowest thoracic (STM 10-45 and STM 11-80) or pelvic (STM 29-8 and STM 10-12) region. As previously noted, consumulite preservation is most likely with long gut retention times [2], so preservation of these plant propagules specifically in the region inhabited by the hindgut in life implies a prolonged retention in this area where plant matter could ferment and break down. So, despite likely lacking a crop and gizzard, *Bohaiornis* and *Parabohaiornis* could have sustained folivory with oral processing of plant matter and/or specialised fermentation in the hindgut.

Alternatively, as noted in the FEA discussion, our diagnosis of *Bohaiornis* and *Parabohaiornis* as herbivorous is based on the relatively high strength and efficiency of their jaws. These adaptations indicate evolutionary pressure for the ability to consume plant material, but not necessarily a regular habit of doing so. “Fallback foods”, foods avoided during normal feeding but essential to survival when alternatives are unobtainable, are well-documented evolutionary drivers of extant animals’ functional morphology [79]. The Jehol climate is believed to have been highly variable and affected by local volcanic activity [80], so specialised herbivory may have been restricted to times when this instability limited other food resources. If this is the case, then specialized oral or hindgut processing may not have been necessary to allow *Bohaiornis* and *Parabohaiornis* to survive short-term resource scarcity.

*Zhouornis* has the strongest jaw among bohaiornithids (Fig. 5), with total evidence pointing to either a frugivorous or generalist lifestyle. Only frugivores, generalists, husking granivores, and nectarivores inhabit a region of the FEA strain space where jaw strain is as low and heterogeneous as in *Zhouornis*. This space is generally sparsely occupied and thus seems ecologically meaningful. However, *Zhouornis* is also the largest bohaiornithid (mean estimate 758 g for the largest specimen), and very few extant granivorous or nectarivorous birds reach this size (Fig. 3). Frugivory and generalist feeding are also recovered as likely from MA posterior predictions (Table 2), so these two diets are considered most likely for *Zhouornis*. Pedal TM results could support either diet, with talons similar to both raptors taking small prey (potentially useful for a generalist) and non-raptorial perching birds (as expected of a frugivore). If *Zhouornis* was indeed a generalist it would potentially have an interesting niche overlap with that proposed for *Pengornis* [9]. *Pengornis* shows signs of generalist feeding with ability to raptorially take large prey. While *Zhouornis*’ pes more resembles that of raptors taking prey that is small relative to body size, *Zhouornis*’ absolute body size is much larger than *Pengornis* (mean mass estimate 437 g [26]). This may represent a different approach to taking advantage of the same prey resources. We predict *Zhouornis* would be capable of macrocarnivory but less suited for it than *Pengornis*; the barred owl (*Strix varia*) has a similar body mass and claw shape to *Zhouornis*, and while it is capable of taking large prey like grouse and rabbits it preferentially takes mouse-sized prey [81].

*Longusunguis*, *Shenqiornis*, and *Sulcavis* have less clear diagnoses. Piscivory and generalist feeding are recovered as likely for all three, with folivory also likely for *Shenqiornis* and *Sulcavis*. Unlike *Parabohaiornis*, however, there is no clear hierarchy of these possibilities. Much of the uncertainty stems from FEA giving little useful data, with these taxa having intermediate MWAM strains when loaded (Fig. 5) and inhabiting a region of the strain-space shared by most diets (Fig. 6A) only ruling out scavenging and tetrapod hunting. As mentioned above, their intermediate mass can also only rule out granivory, invertivory, and nectarivory. We are therefore left with MA to discriminate between the remaining diets, which has poor predictive power in extant birds. Generalist feeding and piscivory are recovered as somewhat likely for all three from MA (Table 2), though as previously proposed this could represent a default prediction [8]. Folivory can also not be ruled out in any of these taxa due to uncertainty in quadrate placement (Table S3). Pedal TM, however, points to folivory being unlikely in *Longusunguis* as its talons appear adapted for taking large animal prey. *Longusunguis* notably has more recurved teeth than other bohaiornithids [figures 4 and 6 in 17] which is associated with carnivory [82]. Skull TM of *Longusunguis* are most similar to folivorous birds, but given the uncertainty in what developmental constraints are at play we consider folivory overall less likely in *Longusunguis*. Claw shape may also point to carnivory in *Sulcavis*, though as discussed above its raptorial affinities are questionable. *Shenqiornis* only preserves two disarticulated talons [83], and while these superficially resemble the claws of *Zhouornis* in size and curvature we cannot quantitatively reject hypotheses of the pes not being adapted for prey capture, and in turn hypotheses of folivory, in *Shenqiornis* or *Sulcavis* (*contra* our previous carnivorous assignment of *Shenqiornis* [2]). Diagnosis of *Shenqiornis* is further complicated by the reliance on cf. *Sulcavis* material to reconstruct its skull (Fig. 2D), artificially inflating the functional similarity of the two genera. Discovery of additional *Shenqiornis* specimens should clarify the accuracy of this model. In any case, even with four lines of evidence the dietary diagnosis of these three taxa remains tentative, and investigating additional lines of evidence (e.g. dental microwear, muscle reconstruction, or stable isotope analysis [2]) is necessary.

*Beiguornis* preserves neither a skull nor talons [22], so only mass data is available. As with most bohaiornithids, the only diets which can be considered unlikely based on its mass are invertivory, granivory, and nectarivory. Further ecological information will require new techniques or more complete specimens.

#### Evaluating bohaiornithid ecological hypotheses

Our results support hypotheses of raptorial behaviour [2, 12, 16 pg. 270, 18] in bohaiornithids, though not for every taxon. We interpret *Longusunguis* as a small predator adapted to take prey too large to be completely encircled by the pes, and *Zhouornis* as potentially a large predator whose talons were adapted to take prey which could be fully encircled by the pes. *Sulcavis* has a greatly enlarged and recurved digit I as expected in a raptor, but lacks any enlarged opposed digit that could be used to grip. As noted above, this could be taphonomy obscuring the full extent of digit II or a different, unaccounted for use of the pes. The talons of *Parabohaiornis* are most akin to non-raptorial perching birds, and while the talons of *Bohaiornis* are between those of raptors and non-raptors we interpret them as non-raptorial given the taxon’s herbivorous skull adaptations. Note that the alleged rangle (raptorial gastroliths) used as previous evidence of raptorial behaviour in *Bohaiornis* [18] has been reinterpreted as post-mortem mineral growth [84]. Specimens of *Shenqiornis* preserving talons are needed to opine on whether it was adapted for raptorial behaviour. We previously interpreted *Shenqiornis* as carnivorous from a combination of its MA and prior interpretations of all other bohaiornithids as raptorial [2], but with the variety of pedal ecologies identified in this study we now consider its diet indeterminate.

We also support hypotheses of durophagy [14, 15, 16 pg. 83, 17] in bohaiornithids, though with the caveat that their adaptations are less extreme than what one usually considers an avian durophage. The first proposal of durophagy in a bohaiornithid is made in contrast to proposing most other enantiornithines as “general insectivore[s], limited to small arthropods with softer cuticles” [14 pg. 153]. We support Bohaiornithids as durophagous in the sense that they have higher jaw strengths than other modelled enantiornithines (see above). They appear adapted for taking foods harder and tougher than soft-bodied arthropods, but the same could be said for pengornithids. In living birds, “durophagy” typically refers to cracking hard foods like seeds or nuts within the beak [85]. As examined in the MA and FEA discussion sections, bohaiornithids resemble extant husking granivores in both the relative height of their upper jaw and the overall strain under loading of their lower jaw; however, they lack the adaptations for bite force production of husking granivores, and fall short of parrots in all potential durophagy metrics. Bohaiornithids, therefore, do not display adaptations for taking foods as hard as extant birds commonly considered durophages.

Based on our findings herein, we encourage caution in future works using hypothesized diet as a variable. Notably when [17] investigates avialan tooth shape in relation to diet, bohaiornithids are placed into the single “durophage” diet category and are its only members. Bohaiornithid tooth shape does tightly cluster in the study’s phylomorphospaces [figures 4–6 in 17], but they notably tend to take a central position in the morphospace and overlap with purported granivores, insectivores, and piscivores. Given this overlap and the ecological diversity within the group we present here, we suggest the similarity in bohaiornithid tooth shape is driven by phylogeny rather than, as proposed by [17], diet. The robust teeth in Bohaiornithidae may serve as general-use tools capable of processing diverse foods effectively rather than adaptations to a specialised diet shared across the clade.

### Evolutionary History of Enantiornithine Ecology

#### Trends within the three major enantiornithine families

Our data suggest that bohaiornithids cover a greater ecological breadth than previously-studied enantiornithine families (Fig. 9). Longipterygids are reconstructed as mainly invertivorous with variability in foraging environment [8] (Fig. 1 left; though our mass results here suggest the hypothesis of *Longipteryx* as a piscivore cannot be rejected, *contra* [8], see supplemental Discussion), and pengornithids are reconstructed as generally piscivorous with *Pengornis* diversifying into a more generalist and macrocarnivorous diet [9] (Fig. 1 right). In contrast, *Bohaiornis* and *Parabohaiornis* show adaptations reminiscent of extant herbivores while *Longusunguis* and possibly *Zhouornis* were adapted for taking vertebrate prey. The differences in the mass and talon shape of the latter pair further imply partitioning of the carnivorous niche.

The increased dietary breadth in Bohaiornithidae appears to stem from increased strength in the bohaiornithid skull. In particular, Longipterygidae and Bohaiornithidae may represent opposite evolutionary trajectories from the skull architecture of Pengornithidae, which is considered plesiomorphic for enantiornithines [86], and from the ancestral enantiornithine skull (Table 5). Past works [12, 19] have qualitatively described bohaiornithid skulls and teeth as robust, though when using ACH to quantify robusticity bohaiornithids are not particularly remarkable. While there is a large difference between longipterygids (ACH 0.09–0.11, x̄ = 0.10 [87]) and pengornithids (ACH 0.22–0.23, x̄ = 0.23 [88]), bohaiornithid skulls (ACH 0.21–0.26, x̄ = 0.24) are only slightly more robust than those of pengornithids. Ancestral state reconstruction (Fig. S7K) predicts Bohaiornithidae and Pengornithidae converged upon this level of robusticity from an ancestor whose ACH was between them and Longipterygidae. What we find more noteworthy than robusticity is a distinct stepwise increase in jaw strength between Jehol enantiornithine families: longipterygids have the weakest jaws (259–345 µε, x̄ = 309 µε [87]), with pengornithid jaws stronger (190–275 µε, x̄ = 222 µε [88]), and bohaiornithids’ stronger still (89–156 µε, x̄ = 128 µε). Ancestral state reconstruction supports bohaiornithids and longipterygids diversifying from a common ancestor with pengornithid-like jaw strength (Fig. S7), though a lack of jaw strength data from families outside these three limits the reliability of this reconstruction. An increase in in jaw strength may have allowed Bohaiornithidae to process a wider variety of difficult-to-process foods than contemporary groups and ecologically diversify, but the poor resolution of relationships within and near the clade [86, 89] (Table S1) makes this possibility difficult to test. For instance the skull of *Fortunguavis*, while overall poorly-preserved, does appear to have a robust surangular and pterygoid [90] and is commonly recovered as sister to Bohaiornithidae. Strengthening of the skull may therefore be present in more “bohaiornithid-like” enantiornithines (*sensu* [20]) and even convergent between them. *Shenqiornis*, the geologically oldest bohaiornithid, has a jaw strength intermediate within bohaiornithids (Fig. 5) which does imply the skull strengthening adaptations in Bohaiornithidae at least predate 124 Ma. Longipterygids, conversely, were limited by their weak and gracile jaws to relatively soft and compliant food sources [8]. However, whether the gracile jaws of longipterygids were adaptations to specialise in soft and compliant prey, or had unrelated benefits and coincidentally constrained their diet, remains unclear.

Bohaiornithids and Pengornithids are notably larger than other Jehol enantiornithines [2]. While we interpret only *Longusunguis* and possibly *Zhouornis* as inhabiting similar niches to pengornithids, this size increase does make invertivory unlikely in every pengornithid and bohaiornithid. The food web of the Jehol Biota is generally understood to be dominated by small animals [91, 92], particularly in the Jiufotang formation in which bohaiornithids and pengornithids are most diverse. Accordingly, one would expect interspecific competition for invertebrates to be high in this ecosystem. Bohaiornithidae and Pengornithidae (alongside the ornithurine family Songlingornithidae [2]), then, may be lineages which attempted to alleviate invertivorous competition by diversifying into consuming vertebrate or plant prey. Alternatively, size increase is also commonly interpreted as defence against predation. *Microraptor* was definitely capable of preying on smaller enantiornithines [93] and the Jehol biota had many carnivores larger than *Microraptor* [91, 92]. These possibilities need not be mutually exclusive: the more abundant large carnivores in the Yixian formation [91] may have provided pressure to grow larger when these clades first emerged, and the subsequent reduction in large carnivores by the Jiufotang formation [91] may have allowed them to further diversify. Indeed, ancestral state reconstruction (Fig. S5) predicts a concurrent increase in body mass for Bohaiornithidae and Pengornithidae near 125 Ma. Interestingly, though, it recovers the large mass of pengornithids as convergent, not present ancestrally in the clade. This is driven by the earliest pengornithid, *Eopengornis*, being smaller than its more recent relatives. No bohaiornithid is known from this timeframe, so we cannot yet comment on if their increased body mass is similarly convergent or ancestral to Bohaiornithidae as a whole.

#### The common ancestor of Enantiornithes and the speed of ecological diversification

Given the above trends, we can begin to speculate on the ancestral enantiornithine diet. The common ancestor of Enantiornithes likely had a body mass near 200 g (Fig. S5, Table 5), pointing to frugivory or generalist feeding. Its MA and jaw robusticity was likely intermediate to low (Fig. S6), overall resembling extant generalists, piscivores, and invertivores (Table 6). Its jaw strain during a bite is estimated at 227 µε (Table 5), within the range of Pengornithidae (Fig. S7) and thus consistent with folivory, swallowing granivory, generalist feeding, and invertivory [9]. The claws of the common ancestor of Enantiornithes are shaped in a way that is dissimilar from any extant bird included in this study, plotting between non-raptorial perchers and shrikes (Fig. S9). This is reminiscent of *Eopengornis*, and this claw shape has been interpreted as arboreal and allowing for very limited prey manipulation [9]. Overall, these lines of evidence point to the common ancestor of Enantiornithes being an arboreal generalist feeder. If this is the case, then the numerous ecological specialisations seen within Enantiornithes likely represent lineages already exploiting a resource further specialising in it, rather than dramatic trophic shifts. One enantiornithine lineage, that leading to both Longipterygidae and *Eoalulavis*, does appear to be ancestrally invertivorous (Fig. 9). Thus the purportedly vertivorous avisaurids [94, 95], late-diverging members of this lineage, would indeed represent a distinct increase in trophic level. However, as illustrated in Fig. 9, the diet of many of the earliest-diverging enantiornithines remains obscure, and these taxa strongly affect the interpretation of the ancestral state. Our quantitative reconstruction of the common ancestor of Enantiornithes in many ways resembles a smaller pengornithid, but that is to be expected when this family is much closer to the root than other taxa included in MA, FEA, and pedal TM reconstructions. This highlights the importance of further ecological study of the earliest-diverging enantiornithine taxa.

While the ancestral enantiornithine diet remains somewhat speculative (Fig. 9), it is clear that much ecological change and diversification occurred within Enantiornithes by 120 Ma. If current estimates of the Ornithothoraces node at 145 Ma [96, 97] are correct, then this level of ecological diversity took only 25 million years to achieve. For comparison, Neoaves (containing most trophic diversity in crown birds [1]) had only just diverged 25 million years after the origin of crown birds [98]. This suggests, then, that adaptations within crown Aves like edentulism [17, 99] or cranial kinesis [100] cannot fully explain the rate or extent of ecological diversification of crown birds after the K-Pg extinction (in agreement with several recent studies [58, 101]). In fact, crown birds may have been slower to occupy the same ecological breadth as their Mesozoic counterparts. This reframes some questions in evolutionary biology: while the question of what made crown birds uniquely fit to survive the K-Pg extinction [58, 100, 102] is still relevant, to find the traits which allowed for their ecological success and diversity we should look at Ornithothoraces as a whole. For instance, Ornithothoraces is defined by a variety of adaptations for improving flight capabilities [103]. Flight allows an organism easy access to multiple vertical strata within an environment, and flighted vertebrates tend to have larger home ranges than terrestrial vertebrates [104, 105]. Both of these factors provide volant animals with a greater variety and amount of resources, and may be a driver of repeated rapid diversification in ornithothoracine lineages.

## Methods

### Taxonomic Reference

We refer to extant taxa based on their genus and species in the Birds of the World database for consistency [41]. Within data files, taxa are referred to by their name in the data source (Skullsite Bird Skull Collection [106] or museum specimen designation). Comments in data files note where these identifications differ from Birds of the World or the bird diet database EltonTraits 1.0 [40]. Designations and relationships of fossil clades are based on [107], though use of “Bohaiornithidae” in this work is necessarily imprecise due to the instability of the clade (Table S1, [21]). For reasons explained below, we used Bohaiornithidae to refer to the six taxa referred when the clade was originally defined [12] plus *Beiguornis* [22].

### Taxa Included in Bohaiornithidae

The taxa included within Bohaiornithidae vary between studies. A summary of taxa included in the group in multiple phylogenies is provided in Table S1. The original definition of Bohaiornithidae is “The most recent common ancestor of *Shenqiornis mengi* and *Bohaiornis guoi*, and all its descendants” [12], but in practice this definition is used less strictly. Several studies [22, fig. 3B in 107, 108, 109] place the Bohaiornithidae node at an earlier diverging position than the strict clade definition in order to include *Zhouornis*, and others [19, 21, fig. 5a in 89, 110] recover *Shenqiornis* as so early-diverging that most or all enantiornithines would be considered bohaiornithids by the strict definition. Liu et al. [21] call for a redefinition of the clade after these findings, but do not provide a formal one. They place the Bohaiornithidae node on its phylogeny with a set of synapomorphies, but include *Gretcheniao* and *Junornis* in the clade which is not supported by subsequent studies [20, 22, 86]. Overall, the six original taxa proposed as members of Bohaiornithidae by [12] tend to be recovered as closely-related to *Bohaiornis*. *Longusunguis* is recovered in a clade with *Bohaiornis* in 16 of 22 (73%) studies, *Sulcavis* and *Zhouornis* in 18 of 23 (78%), *Shenqiornis* in 18 of 22 (82%), and *Parabohaiornis* in 19 of 21 (90%). Conversely, other taxa recovered in Bohaiornithidae tend to only rarely resolve near *Bohaiornis*. *Linyiornis* is recovered in a clade with *Bohaiornis* in 1 of 9 (11%) studies, *Eoenantiornis* in 3 of 21 (14%), and *Fortunguavis* in 4 of 19 (21%). *Gretcheniao* and *Musivavis*, both described as bohaiornithid-like but not bohaiornithid, are never recovered close to *Bohaiornis* (closest in [21] where *Gretcheniao* is in a clade sister to the six original bohaiornithids plus BMNHC-Ph1204). *Beiguornis* has only been included in one study [22] where it is recovered as a bohaiornithid, so it is tentatively included in the group pending future studies.

### Phylogenetic Tree Topology

Extant bird phylogenetic trees in this study are trimmed versions of the maximum clade credibility supertree used in [111]. This supertree follows [112], a tree time-scaled using Bayesian uncorrelated relaxed molecular clock data from 15 genes in 6,663 avian species constrained by seven fossil taxa. These data were mapped onto the backbone of [113] which used Bayesian uncorrelated relaxed molecular clock data from 259 genes in 200 species constrained by 19 fossil taxa. The tree files and R code to merge them were taken from the supplement of [111].

All grafted bohaiornithid branch lengths were scaled linearly so that the total length of the avian portion of the tree was equal to 94 Ma following the estimate of [98]. The Ornithothoraces node was placed at 145 Ma after Bayesian morphological clock analysis of two independent character sets [96, 97]. For the bohaiornithid portion of the tree, a topology recovered by two recent analyses [86, 89] was used where *Bohaiornis* and *Parabohaiornis* are sister taxa and the remaining bohaiornithids are in a basal polytomy. The position of *Beiguornis* is considered uncertain as the phylogeny in its description [22] differs from this topology, but it was not included in any analyses where phylogeny was incorporated.

### Bohaiornithid Fossil Dating

*Shenqiornis* is referred to the Qiaotou Formation [83]. This has been correlated to the Dawangzhangzi bed of the Yixian Formation [114], which is within the “Upper Undivided Yixian Formation” of [115] dated to 124 Ma. All other bohaiornithids are referred to the Jiufotang Formation. Most of these [12, 21] are from the Xiaotaizi/Lamadong locality dated to 119 Ma [116]. *Longusunguis* IVPP V18693 from the Lingyuan locality [53], *Zhouornis* BMNHC Ph756 from the Xiaoyugou locality [57], and *Zhouornis* CNUVB 903 of uncertain provenance [56] are referred to 121 Ma as the median age of the Jiufotang Formation [116].

### Taxonomic Status of BMNHC-Ph1204

BMNHC-Ph1204 was previously identified to Bohaiornithidae indet. [21], though we herein refine the diagnosis to cf. *Sulcavis*. As noted in [21], the large number of dentary teeth in BMNHC-Ph1204 is only seen in *Sulcavis* among known bohaiornithids, and the maxillary process of the nasal and interclavicular angle of the specimen are also consistent with those seen in *Sulcavis*. The final defining feature of BMNHC-Ph1204 noted by [21] is the lack of a fenestra in the maxilla, and the maxilla is poorly preserved in the *Sulcavis* holotype with no fenestra visible [15]. Most of the synapomorphies of *Sulcavis* are present in BMNHC-Ph1204: teeth with a flat lingual margin creating a ‘D-shaped’ cross-section [21 p. 5]; a broad nasal with a short, rostrally-directed maxillary process [21 p. 3]; the caudal-most process of the synsacrum extending far caudally to the vertebra’s articular surface [fig. 8 in 21]; a long and delicate acromion process on the scapula [fig. 7 in 21]; a convex lateral margin of the coracoid [fig. 8 in 21]; and an alular claw larger than that on the major digit [fig. 9 in 21]. The only missing synapomorphy is enamel with longitudinal ridges, autapomorphic for *Sulcavis* [15]. BMNHC- Ph1204 is less mature than the *Sulcavis* holotype (Fig. S10; [15, 21]), and ontogenetic change in enamel ornamentation has been noted previously in reptiles [117, 118], so it is possible that this is an ontogenetic difference. The same can be said of the additional carpal element in BMNHC- Ph1204, as carpometacarpal fusion is believed to occur relatively late in bohaiornithid ontogeny [12, 21]. Therefore, with this uncertainty from ontogeny, we refer BMNHC-Ph1204 to cf. *Sulcavis*. Should this specimen later be referred to its own taxon, the similarities above imply the new taxon would be very closely-related to *Sulcavis*, and this specimen would still be the most appropriate for filling in missing pieces of the skull of *Sulcavis*, as in Fig. 2.

**Figure 8.**
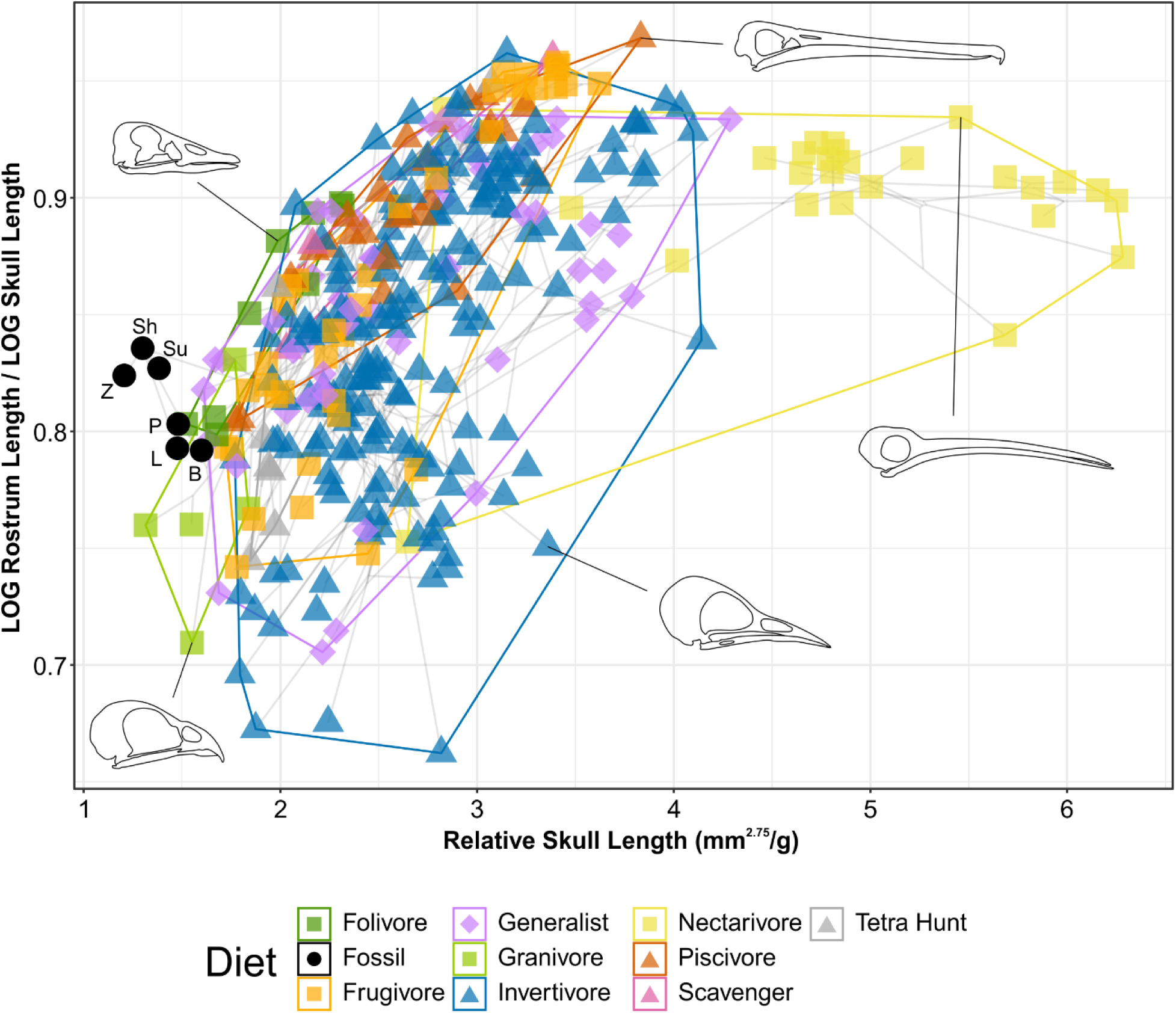
Phylomorphospace of extant and fossil bird skull proportions, grouped by diet. Grey lines indicate phylogenetic relationships. Line drawings of skulls for selected taxa are provided for reference. The data presented are modified from [10], see Fig. S4 for data more directly comparable to that study. Diet abbreviation: Tetra Hunt, Tetrapod Hunter. Fossil taxon abbreviations: B, *Bohaiornis*; L, *Longusunguis*; P, *Parabohaiornis*; Sh, *Shenqiornis*; Su, *Sulcavis*; Z, *Zhouornis*.

**Figure 9.**
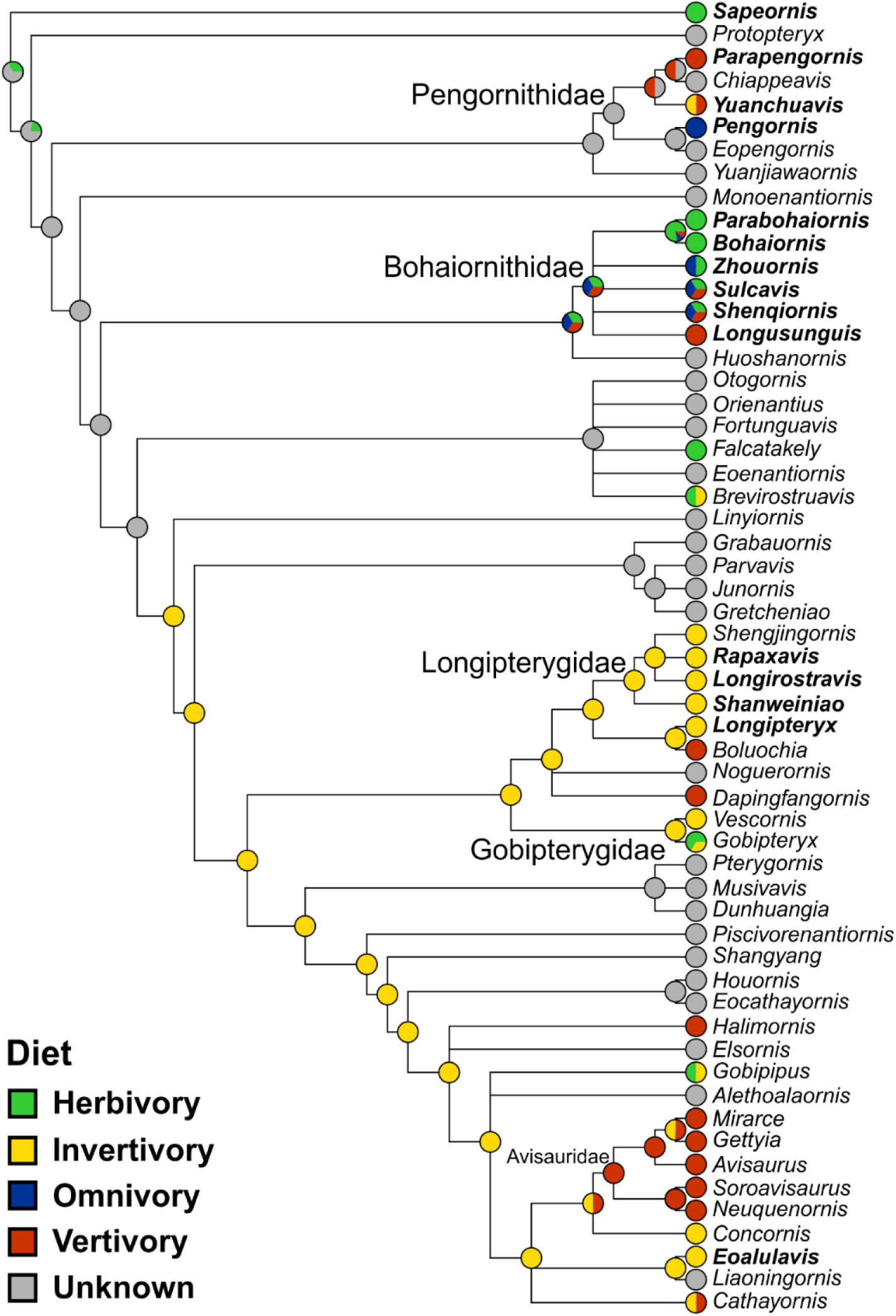
Ancestral state reconstruction of enantiornithine diet. Phylogeny is presented as not time-scaled for node visibility. All enantiornithine taxa with dietary hypotheses are included, as well as enantiornithines complete enough to create robust mass estimates [2, 8, 9, 26] and the non- ornithothoracine pygostylian *Sapeornis* as an outgroup. Taxa with bold names have diet assigned based on preserved meals (*Eoalulavis* [153], *Sapeornis* [3]) or quantitative diet proxies [8, 9]. Remaining diets assignments are based on qualitative morphology and depositional setting (Table S5). The diet of the common ancestor of Enantiornithes remains obscure, though many late-diverging enantiornithines are recovered as ancestrally invertivorous.

### Bohaiornithid Skull Reconstruction for MA and FEA

Reconstructions of bohaiornithid skulls are provided in Fig. 2. No complete bohaiornithid skulls are currently known, so digital reconstruction was necessary to create workable biological models [119]. Reconstructions of a given taxon used material from that taxon where possible, but each reconstruction required at least one bone from another taxon. Ideally these replacement bones would come from a taxon’s closest relative, but the relationships within Bohaiornithidae are unstable aside from the sister relationship of *Bohaiornis* and *Parabohaiornis*. For reconstruction purposes, we treated *Shenqiornis* and *Sulcavis* as sister taxa due to them having the next most frequent sister pairing among bohaiornithids [20, 57, 90] and the superficially similar shallow angle of their rostra. *Longusunguis* and *Zhouornis* have the most unstable placement within Bohaiornithidae, so supplemental parts were taken from taxa arbitrarily. Fortunately these taxa each had two moderately complete skulls, so little material from other taxa was needed.

Only *Bohaiornis* IVPP V17963 preserves the posteroventral region of the skull, so this region in all reconstructions is based on this. This affects jaw-opening mechanical advantage (OMA) of the upper jaw, and thus this functional index should be interpreted cautiously. We interpret the bone in *Bohaiornis* LPM B00167 previously labelled as the postorbital [55] to be the quadratojugal, as its shape is more similar to the short and weakly-forked quadratojugal typical of enantiornithines [120] than the elongate postorbital of *Longusunguis* IVPP V18693 [53], *Parabohaiornis* IVPP V28398 [121], *Sulcavis* BMNHC Ph-805 [15], and cf. *Sulcavis* BMNHC Ph-1204 [21]. The alternative is that this bone is in fact the postorbital, meaning *Bohaiornis* (and possibly *Shenqiornis* and *Zhouornis*) lacked a complete postorbital bar.

The jugal and postorbital bones of *Longusunguis* are elongate and believed to be in contact in life [53], which would increase the overall rigidity and stability of the skull. This is also very likely in *Parabohaiornis* and *Sulcavis*, which preserve both a dorsally-curving jugal and elongate postorbital (Fig. 2C,D). We reconstruct these bones as also in contact in other bohaiornithids (Fig. 2A,E,F), though this is less certain. Notably if we are incorrect in our reinterpretation of a bone in LPM B00167 identified as the postorbital [55] as the quadratojugal, then this much smaller element could not contact the jugal in life.

### Bohaiornithid Ontogeny

Ontogeny is an important dimension to account for when discussing the morphology and ecology of extinct animals, but the ontogeny of Bohaiornithidae (and enantiornithines in general) is poorly understood. Hu and O’Connor [122] devised a useful “character stage” system for tracking enantiornithine maturity based on the fusion of compound bones. Bohaiornithids are aged using this framework in Fig. S10. Hu and O’Connor [122] combine the fusion of the proximal metatarsus, the tibiotarsus, and carpometacarpus into a single Stage 3 (presumably due to a lack of resolution). *Longusunguis* IVPP V18693 has a fused tibiotarsus and unfused tarsometatarsus and carpometacarpus and *Sulcavis* BMNHC-Ph805 has a fused tibiotarsus and tarsometatarsus but unfused carpometacarpus, indicating the tibiotarsus fused first and the carpometacarpus last within this family. We thus subdivide the Stage 3 of [122] into 3a (formation of the tibiotarsus), 3b (proximal fusion of the tarsometatarsus), and 3c (fusion of the carpometacarpus) in bohaiornithids.

The above stages only approximate biological maturity, however. Histological data from two bones has been used to suggest that *Zhouornis* CNUVB-0903 had achieved sexual maturity but not skeletal maturity [56]; a recent study [123] has shown that histological maturity can vary greatly within individual enantiornithines but upholds the diagnosis of CNUVB-0903 as an adult. The indeterminate bohaiornithid CUGB P1202 is considered the least mature published member of the family [13]. It was proposed as sexually mature due to its elongate tail feathers (with the assumption they served a sexual display purpose) [13], but similar feathers in decidedly juvenile enantiornithines (STM 34-7 [124] and 34-9, IVPP V15564 [125]) call this into doubt. By correlating the histological maturity of [123] with the character stage system (Table S4), we see that subadulthood begins at or before character stage 1 and full maturity occurs after completion of character stage 3. We thus consider all bohaiornithids aside from the two mentioned above to be mature subadults or young adults, as proposed previously [12, 21, 53]. Most extant animals are considered “mature” upon reaching the subadult stage [126], so all specimens examined herein should be adequately mature for comparison to mature extant birds.

### Extant Sampling

Extant bird masses were taken from the bird ecology database AVONET [42], and combined with bird diet data from EltonTraits 1.0 [40] following the diet cut-offs in Table S2. The mass dataset consists of 8,758 birds: 169 folivores, 931 frugivores, 1,122 generalists, 475 granivores, 5,061 invertivores, 450 nectarivores, 207 piscivores, 35 scavengers, and 308 tetrapod hunters.

No additional extant birds are sampled for MA or FEA beyond the dataset of [9], meaning these datasets consist of 141 birds: nine folivores, 17 frugivores, 17 generalists, eight husking granivores, eight swallowing granivores, 43 invertivores, seven nectarivores, 15 piscivores, eight scavengers, and nine tetrapod hunters.

Five additional taxa were added to the ungual traditional morphometric (TM) dataset of [9] to increase the sample of anisodactyl non-raptorial perching birds. Target taxa were identified from the literature as birds with well-recorded perching behaviour and no record of grasping or manipulating prey with the pes. This list of target taxa was sent in a formal loan request to Serina Brady of Carnegie Museum of Natural History, who provided photographs of all available specimens fitting the criteria. In total, the extant TM dataset includes 66 birds: nine ground birds, 23 non-raptorial perching birds, four shrikes, 18 raptors taking large prey, and twelve raptors taking small prey.

Of the 408 extant bird samples included in the skull TM analysis of [10], 61 were removed due to not meeting the definition of any diet category in Table S2 (they were “in between” diets as defined here, often with 50% of the diet from two sources). Thus, the final skull morphometric dataset consisted of 347 birds across 298 species: nine folivores, 40 frugivores, 53 generalists, five granivores, 175 invertivores, 26 nectarivores, 27 piscivores, four scavengers, and eight tetrapod hunters.

### Analytical Techniques

#### Body Mass

The granivore diet category was not split into husking and swallowing granivores; past work [8, 9] did not find the two subcategories to meaningfully differ in body mass. Average masses of groups were compared via phylogenetic HSD (pairwise function of R package RRPP [43] used to compare group means). Diagnostic cut-off values between groups were found with the R package OptimalCutpoints [127] version 1.1-5 by optimizing the Youden index, with bootstrap estimation of 95% confidence intervals using the R package boot [128] version 1.3-28.

#### Pedal Traditional Morphometrics, Mechanical Advantage, and Finite Element Analysis

The extant MA and FEA data in this study is unchanged from [9], and new data collected from fossil specimens followed the same procedures as [9]. Pedal TM data are measured from scale photographs where available, though the five new included taxa were photographed without scale. All digits from a single individual were measured from a single photograph, so their relative scale is preserved. Our TM data is based solely on angles and length ratios, so this lack of scale should not be an issue. MA data were measured from the reconstructions in Fig. 2 in CorelDraw X8. Pedal ecological categories for TM follow [9]; of note, we define raptors taking large prey as those with records of regularly taking prey which cannot be completely encircled by the pes, and raptors taking small prey as those without such records. As pointed out in [36] this division commonly follows phylogenetic lines, but we observe exceptions such as *Bubo virginianus* taking large prey [129] and *Buteogallus anthracinus* not taking large prey [130] clustering with birds in the same pedal ecological group rather than their phylogenetic relatives. FEA models based on the reconstructions in Fig. 2 were created and solved within HyperWorks 2022 Student Edition (*HyperMesh* and *Optistruct*, Altair Engineering, Inc., USA). All analyses of the data (PCA, FDA, pFDA, ancestral state reconstruction, etc.) were performed in R version 4.1.2 [131], with scripts and raw data including measurements and FE models available from Mendeley Data (doi 10.17632/7xtpbv27zh.2).

#### Skull Traditional Morphometrics

Clark et al. [10] recently used skull traditional morphometrics to great effect in both differentiating extant bird diets and diagnosing the diet of longipterygid enantiornithines. Thus, we incorporate their data and methodology here as well to expand the evidence supporting our diet diagnoses. Measurements of skull and rostrum length for extant birds are unchanged from [10]. Clark et al. [10] do not diagram their measurement landmarks, and the caudal landmark of the rostral length as “the caudal margin of the lacrimal (i.e., the rostral margin of the orbit)” does not specify a dorsoventral height. So, for all new rostrum length measurements of bohaiornithids, we measured to the dorsoventral midpoint of the caudal margin of the ventral ramus of the lacrimal.

Diet assignments of extant taxa were changed from those used by Clark et al. [10] to those in Table S2 for consistency, following information in EltonTraits 1.0 [40]. We also use mass estimates for taxa from AVONET [42], rather than those listed by Clark et al. [10], for consistency within this work.

Clark et al. [10] visualise (their figure 4) and generally interpret their skull morphometrics in terms of proportional rostral length (i.e. rostrum length divided by skull length) vs. log10 skull length. We have modified both of these factors in Fig. 8. Firstly, we log10- transform both rostrum and skull length before taking their ratio, as this better normalises the distribution of proportional rostral length. Actual distribution of the data along this axis is minimally affected. Secondly, we use a relative skull length instead of absolute skull length.

Absolute skull length is essentially a size proxy; Fig. S4 shows skull length and body mass creating similar distributions, with the position of diet groups maintained relative to one another (while spreading more evenly along the x-axis when using body mass). This work already discusses body size as a diet proxy at length, so including it in morphometric discussion would be redundant. Through log10-log10 regression, we found that skull length generally increases with body mass with a scaling exponent of 2.75 (slightly negative allometry, R^2^ = 0.63) for the extant birds sampled. Thus we calculated relative skull length as:

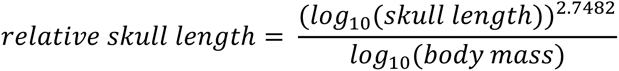

Relative skull length approximates how much longer or shorter a given bird’s skull is than the expected length for a bird of the same body mass.

#### Ancestral State Reconstruction

The backbone for the enantiornithine tree topology used in ancestral state reconstruction follows [86]. This tree contains 34 of the 56 species used in ancestral state reconstruction. Additional taxa were grafted onto this backbone following [20–22, 89, 110, 132–135]. Some taxa have only been included in one phylogenetic analysis, which placed said taxa within Bohaiornithidae, Longipterygidae, or Pengornithidae (e.g. *Noguerornis* placed within Longipterygidae [132]).

These taxa all lack the synapomorphies used to formally define the relevant clade, and their placements within them are likely artefacts of sharing a few characters while most data is missing. We therefore maintained monophyly of the Bohaiornithidae, Longipterygidae, and Pengornithidae by placing these ambiguous taxa as sister to the family they were placed within. Time-scaling was performed using the R package paleotree [136]. All enantiornithine taxa were placed at their earliest occurrence, with species divergence arbitrarily chosen to take 1,000 years. The enantiornithine phylogeny was rooted at 144 Ma after [96, 97]. Ages of localities follow [115, 116, 137–146], with precise notes in the data repository. If the horizon of a specimen was in doubt, it was placed at the median age of the formation.

The diet of the common ancestor of Enantiornithes was reconstructed in two ways: qualitatively, directly labelling diets onto taxa to reconstruct the diet of ancestral nodes; and quantitatively, reconstructing ancestral body mass, jaw MA and functional indices, jaw FEA MWAM strain and interval values, and pedal TM values, and predicting ancestral diet from these reconstructed values. The qualitative set includes quantitative diet reconstructions here and in [8, 9], all qualitative hypotheses of diet in enantiornithines (Table S5), and all enantiornithines with mass data (approximating enantiornithines known from substantial fossil material, as they need to be somewhat complete for mass estimation; *Cuspirostrisornis*, *Dalingheornis*, *Gracilornis*, *Jibeinia*, *Longchengornis*, *Microenantiornis*, and *Paraprotopteryx* were excluded due to never being included in a phylogenetic analysis). The latter were coded with “Unknown” diet to illustrate uncertainty in the reconstruction. As hypothesised diets are less precise than the diet categories used elsewhere in this study, they were lumped together as herbivores (folivores, frugivores, granivores, and nectarivores), invertivores, omnivores (generalists), or vertivores (piscivores, tetrapod hunters, and scavengers). 14 of 30 (47%) of diets in this reconstruction are supported by preserved meal evidence or quantitative diet proxies. A total of 56 taxa are included in the qualitative ancestral state reconstruction.

Quantitative ancestral state reconstruction includes data from the present study and our two other quantitative analyses of enantiornithines [8, 9], as well as pedal TM data for *Fortunguavis* from [147] and mass data from [2, 26]. Only MWAM strain of FEA models was reconstructed, not full intervals data, as we found the validity of reconstructing an individual strain interval to be dubious. If pedal TM data was available for multiple specimens of a genus, we favoured the most mature specimen to represent the genus. If mass data was available for multiple specimens of a genus, we favoured the largest mass estimate as an assumption of it representing the most mature sample. The large-toothed *Longipteryx* morphotype [8] was assumed more mature than the small-toothed morphotype and used for all reconstructions. A total of 44 taxa are used in ancestral state reconstruction of body mass, 13 in upper jaw MA, 9 in lower jaw MA, 13 in FEA, and 16 in pedal TM.

*Sapeornis* was used as the outgroup in ancestral state reconstruction. Mass data for the taxon was taken from [26]. MA data was measured from the reconstruction in [148]. FEA data was taken from a model constructed by Yuen Ting (Athena) Tse for an in-progress collaborative work based on the same reconstruction [148]. TM data were taken from [147]. Its qualitative diet was classified as herbivorous based on preserved seed meals [3]. Its divergence from Enantiornithes was placed at 147 Ma after [97].

Ancestral states were estimated using the fastAnc() function in Phytools 0.7-90 [149]. As polytomies are present in the enantiornithine tree used, ancestral states calculated 10,000 times with random resolutions of the polytomies and averaged. As some diets were uncertain (e.g. *Zhouornis* may be either herbivorous or a generalist), and fastAnc() cannot accept probabilistic qualitative states for terminal taxa, we coded diets quantitatively as percent likelihood for each diet.

## Acknowledgements

The authors would like to thank Serina Brady, Chase Mendenhall, Stephen Rogers (Carnegie Museum of Natural History), Andrew Kratter, and David Steadman (Florida Museum of Natural History) for their assistance and expertise in selecting physical specimens for this study. We would also like to thank Kathryn C. Gamble DVM, MS, Dipl ACZM, Dip ECZM (ZHM), Veterinary Advisor Coraciiformes/Bucerotiformes TAG for providing coraciiform radiographs used in this study. We also thank Gavin Thomas and Ryan Felice for their insight in constructing the consensus phylogeny of extant birds. Finally, we thank Yuen Ting (Athena) Tse for constructing the FEA model of *Sapeornis* used as an outgroup in ancestral state reconstruction. CVM is supported by a Postgraduate Scholarship from The University of Hong Kong (HKU PGS), the Jurassic Foundation, and a Stephen Jay Gould Student Research Award from the Paleontological Society. MP is supported by the Research Grant Council of Hong Kong’s General Research Fund (17120920; 17103315; 17105221) and the School of Life Sciences at The Chinese University of Hong Kong. X.W. is supported by the Taishan Scholars Program of Shandong Province (Ts20190954).

