## supplemental information for "Trophic diversity and evolution in Enantiornithes: a synthesis including new insights from Bohaiornithidae"

#### Supplemental Text

#### Results

##### Body Mass

Of the 9,994 bird species in [1], 1,236 birds either had no mass data or did not fall into one of our diet categories (Table S2), leaving a sample size of 8,758. Significant phylogenetic signal was found in extant bird mass, with trends resembling those expected under a Brownian motion (BM) model of evolution (K = 0.93; Table S6).

##### Mechanical Advantage and Functional Indices

Significant phylogenetic signal is present in the extant MA dataset overall (Table S6) and in each individual functional index (Table S7). Indices are usually less similar than expected under a BM model (K = 0.35–0.78), with the exception of relative average cranial height (ACH; K = 1.15). Phylogenetic honest significant differences (HSD; i.e. comparison of means using the pairwise function in R package RRPP [2]) recovered no change in significant differences from HSD with the older phylogeny used in [3] (Table S8). Skull reconstructions used in MA and FEA calculations are provided in Fig. 2.

Sensitivity analysis of the quadrate placement in fossil taxa (Fig. S11) agrees with [3, 4] that anterior shifts make folivory more likely and posterior shifts make piscivory more likely (Table S3). Scavenging was recovered as likely for *Parabohaiornis* and folivory was recovered as likely for *Bohaiornis* regardless of the quadrate's position.

#### Finite Element Analysis

No significant phylogenetic signal was recovered in the extant FEA data (Table S6). Phylogenetic HSD does recover less significant differences in diet than [3]. We no longer find significant differences between folivores and husking granivores or scavengers, between frugivores and scavengers, or between husking granivores and invertivores (Table S9).

#### Traditional Morphometrics

##### Pes

Significant phylogenetic signal is present in the extant pedal TM dataset overall (Table S6) and in each individual measurement (Table S10). Measurements are all less similar than expected under a BM model, with measurements involving digits I and II having higher K values. Phylogenetic HSD recovered no change in significant differences from HSD with the older phylogeny used in [3] (Table S11).

The additional extant data has produced little change from [3], primarily extending the morphospace of non-raptorial perching birds into more negative PC1 space (Fig. 7A) driven by the inclusion of *Podargus strigoides*.

##### Skull

Significant phylogenetic signal is present in the skull TM data. When using the data as-is, a K value near zero is returned, driven by the variation observed within the taxa sampled multiple times. While this has interesting implications for the importance of individual variation in morphometric studies, for simplicity in this work we averaged the traits of all species with multiple samples when calculating phylogenetic signal. Skull and rostrum length distribution both resemble a BM model, while the ratio between them is less similar than expected under BM and relative skull length is more similar (Table S12).

Granivores and tetrapod hunters tend to have relatively short rostra (rostrum length/skull length < 0.8) while folivores, nectarivores, and piscivores tend to have more elongate rostra (rostrum length/skull length > 0.8). Frugivores, generalists, and invertivores display a variety of rostral proportions.

Nectarivores have distinctly elongate skulls (relative skull length > 4.5), though this is exclusively true for hummingbirds (Trochilidae). Somewhat elongate skulls (relative skull length > 3.5) are primarily found in invertivores and generalists, though these diet groups overall display a range of relative skull lengths.

#### Ancestral State Reconstruction

Precise values for nodes of interest in Fig. 9 are provided here. The common ancestor of Avisauridae is recovered as 100% likely to be a vertivore. The common ancestor of Bohaiornithidae is recovered as 34% likely to be an herbivore, 33% likely to be a vertivore, and 33% likely to be an omnivore. The common ancestor of Longipterygidae is recovered as 100% likely to be an invertivore. The diet of the common ancestor of Pengornithidae is recovered as 99% unknown, with the next most likely diet invertivory at 0.02%. The same is true for the common ancestor of all Enantiornithes. The common ancestor of *Linyiornis* and a large group of late-diverging enantiornithines is recovered as 99% likely to be an invertivore

(1% unknown), and this is true for the rest of the backbone of the tree crownward from this point.

### Discussion

#### Body Mass

An increased sample size has improved resolution of the relationship between mass and diet over our previous work [3, 4]. Notably, the separation between vertivores and invertivores is at a lower mass (324–429 g in [3, 4], vs 80–148 g here). This is of particular note for interpreting longipterygid diet. In a previous study [4], we found *Longipteryx* unlikely to be a piscivore due to its low body mass, but three specimens of *Longipteryx* (DNHM D2889, est. 154 g; IVPP V12325, est. 193 g; and STM 8-117, est. 206 g) are above the mass cutoff in this study. While MA results do still indicate piscivory not being particularly likely in *Longipteryx* [4], we believe that with this refined mass data the hypothesis of *Longipteryx* as a specialist piscivore [5-10, 11 pg. 83 and 270, 12] can no longer be rejected. Specific analogy to kingfishers, whose jaw strain is lower than *Longipteryx* under loading, is still rejected.

#### Mechanical Advantage and Functional Indices

The quadrate is disarticulated in every published bohaiornithid skull, and its position is highly influential on MA measurements. Our sensitivity analysis of the quadrate position (Fig. S11, Table S3) indicates that folivory cannot be ruled out for any bohaiornithid. In fact, it is the most likely diet for every taxon except *Parabohaiornis* when the quadrate is shifted to the anteriormost plausible position. Unlike in longipterygids and pengornithids [3, 4], scavenging is generally not recovered as likely with a posterior quadrate shift except in *Parabohaiornis*, and in this taxon it is also recovered as likely with an anterior shift as well. A posterior-shifted quadrate does, however, make piscivory plausible in *Sulcavis* and *Zhouornis* and the most likely diet for *Shenqiornis*. A posterior quadrate shift also makes invertivory more likely than generalist feeding for *Zhouornis*.

#### Traditional Morphometrics

##### Pes

It has come to our attention [13] that in the work our pedal morphometric results were originally reported in [4] only one non-raptorial perching taxon has an anisodactyl toe arrangement, as in all known enantiornithines. Most are zygodactyl (Cuculidae, Psittaciformes) or semi-zygodactyl (Musophagidae) [14]. The category revision in [3] increased anisodactyl non-raptorial perching to 5 taxa, but this still represents only about 25% of the non-raptorial perching taxa in the sample. In response to this, we added an additional 5 representatives of non-raptorial perching taxa with anisodactyl toe arrangements (Cracidae, Fregatidae, Meropidae, Phoeniculidae, Podargidae). This increased the anisodactyl percentage of the non-raptorial perching birds to 43%.

The new extant data had minimal effect on ecological category boundaries and phylogenetic signal of pedal morphometrics. The largest difference between these results and those in [3] is the non-raptorial perching bird space infiltrating a space of slightly less claw curvature (driven by the blunt unguals of the tawny frogmouth *Podargus strigoides*) and shrikes being slightly more distinct in pFDA. No categories gain or lose significant differences in phylogenetic HSD from [3]. Given the minimal change caused by adding more anisodactyl birds, that the anisodactyl birds plot alongside our zygodactyl non-raptorial perchers, and that owls (Strigidae) and the osprey (Pandionidae) are semi-zygodactyl [14] but plot far from zygodactyl non-raptorial perchers, we believe that toe arrangement is not a driving force of the trends observed in our pedal morphometric data. Certain toe arrangements in which digits I and II are not the primary grasping digits, such as heterodactyly, are not included in this study and may have meaningfully different patterns of claw curvature and size.

Bohaiornithid claw curvature and relative size are generally conserved through ontogeny (Fig. 7A), more like *Bubo virginianus* [fig. 3 in 15] than *Longipteryx* [fig. 5 in 4]. Thus, ontogeny is not considered a major factor in claw TM in this study.

Skull

Extant results of our skull TM analysis generally resemble those presented by Clark et al. [16]. Our Fig. S4A, compared to their figure 4, identifies additional separation within herbivorous birds with granivores and folivores separating along the LOG skull length axis. Combining the insectivore and (non-insectivorous) invertivore classes of Clark et al. [16] had little effect on the data, as the latter was completely within the former's region of the morphospace in the original dataset. We additionally plotted the ratio of rostrum length to skull length against LOG body mass to investigate if the separation along the skull length axis was a size effect, and the results in Fig. S4B support this hypothesis. The structure of the data is overall similar with a slightly more normal spread along the x-axis, but group separation along the x-axis is maintained and overall follows the larger trends in body mass presented in this work.

As size trends in diet are already discussed here, we also investigated the relative length of the skull as an alternative variable to plot against the relative rostral length (Fig. 8). Nectarivorous taxa largely display very long skulls relative to body size, though this is driven entirely by hummingbirds (Trochilidae). Hummingbirds are known to have skulls that are peramorphic relative to other Strisores [17], possibly to support requisite neural tissue despite miniaturisation [18]. Aside from this group, relative rostrum length and relative skull length share a general positive correlation. In other words, elongation of the rostrum tends to parallel elongation of the skull. This is unsurprising, as the rostrum in extant birds has relatively high integration with the rest of the skull (tables S2 and S3 in [19]). General relationships between skull and rostrum length and diet parallel those we discussed previously [4] for AMA and OMA. Short-skulled birds (e.g. granivores) have relatively sturdy jaws more efficient at processing hard and tough foods, while those with longer jaws (e.g. piscivores) can more efficiently move their jaws at high speeds and thus better catch quick-moving prey.

### Supplemental Figures

#### Figure S1

Violin plots of individual functional indices, organized by diet. AMA (A,B), PMA (C,D), OMA (E,F), AO (G,H), MCH (I), MMH (J), ACH (K),and AMH (L) values are provided for the upper (A,C,E,G,I,K) and lower (B,D,F,H,J,L) jaws. Character abbreviations: AMA, anterior jaw-closing mechanical advantage; PMA, posterior jaw-closing mechanical advantage; OMA, jaw-opening mechanical advantage; AO, relative articular offset; MCH, relative maximum cranial height; MMH, relative maximum mandible height; ACH, relative average cranial height; AMH, relative average mandible height. Diet abbreviations: GranivoreH, Husking Granivore; GranivoreS, Swallowing Granivore; Tetra Hunt, Tetrapod Hunter. Fossil taxon abbreviations: B, *Bohaiornis*; L, *Longusunguis*; P, *Parabohaiornis*; Sh, *Shenqiornis*; Su, *Sulcavis*; Z, *Zhouornis*.

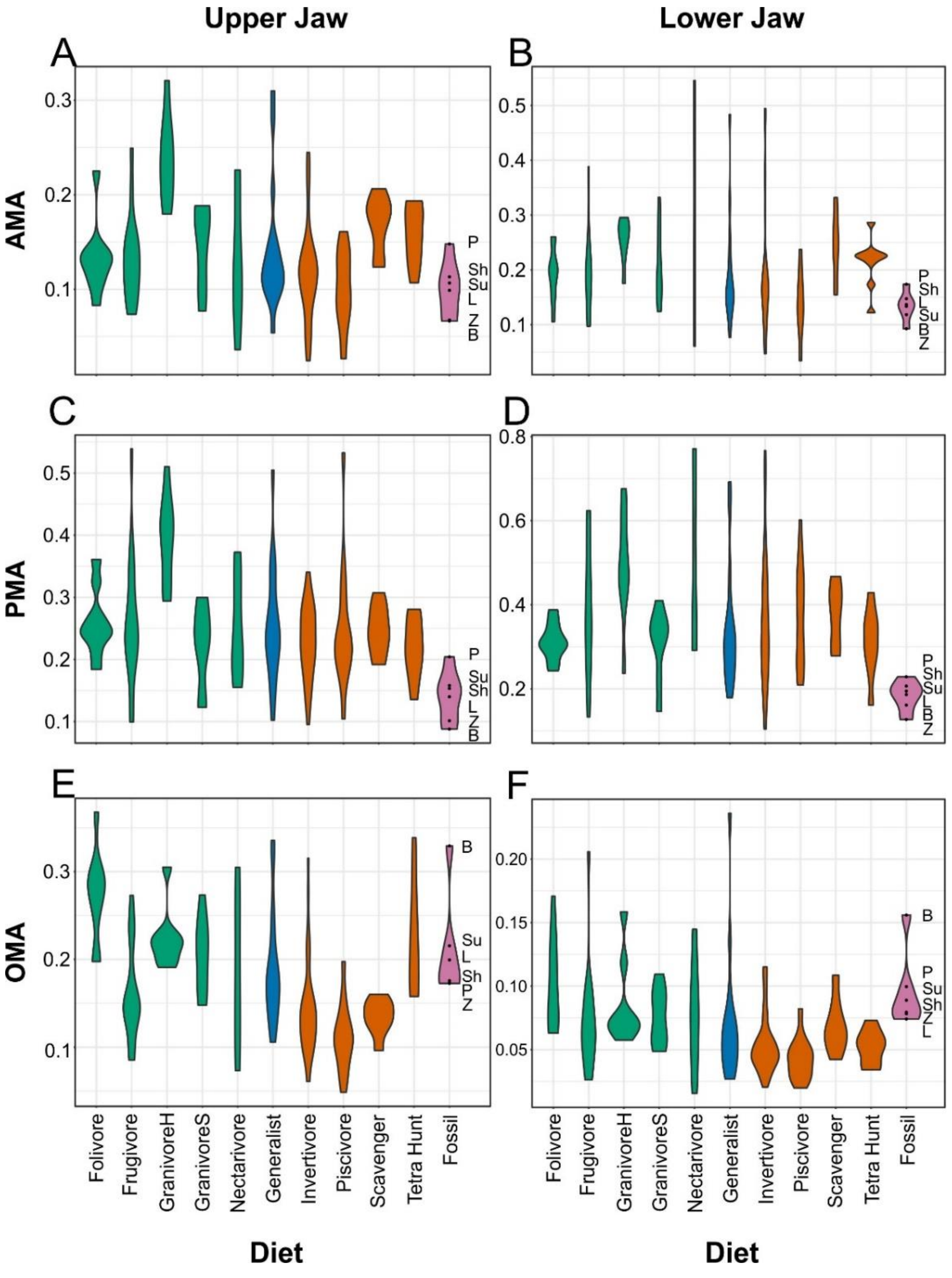

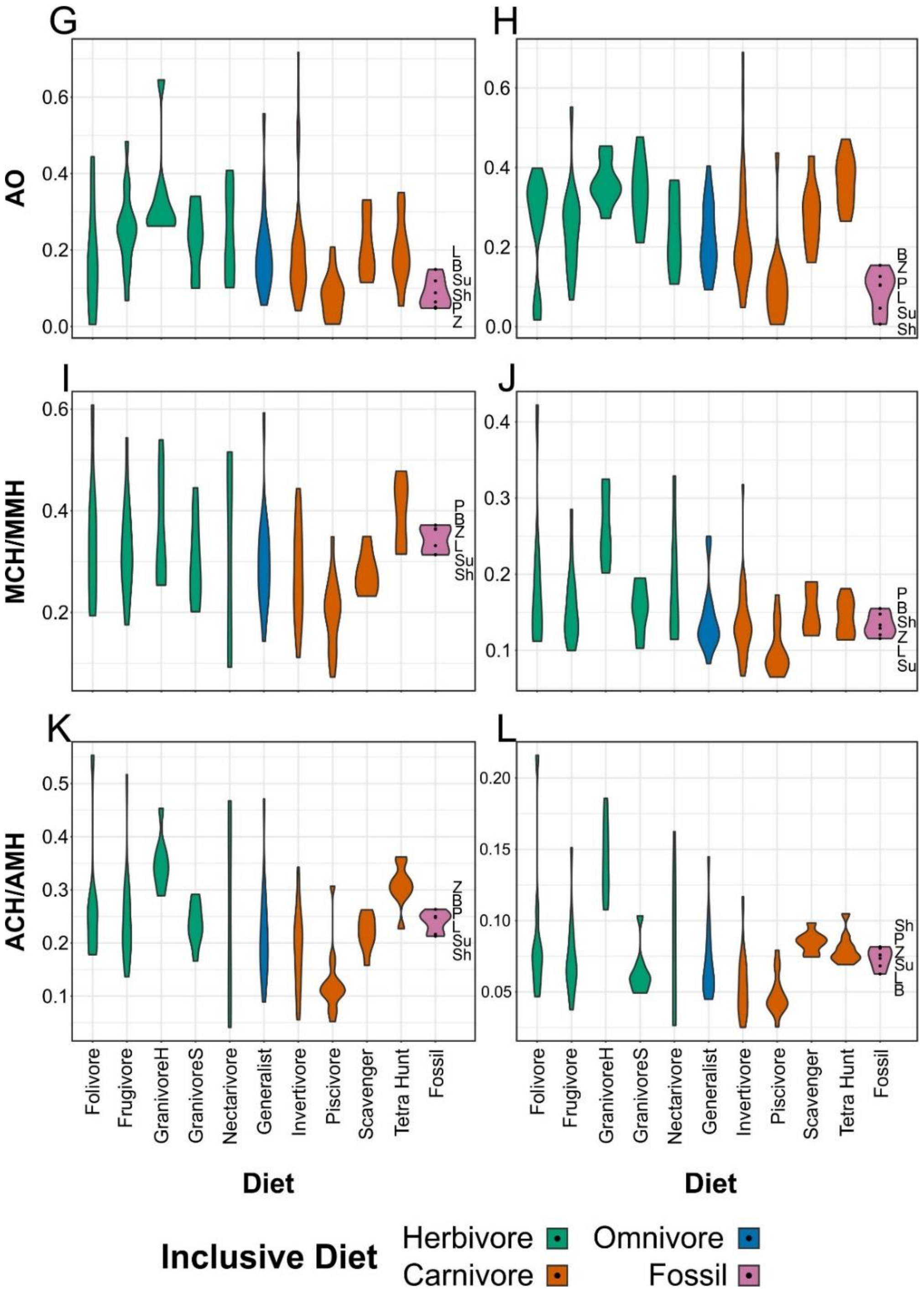

G
H
I
J
K
L
AO
MCH/MMH
ACH/AMH
Folivore
Frugivore
GranivoreH
GranivoreS
Nectarivore
Generalist
Invertivore
Piscivore
Scavenger
Tetra Hunt
Fossil
Diet
Inclusive Diet
Herbivore
Carnivore
Omnivore
Fossil

Figure S2
Plot of character weightings for the graphs in Fig. 4. Plots are provided for PCA (A) and FDA (B). Character abbreviations: AMA, anterior jaw-closing mechanical advantage; PMA, posterior jaw-closing mechanical advantage; OMA, jaw-opening mechanical advantage; AO, relative articular offset; MCH, relative maximum cranial height; MMH, relative maximum mandible height; ACH, relative average cranial height; AMH, relative average mandible height. Suffixes of upr and lwr respectively denote the measurement is taken of the upper or lower jaw. Diet abbreviations: GranivoreH, Husking Granivore; GranivoreS, Swallowing Granivore; Tetra Hunt, Tetrapod Hunter.

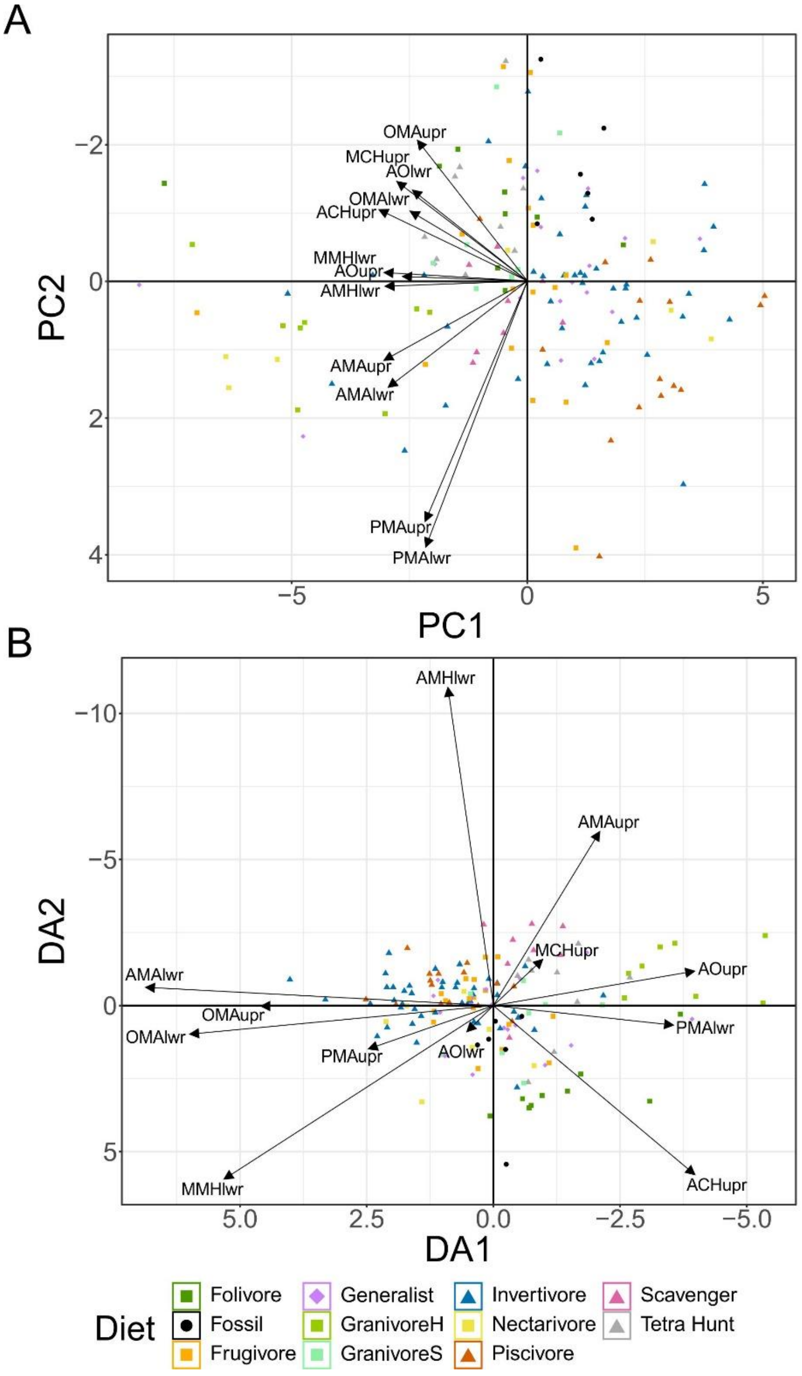

Figure S3
Plot of character weightings for the graphs in Fig. 7. Plots are provided for PCA (A) FDA (B), and pFDA (C). Category abbreviations: Large Raptor, raptor taking prey which does not fit in the foot; Small Raptor, raptor taking prey which can fit in the foot.

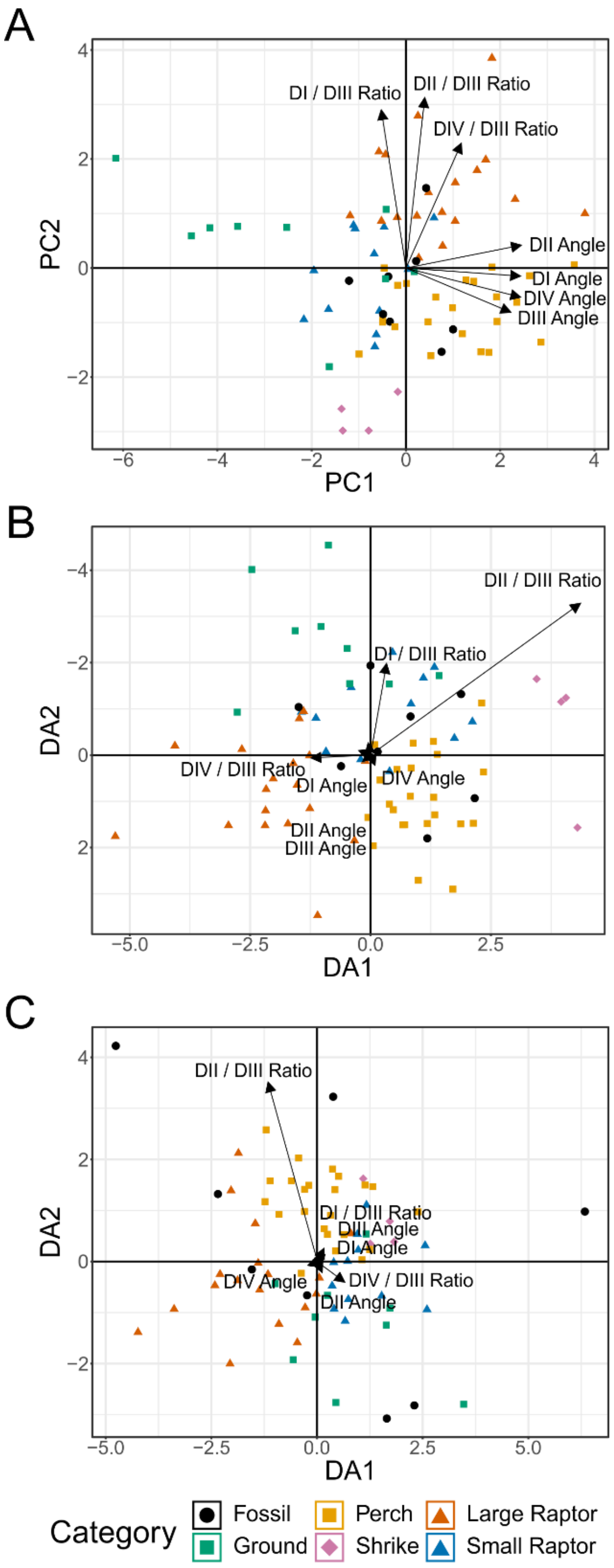

Figure S4
Phylomorphospace of extant and fossil bird skull proportions, grouped by diet. Grey lines indicate phylogenetic relationships. Axes are presented as shown in [16] (A) and with mass as the x-axis to investigate if spreading is driven by size effects (B). As we interpret these trends as being size-dependent, most discussion of skull proportions in this work are based on Fig. 8. Diet abbreviation: Tetra Hunt, Tetrapod Hunter. Fossil taxon abbreviations: B, *Bohaiornis*; L, *Longusunguis*; P, *Parabohaiornis*; Sh, *Shenqiornis*; Su, *Sulcavis*; Z, *Zhouornis*.

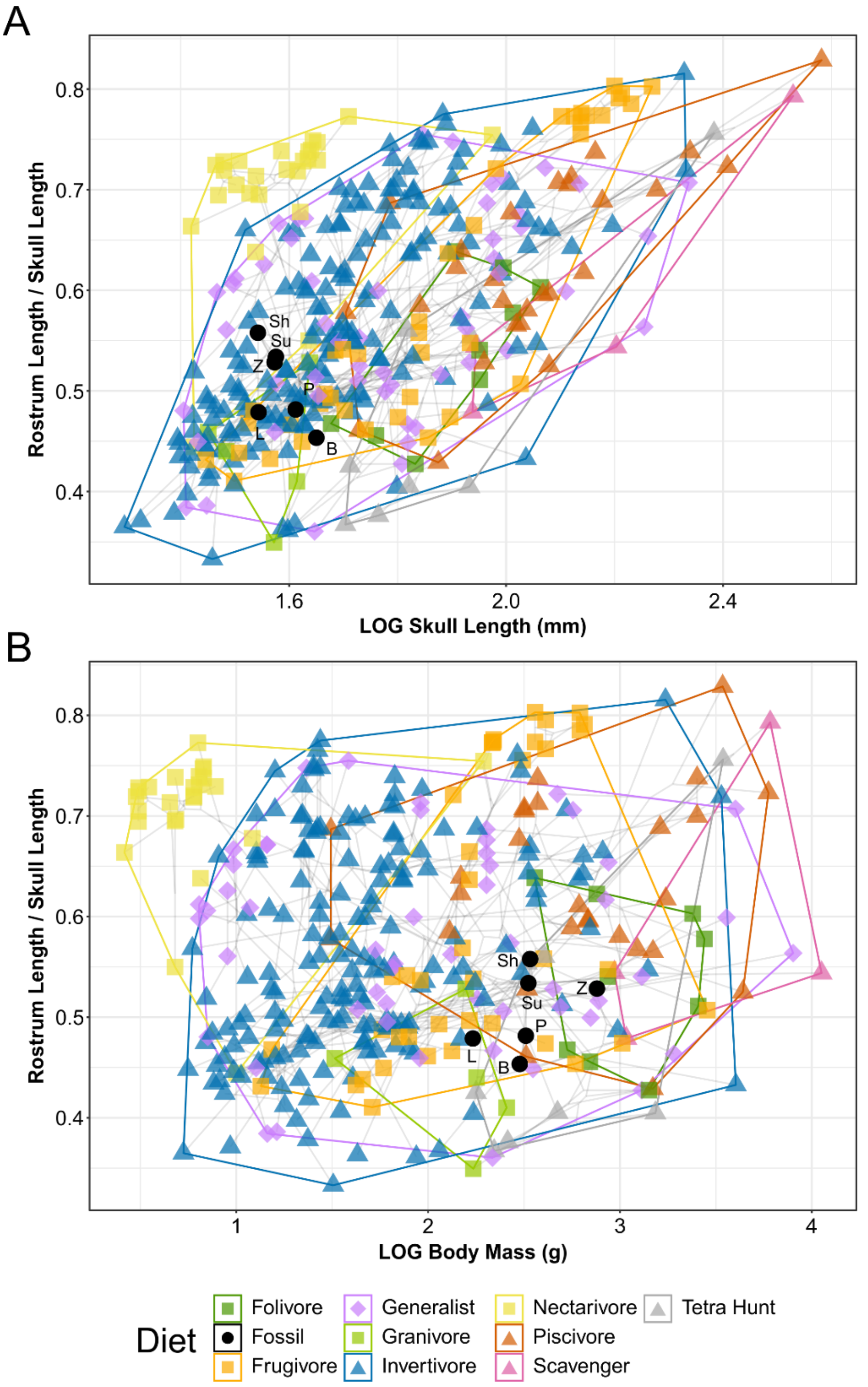

Figure S5
Ancestral state reconstruction of enantiornithine body mass. Sapeornis is included as an outgroup. Mass data is taken from Table 1 and past work [3, 4, 20]. Tree is time-scaled and resolves polytomies in the order taxa appear on the tree. This does not account for other possible permutations of the tree topology; a more valid mass estimate for the common ancestor of Enantiornithes is provided in Table 5.

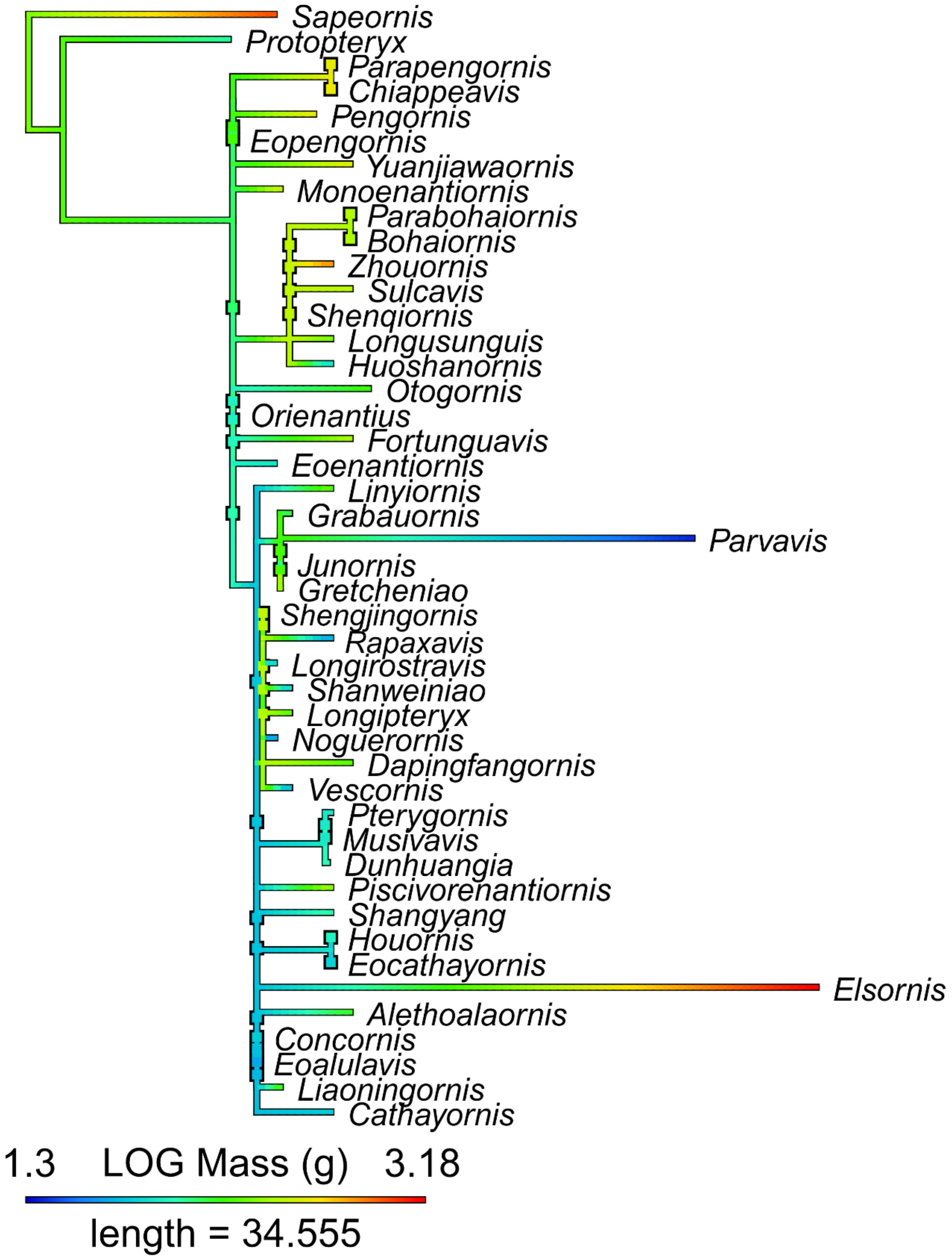

Figure S6

Ancestral state reconstruction of enantiornithine MA and functional indices. Sapeornis is included as an outgroup. AMA (A,B), PMA (C,D), OMA (E,F), AO (G,H), MCH (I), MMH (J), ACH (K),and AMH (L) reconstructions are provided for the upper (A,C,E,G,I,K) and lower (B,D,F,H,J,L) jaws. Character abbreviations: AMA, anterior jaw-closing mechanical advantage; PMA, posterior jaw-closing mechanical advantage; OMA, jaw-opening mechanical advantage; AO, relative articular offset; MCH, relative maximum cranial height; MMH, relative maximum mandible height; ACH, relative average cranial height; AMH, relative average mandible height. Suffixes of upr and lwr respectively denote the measurement is taken of the upper or lower jaw. Enantiornithine data is taken from this study and past work [3, 4]; the outgroup Sapeornis is measured from the reconstruction in [21]. Tree is time-scaled and resolves polytomies in the order taxa appear on the tree. This does not account for other possible permutations of the tree topology; more valid estimates of variates for the common ancestor of Enantiornithes is provided in Table 5.

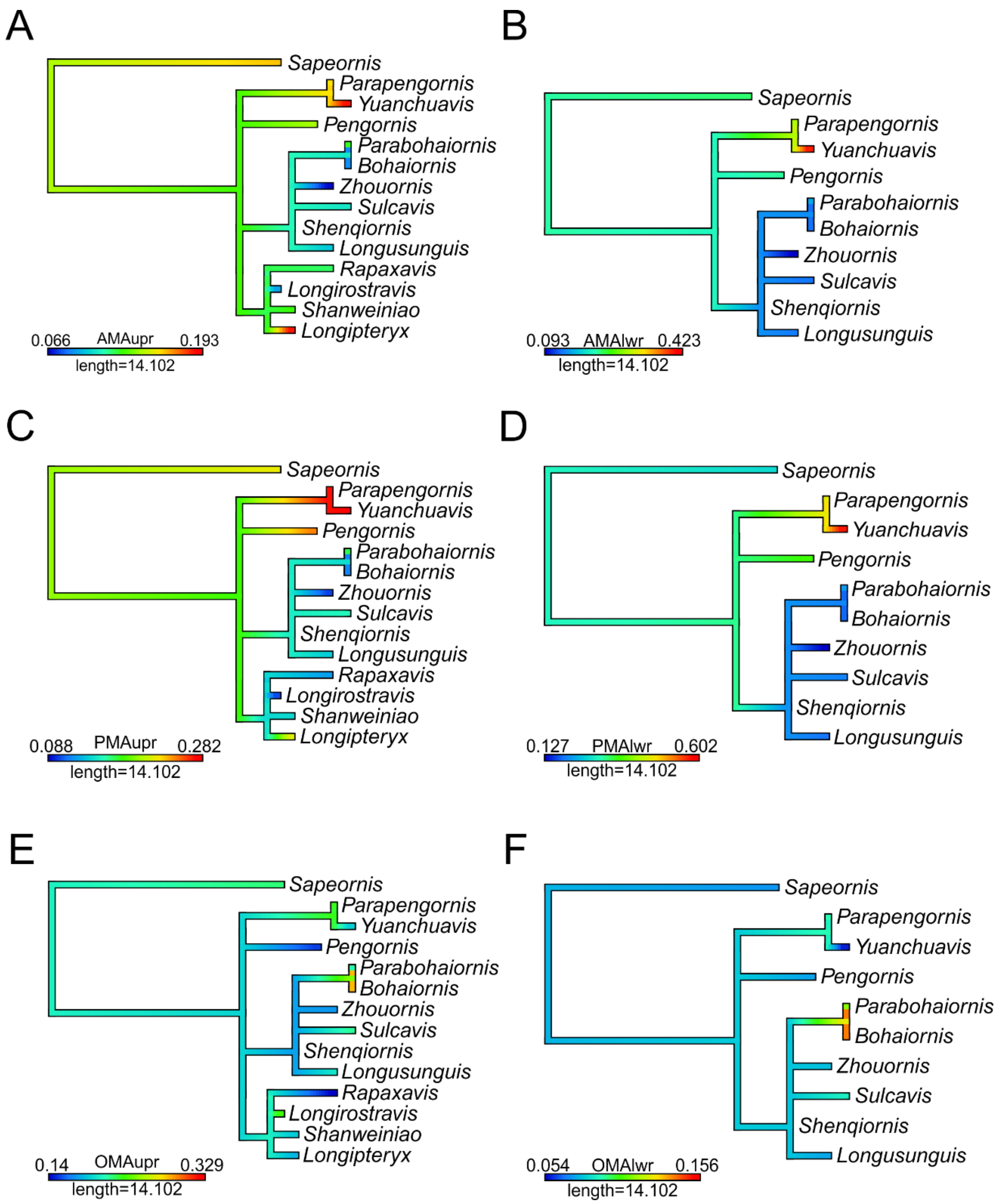

A
Sapeornis
Parapengornis
Yuanchuavis
Pengornis
Parabohaiornis
Bohaiornis
Zhouornis
Sulcavis
Shenqiornis
Longusunguis
Rapaxavis
Longirostravis
Shanweiniao
Longipteryx
0.066 AMAupr 0.193
length=14.102
B
Sapeornis
Parapengornis
Yuanchuavis
Pengornis
Parabohaiornis
Bohaiornis
Zhouornis
Sulcavis
Shenqiornis
Longusunguis
0.093 AMAlwr 0.423
length=14.102
C
Sapeornis
Parapengornis
Yuanchuavis
Pengornis
Parabohaiornis
Bohaiornis
Zhouornis
Sulcavis
Shenqiornis
Longusunguis
Rapaxavis
Longirostravis
Shanweiniao
Longipteryx
0.088 PMAupr 0.282
length=14.102
D
Sapeornis
Parapengornis
Yuanchuavis
Pengornis
Parabohaiornis
Bohaiornis
Zhouornis
Sulcavis
Shenqiornis
Longusunguis
0.127 PMAlwr 0.602
length=14.102
E
Sapeornis
Parapengornis
Yuanchuavis
Pengornis
Parabohaiornis
Bohaiornis
Zhouornis
Sulcavis
Shenqiornis
Longusunguis
Rapaxavis
Longirostravis
Shanweiniao
Longipteryx
0.14 OMAupr 0.329
length=14.102
F
Sapeornis
Parapengornis
Yuanchuavis
Pengornis
Parabohaiornis
Bohaiornis
Zhouornis
Sulcavis
Shenqiornis
Longusunguis
0.054 OMAlwr 0.156
length=14.102

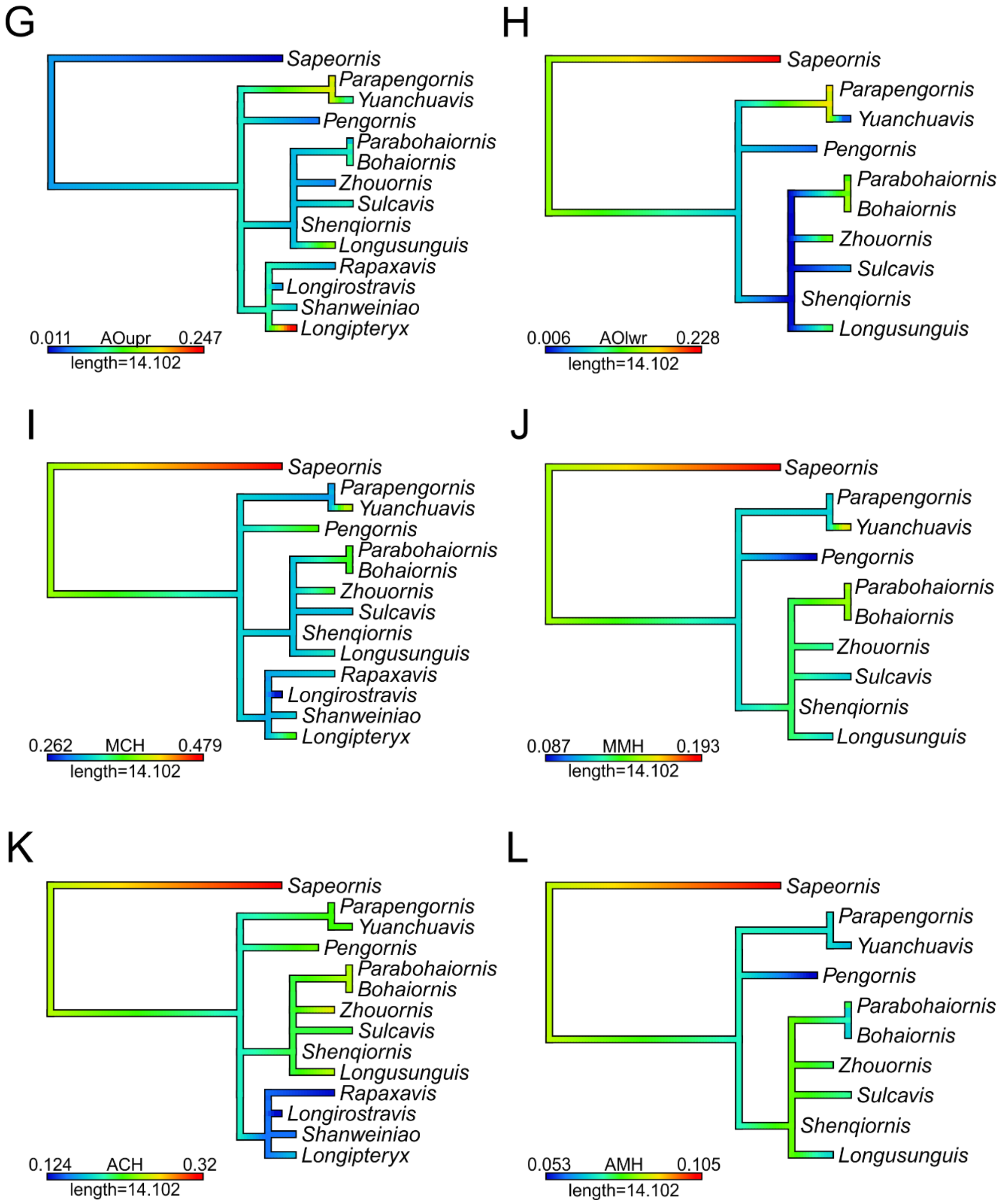
G
Sapeornis
Parapengornis
Yuanchuavis
Pengornis
Parabohaiornis
Bohaiornis
Zhouornis
Sulcavis
Shenqiornis
Longusunguis
Rapaxavis
Longirostravis
Shanweiniao
Longipteryx
0.011 AOupr 0.247
length=14.102
H
Sapeornis
Parapengornis
Yuanchuavis
Pengornis
Parabohaiornis
Bohaiornis
Zhouornis
Sulcavis
Shenqiornis
Longusunguis
0.006 AOlwr 0.228
length=14.102
I
Sapeornis
Parapengornis
Yuanchuavis
Pengornis
Parabohaiornis
Bohaiornis
Zhouornis
Sulcavis
Shenqiornis
Longusunguis
Rapaxavis
Longirostravis
Shanweiniao
Longipteryx
0.262 MCH 0.479
length=14.102
J
Sapeornis
Parapengornis
Yuanchuavis
Pengornis
Parabohaiornis
Bohaiornis
Zhouornis
Sulcavis
Shenqiornis
Longusunguis
0.087 MMH 0.193
length=14.102
K
Sapeornis
Parapengornis
Yuanchuavis
Pengornis
Parabohaiornis
Bohaiornis
Zhouornis
Sulcavis
Shenqiornis
Longusunguis
Rapaxavis
Longirostravis
Shanweiniao
Longipteryx
0.124 ACH 0.32
length=14.102
L
Sapeornis
Parapengornis
Yuanchuavis
Pengornis
Parabohaiornis
Bohaiornis
Zhouornis
Sulcavis
Shenqiornis
Longusunguis
0.053 AMH 0.105
length=14.102

Figure S7

Ancestral state reconstruction of enantiornithine MWAM strain under bite loading in FEA. Sapeornis is included as an outgroup. Enantiornithine data is taken from this study and past work [3, 4]; data from *Sapeornis* is taken from a model constructed by Yuen Ting (Athena) Tse for an in-progress collaborative work based on the skull reconstruction in [21]. Tree is time-scaled and resolves polytomies in the order taxa appear on the tree. This does not account for other possible permutations of the tree topology; a more valid MWAM strain estimate for the common ancestor of Enantiornithes is provided in Table 5.

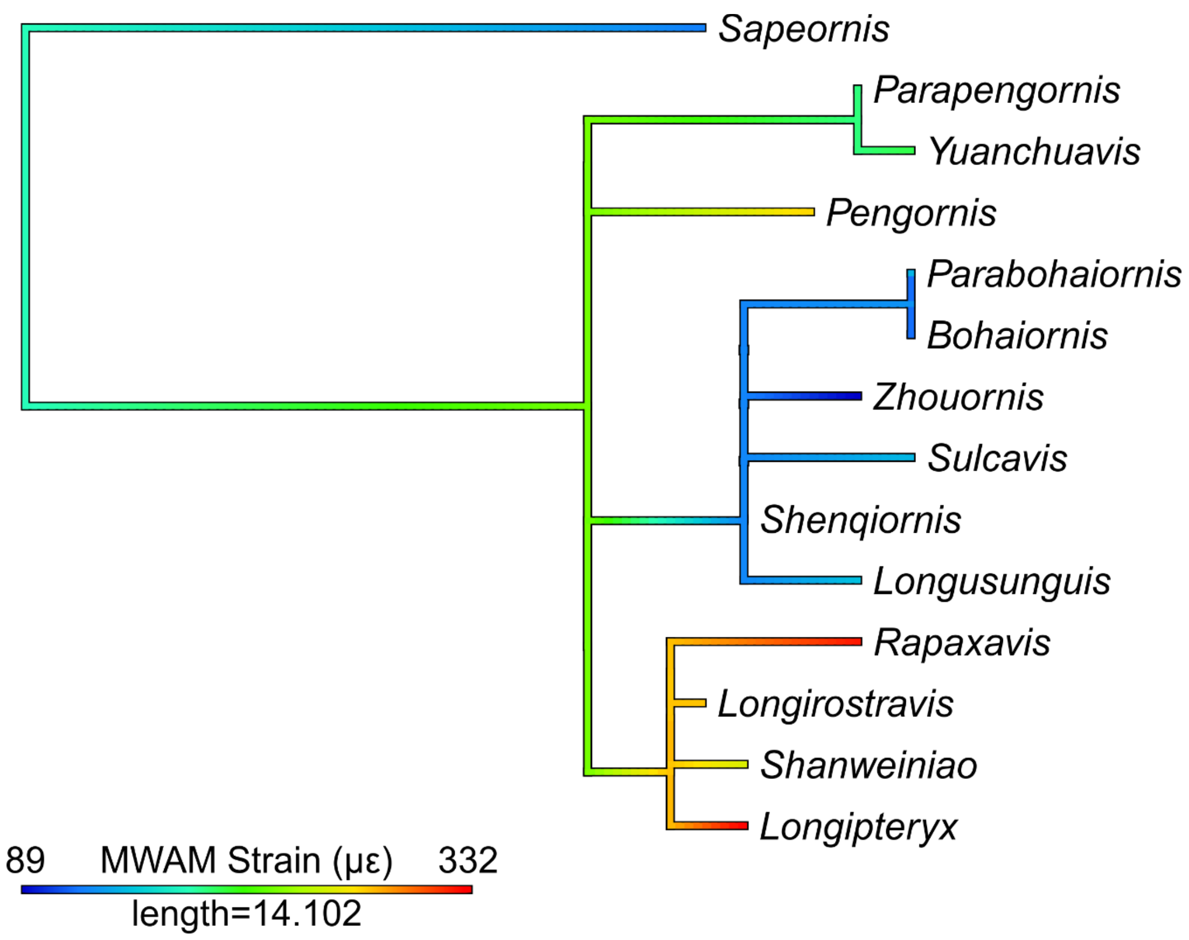

Figure S8

Ancestral state reconstruction of enantiornithine pedal TM variables. Sapeornis is included as an outgroup. Reconstructions of ratios of DI (A), DII (B), and DIV (C) arc length relative to DIII arc length as well as DI (D), DII (E), DIII (F), and DIV (G) angles are provided. Data is taken from this study and past work [3, 4, 22]. Tree is time-scaled and resolves polytomies in the order taxa appear on the tree. This does not account for other possible permutations of the tree topology; more valid estimates of variates for the common ancestor of Enantiornithes is provided in Table 5.

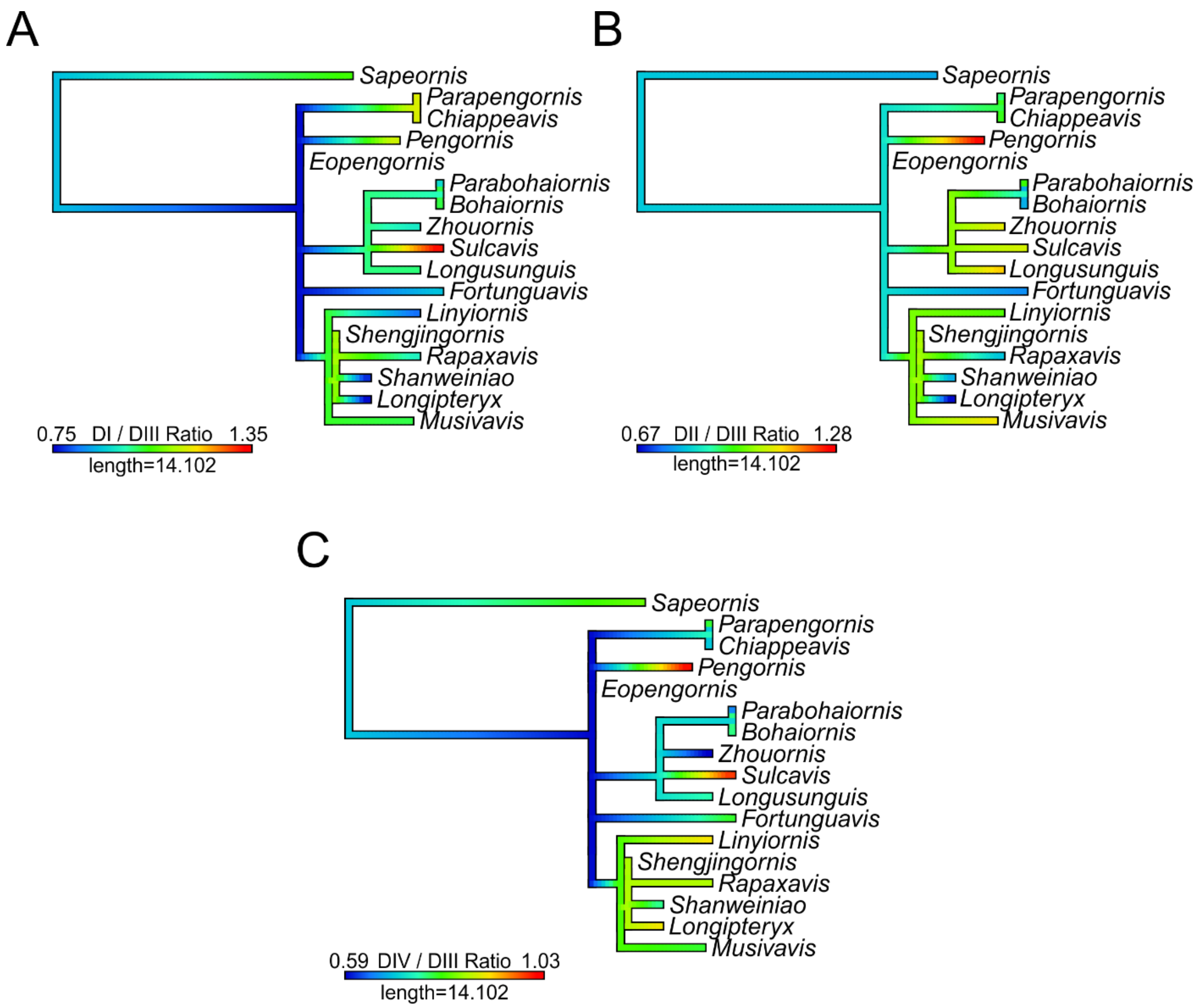
A
Sapeornis
Parapengornis
Chiappeavis
Pengornis
Eopengornis
Parabohaiornis
Bohaiornis
Zhouornis
Sulcavis
Longusunguis
Fortunguavis
Linyiornis
Shengjingornis
Rapaxavis
Shanweiniao
Longipteryx
Musivavis
0.75 DI / DIII Ratio 1.35
length=14.102
B
Sapeornis
Parapengornis
Chiappeavis
Pengornis
Eopengornis
Parabohaiornis
Bohaiornis
Zhouornis
Sulcavis
Longusunguis
Fortunguavis
Linyiornis
Shengjingornis
Rapaxavis
Shanweiniao
Longipteryx
Musivavis
0.67 DII / DIII Ratio 1.28
length=14.102
C
Sapeornis
Parapengornis
Chiappeavis
Pengornis
Eopengornis
Parabohaiornis
Bohaiornis
Zhouornis
Sulcavis
Longusunguis
Fortunguavis
Linyiornis
Shengjingornis
Rapaxavis
Shanweiniao
Longipteryx
Musivavis
0.59 DIV / DIII Ratio 1.03
length=14.102

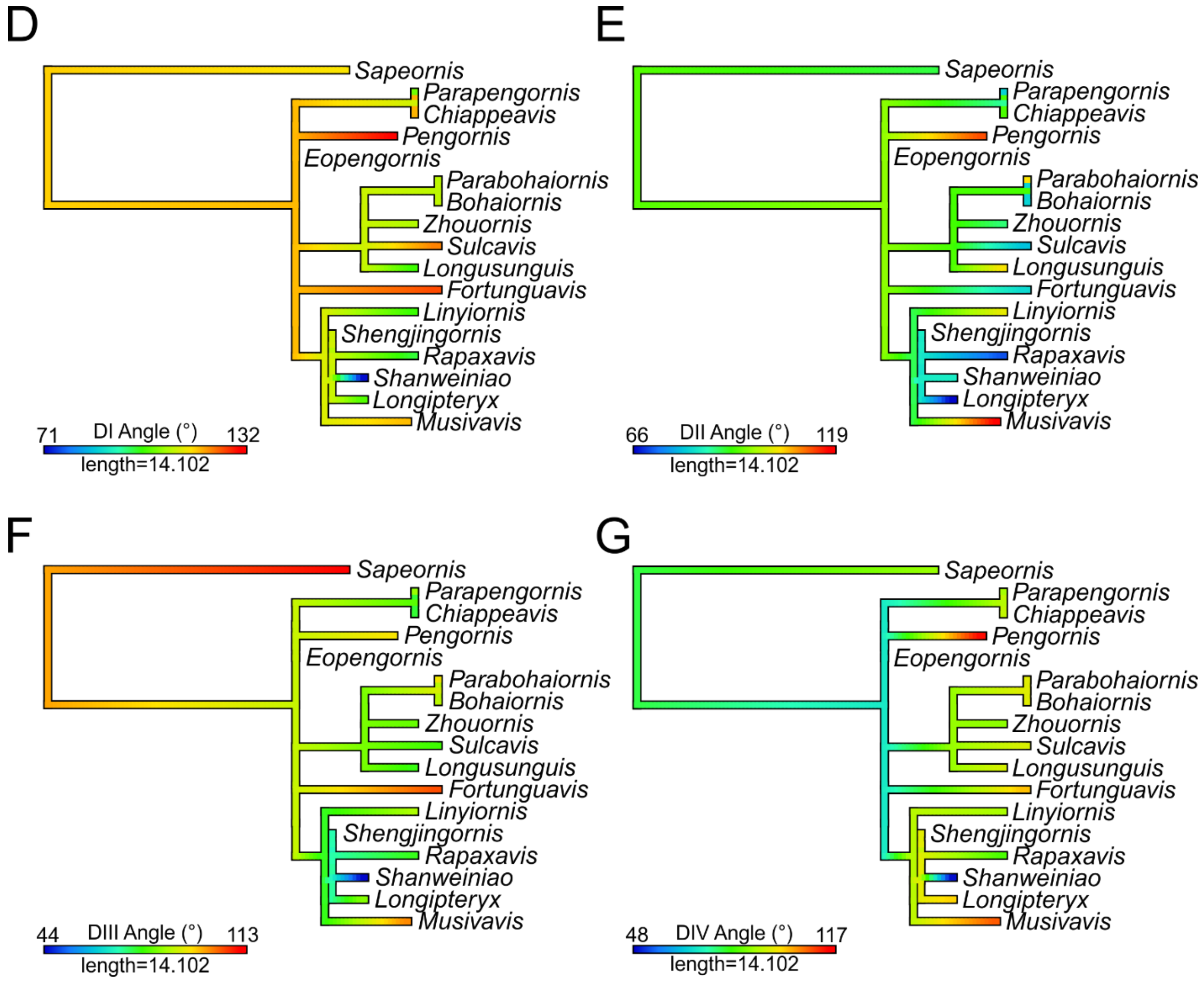
D
Sapeornis
Parapengornis
Chiappeavis
Pengornis
Eopengornis
Parabohaiornis
Bohaiornis
Zhouornis
Sulcavis
Longusunguis
Fortunguavis
Linyiornis
Shengjingornis
Rapaxavis
Shanweiniao
Longipteryx
Musivavis
71 DI Angle (°) 132
length=14.102
E
Sapeornis
Parapengornis
Chiappeavis
Pengornis
Eopengornis
Parabohaiornis
Bohaiornis
Zhouornis
Sulcavis
Longusunguis
Fortunguavis
Linyiornis
Shengjingornis
Rapaxavis
Shanweiniao
Longipteryx
Musivavis
66 DII Angle (°) 119
length=14.102
F
Sapeornis
Parapengornis
Chiappeavis
Pengornis
Eopengornis
Parabohaiornis
Bohaiornis
Zhouornis
Sulcavis
Longusunguis
Fortunguavis
Linyiornis
Shengjingornis
Rapaxavis
Shanweiniao
Longipteryx
Musivavis
44 DIII Angle (°) 113
length=14.102
G
Sapeornis
Parapengornis
Chiappeavis
Pengornis
Eopengornis
Parabohaiornis
Bohaiornis
Zhouornis
Sulcavis
Longusunguis
Fortunguavis
Linyiornis
Shengjingornis
Rapaxavis
Shanweiniao
Longipteryx
Musivavis
48 DIV Angle (°) 117
length=14.102

Figure S9
Phylomorphospace of extant bird claw shape from TM, grouped by pedal ecological category, for comparison to the common ancestor of Enantiornithes. See Fig. S3A for character weights. Category abbreviations: Large Raptor, raptor taking prey which does not fit in the foot; Small Raptor, raptor taking prey which can fit in the foot.

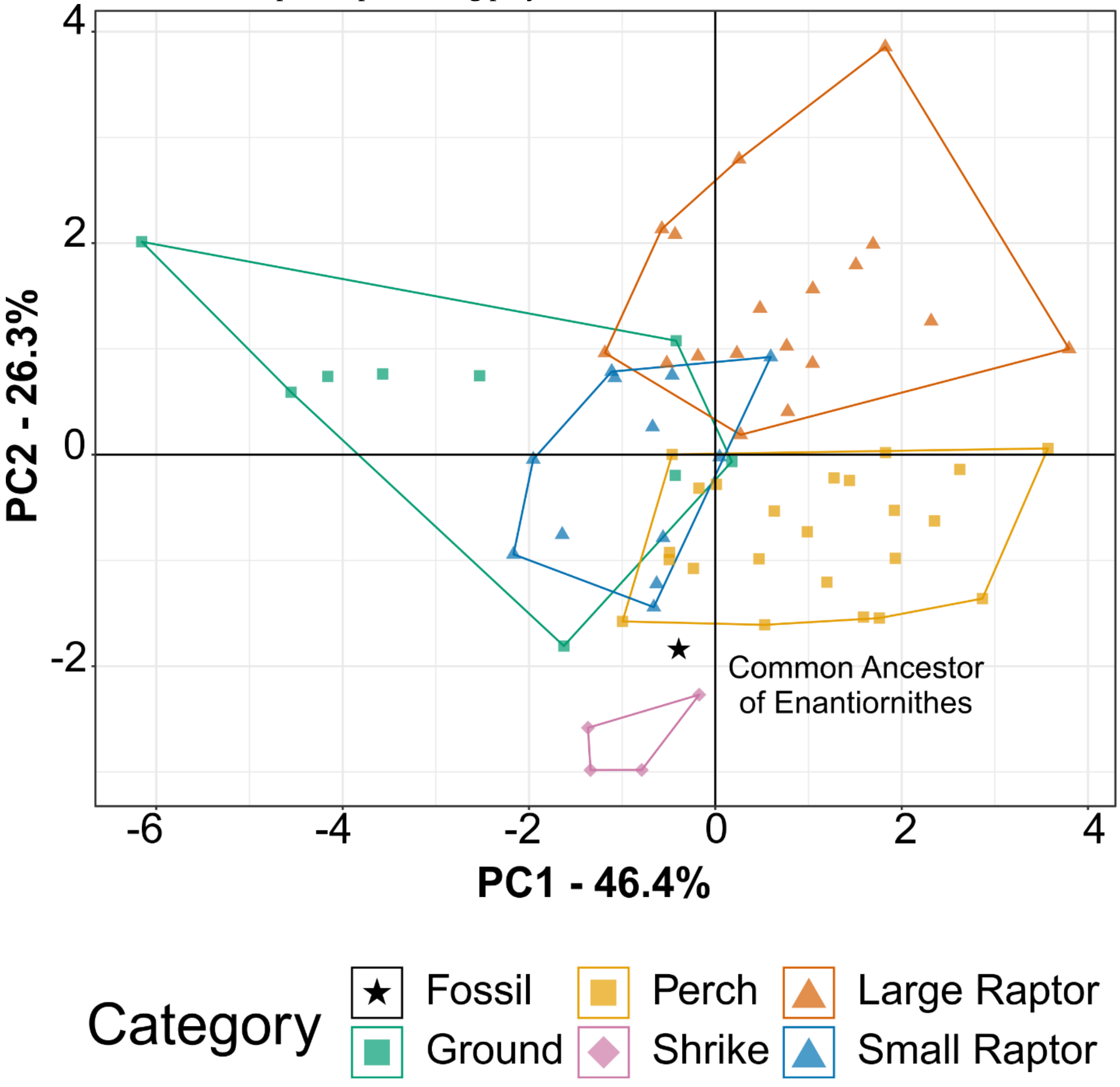

Figure S10

Ontogenetic stages of bohaiornithids. Stages are based on [23], with new subdivisions of Stage 3 (possibly specific to Bohaiornithidae) as noted in Methods. Subadult status is reached at or before Stage 1 and adulthood within Stage 3; see Table S4 for details.

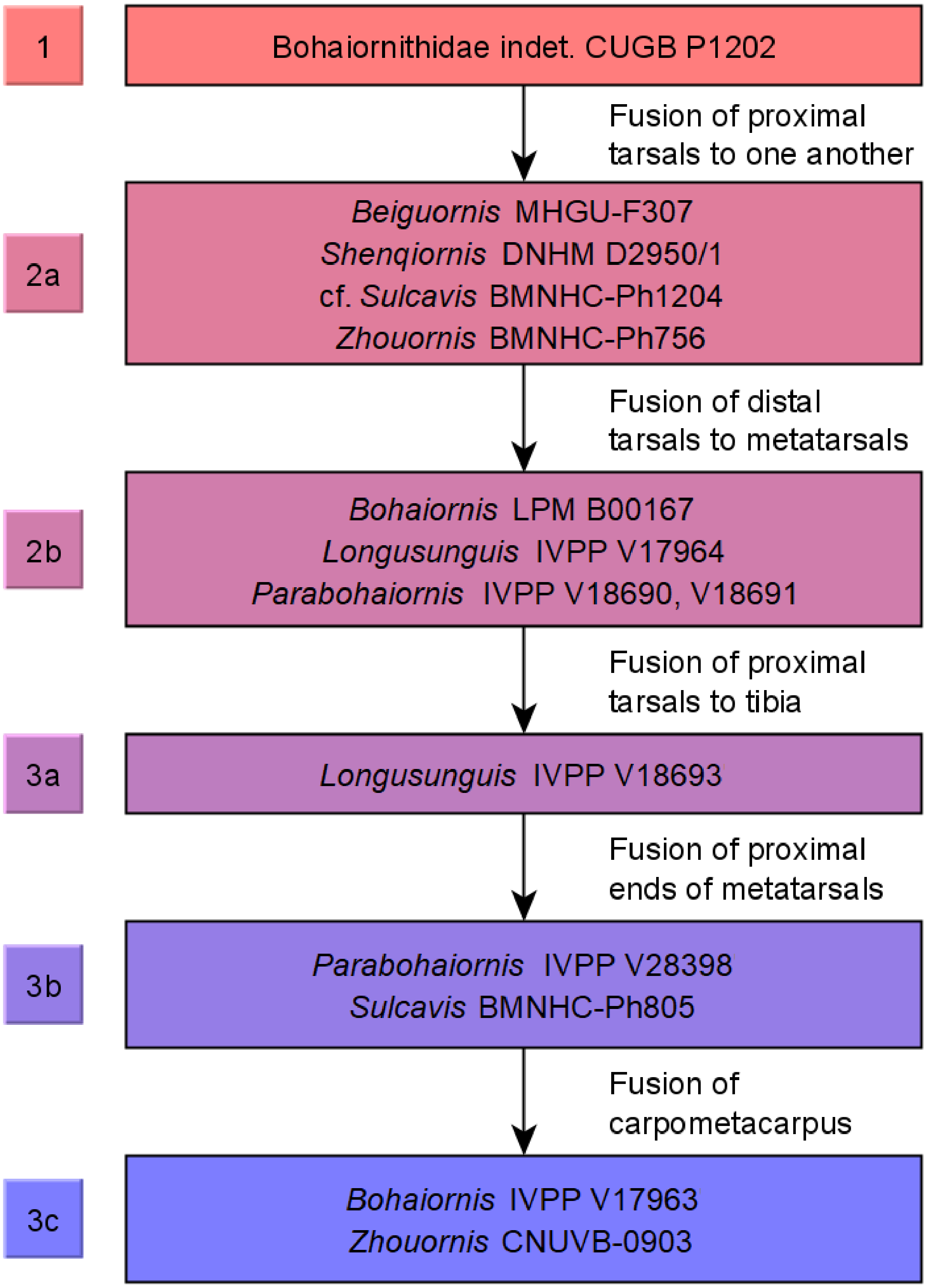

Figure S11

Reconstructions of bohaiornithid skulls for sensitivity analysis of the quadrate's position, which is uncertain due to disarticulation in all specimens. Reconstructions of *Bohaiornis* (A,G), *Longusunguis* (B,H), *Parabohaiornis* (C,I), *Shenqiornis* (D,J), *Sulcavis* (E,K), and *Zhouornis* (F,L) are constructed with the quadrate shifted as far anteriorly (A–F) or posteriorly (G–L) as biologically plausible. Results of the sensitivity analysis are provided in Table S3.

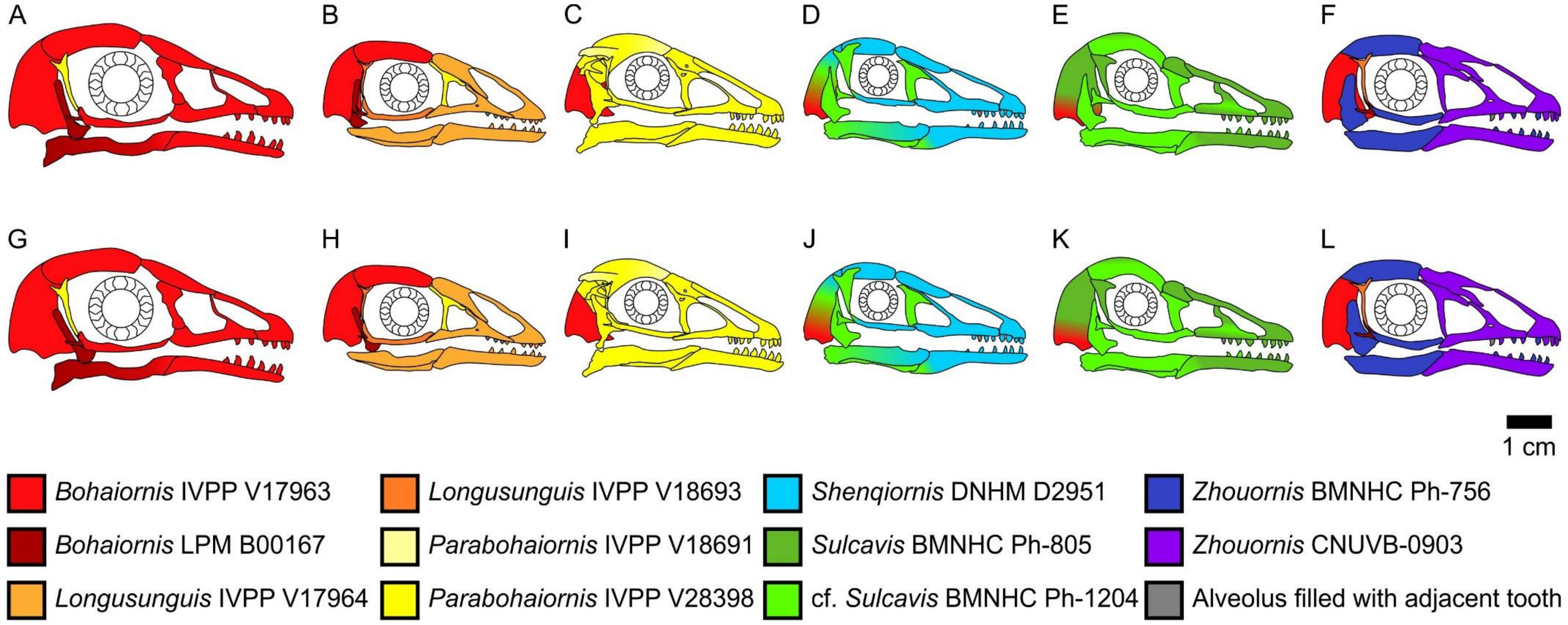

### Supplemental Tables

#### Table S1

Summary of taxa included in Bohaiornithidae. “Bohaiornithidae” is used in an informal term to refer to any distinct clade containing *Bohaiornis*, as the strict clade definition is unstable [24]. If multiple members of the other five originally-defined bohaiornithids [25] resolved near *Bohaiornis*, “Bohaiornithidae” was considered to be the least inclusive clade containing all of them with a tolerance of two bird taxa not in the original six between any two internal nodes.

| Status | |
|---|---|
| "Bohaiornithidae" | + |
| Sister to "Bohaiornithidae" | ~ |
| Not "Bohaiornithidae" | - |
| Not in study | x |

| | *Beiguornis* | *Bohaiornis* | *Longusunguis* | *Parabohaiornis* | *Shenqiornis* | *Sulcavis* | *Zhouornis* | *Gretcheniao* | *Musivavis* | *Eoenantiornis* | *Fortunguavis* | *Linyiornis* |
|---|---|---|---|---|---|---|---|---|---|---|---|---|
| Wang et al [26] | x | + | + | + | + | + | + | x | x | ~ | x | x |
| Wang et al [27] | x | + | + | + | + | + | + | x | x | - | - | x |
| Wang et al [28] | x | + | + | + | + | + | + | x | x | ~ | - | x |
| Wang and Liu [29] | x | + | + | + | + | + | + | x | x | ~ | ~ | x |
| Wang et al [30] | x | + | + | + | + | + | + | x | x | ~ | + | + |
| Hu and O'Connor [23] | x | + | + | + | + | + | + | x | x | - | + | x |
| Wang and Zhou [31] | x | + | + | + | + | + | + | x | x | - | ~ | ~ |
| Cau [32] | x | + | x | x | x | + | + | x | x | x | x | x |
| Chiappe et al [33] | x | + | + | + | - | - | - | - | x | - | - | x |
| Zhang and Wang [34] | x | + | + | + | + | + | + | x | x | x | ~ | ~ |
| Hu et al [35] | x | + | + | + | + | + | - | x | x | - | - | - |
| O'Connor et al [36] | x | + | ~ | + | ~ | ~ | ~ | x | x | ~ | ~ | ~ |
| Pittman et al [37] new technology | x | + | + | + | + | + | + | x | x | - | - | - |
| Pittman et al [37] traditional | x | + | + | + | + | + | + | x | x | ~ | ~ | ~ |

| | | | | | | | | | | | | |
|---|---|---|---|---|---|---|---|---|---|---|---|---|
| Wang and Zhou [38] | x | + | + | + | + | + | + | x | x | - | - | - |
| Li et al [39] strict consensus | x | + | ~ | + | ~ | ~ | ~ | x | x | ~ | x | x |
| Li et al [39] reduced consensus | x | + | + | + | + | + | + | x | x | ~ | x | x |
| Liu et al [24] | x | + | + | + | - | + | + | ~ | x | - | - | x |
| Wang et al [40] | x | + | + | + | + | + | + | - | x | - | - | - |
| Wang et al [41] | + | + | - | + | + | + | + | - | x | + | + | x |
| Wang et al [42] unweighted | x | + | - | x | + | + | + | - | - | + | + | x |
| Wang et al [42] K=20 | x | + | - | - | + | - | - | - | - | - | - | x |
| Wang et al [42] K=5 | x | + | - | - | + | - | + | - | - | + | - | x |

#### Table S2

Diet cut-offs used in this study. Percentages refer to values given in EltonTraits 1.0 [43], with Diet-Tetr being the sum of Diet-Ect and Diet-End (ectotherm and endotherm tetrapod food sources are combined). Granivores were separated into husking and swallowing subdivisions based on feeding descriptions in the literature.

| **Diet** | **Cut-Off** |
|---:|:---:|
| Folivore | 60+% Diet-PlantO |
| Frugivore | 60+% Diet-Fruit |
| Generalist | 40% or less in any category |
| Granivore | 70+% Diet-Seed |
| Invertivore | 60+% Diet-Inv |
| Nectarivore | 60+% Diet-Nect |

| | |
|---|---|
| Piscivore | 50+% Diet-Fish |
| Scavenger | 50+% Diet-Scav |
| Tetrapod Hunter | 60+% Diet-Tetr |

Table S3

Sensitivity analysis of quadrate position (Fig. S11) affecting bohaiornithid predictions diet by FDA from MA and functional indices of extant bird jaws. Values with green backgrounds are more likely, values with red backgrounds are less likely. Compared to results with most likely reconstructions (Table 2), folivory is more likely when the quadrate is shifted anteriorly and piscivory is more likely when the quadrate is shifted posteriorly. Diet abbreviations: GranivoreH, Husking Granivore; GranivoreS, Swallowing Granivore; Tetra Hunt, Tetrapod Hunter.

| **Adjustment** | **Taxon** | **Folivore** | **Frugivore** | **Generalist** | **GranivoreH** | **GranivoreS** | **Invertivore** | **Nectarivore** | **Piscivore** | **Scavenger** | **Tetra Hunt** |
|---|---|---|---|---|---|---|---|---|---|---|---|
| Anterior | *Bohaiornis* | 1.00E+00 | 4.51E-07 | 2.86E-05 | 1.86E-14 | 1.72E-05 | 1.83E-07 | 1.32E-06 | 5.17E-09 | 1.78E-12 | 4.54E-09 |
| | *Longusunguis* | 9.90E-01 | 5.57E-04 | 8.75E-03 | 1.98E-11 | 4.03E-04 | 1.17E-04 | 1.83E-04 | 1.73E-05 | 8.58E-09 | 1.78E-06 |
| | *Parabohaiornis* | 5.53E-05 | 2.36E-07 | 1.92E-03 | 2.97E-07 | 3.54E-05 | 4.06E-07 | 1.97E-09 | 1.37E-05 | 9.98E-01 | 1.14E-05 |
| | *Shenqiornis* | 6.44E-01 | 2.07E-03 | 3.36E-01 | 2.57E-08 | 2.96E-03 | 3.62E-03 | 1.81E-03 | 9.11E-03 | 1.67E-04 | 2.96E-05 |
| | *Sulcavis* | 8.00E-01 | 1.77E-03 | 1.92E-01 | 6.72E-09 | 1.95E-03 | 1.37E-03 | 5.64E-04 | 1.87E-03 | 1.01E-05 | 2.73E-05 |
| | *Zhouornis* | 4.33E-01 | 8.93E-02 | 2.80E-01 | 9.73E-08 | 3.59E-02 | 9.03E-02 | 3.62E-02 | 1.79E-02 | 1.49E-04 | 1.71E-02 |
| Posterior | *Bohaiornis* | 9.97E-01 | 3.38E-05 | 1.20E-03 | 4.70E-12 | 1.22E-03 | 4.75E-05 | 7.74E-05 | 2.00E-06 | 3.22E-09 | 4.15E-07 |
| | *Longusunguis* | 5.55E-04 | 1.91E-01 | 3.21E-01 | 2.48E-06 | 1.56E-03 | 2.74E-01 | 2.62E-02 | 1.78E-01 | 3.90E-04 | 7.80E-03 |
| | *Parabohaiornis* | 6.97E-06 | 5.93E-07 | 1.36E-03 | 1.56E-07 | 3.46E-05 | 1.40E-06 | 3.02E-09 | 2.85E-05 | 9.99E-01 | 7.54E-06 |
| | *Shenqiornis* | 5.21E-04 | 1.03E-02 | 3.93E-01 | 5.36E-06 | 8.81E-04 | 7.07E-02 | 6.17E-03 | 4.91E-01 | 2.74E-02 | 8.14E-04 |
| | *Sulcavis* | 1.52E-03 | 2.32E-02 | 6.21E-01 | 6.20E-06 | 1.61E-03 | 6.78E-02 | 3.56E-03 | 2.74E-01 | 4.98E-03 | 2.03E-03 |
| | *Zhouornis* | 3.09E-05 | 8.31E-02 | 6.40E-02 | 1.55E-05 | 1.76E-03 | 3.91E-01 | 1.94E-02 | 2.74E-01 | 8.34E-03 | 1.59E-01 |

Table S4
Comparison of the bone character stages of Hu and O'Connor [23] with the histological maturity stages of Atterholt et al. [44]. Subadult maturity appears to be obtained by Stage 1 (the most immature fusion stage of any specimen examined in this work), while full skeletal maturity is not obtained until after Stage 3 (the most mature fusion stage). Histological maturities are identified by [44] directly, bone fusion stages are taken from [23] where possible and intuited based on their fusion criteria where not.

| **Specimen** | **Atterholt et al Histological Maturity** | **Hu & O'Connor Character Stage** |
|---|---|---|
| *Parapengornis* IVPP V18687 | young subadult | 1 |
| *Eopengornis* STM24-1 | young subadult | 2a |
| *Monoenantiornis* IVPP V20289 | subadult | 2a |
| *Parvavis* IVPP V18586 | subadult | 2b |
| *Cruralispennia* IVPP V21711 | subadult | 3 |
| *Pterygornis* IVPP V16363 | young adult | 3 |
| *Avimaia* IVPP V25371 | adult | 3 |
| *Mirarce* UCMP139500 | adult | 3 |
| *Mirusavis* IVPP V18692 | adult | 3 |
| STM 29-8 | adult | 3 |
| *Zhouornis* CNUVB-0908 | adult | 3 |

Table S5
Qualitative dietary hypotheses for enantiornithine taxa. The hypothesised diet listed may overgeneralise the hypothesis of the original publication for the sake of brevity. Most notably we often simplify hypotheses like "carnivory taking small animals" to invertivory, as when these authors provide analogous extant birds for context, they predominantly were invertivorous birds.

| **Taxon** | **Hypothesized Diet** | **Reference** |
|---|---|---|
| *Avisaurus archibaldi* | Raptorial | [9] |
| *Bohaiornis guoi* | Durophagy, invertivory, raptorial | [11 pg. 270, 12, 45, 46] |
| *Boluochia zhengi* | Piscivory, raptorial | [12, 47, 48, 49 pg. 152, 50] |

| | | |
|---|---|---|
| *Brevirostruavis macrohyoideus* | Invertivory, nectarivory | [12, 39] |
| *Cathayornis yandica* | Invertivory | [9] |
| *Concornis lacustris* | Invertivory | [9] |
| *Cuspirostrisornis houi* | Raptorial | [9] |
| *Chiappeavis magnapremaxillo* | Invertivory | [12 pg. 149] |
| *Dapingfangornis sentisorhinus* | Piscivory | [9] |
| *Falcatakely forsterae* | Frugivory | [12] |
| *Gettyia gloriae* | Scavenging | [12 pg. 179] |
| *Gobipipus reshetovi* | Granivory, invertivory | [10] |
| *Gobipteryx minuta* | Granivory, invertivory | [10, 12] |
| *Halimornis thompsoni* | Piscivory | [9, 12] |
| *Longipteryx chaoyangensis* | Piscivory | [5-10, 11 pg. 83 and 270, 12] |
| *Longirostravis hani* | Invertivory, probing | [9, 10, 11 pg. 270, 12, 51] |
| *Mirarce eatoni* | Raptorial | [12] |
| *Neuquenornis volans* | Raptorial | [9, 10, 12, 50] |
| *Parabohaiornis martini* | Invertivory | [11 pg. 270] |
| *Pengornis houi* | Hard invertivory, soft invertivory | [7, 8, 9 pg. 136, 10, 11 pg. 83, 12] |
| *Rapaxavis pani* | Invertivory, probing | [9, 10, 11 pg. 270, 12] |
| *Shanweiniao cooperorum* | Invertivory, probing | [11 pg. 270, 12] |
| *Shengjingornis yangi* | Invertivory, probing | [12] |
| *Shenqiornis mengi* | Durophagy, invertivory | [8, 9, 12, 52] |
| *Sinornis santensis* | Carnivory, folivory | [9, 49 pg. 150, 50] |
| *Soroavisaurus australis* | Raptorial | [9, 10, 50] |
| *Sulcavis geeorum* | Durophagy | [11 pg. 83, 12, 53] |

| | | |
|---|---|---|
| *Vescornis hebeiensis* | Invertivory | [9] |
| *Zhouornis hani* | Invertivory | [11 pg. 270] |

Table S6

K (mass) and $K_{mult}$ (all others) values for each extant dataset investigated. $K_{mult}$ values are calculated from the same datasets as [3], with an updated phylogeny. These $K_{mult}$ values are similar to those reported in [3], if slightly lower. Body mass is an exception, as its extant dataset has been greatly expanded here and the resulting K value is much lower than in [3]. A K/$K_{mult}$ value of 1 indicates trait variation distributed as expected under a BM model of evolution; values below 1 indicate closely-related species are less similar than expected by BM, values above 1 that they are more similar. Three asterisks (***) indicates significance at the p = 0.001 level.

| **Dataset** | $\mathbf{K_{mult}}$ | **p-value** |
|---|---|---|
| Mass | 0.932 | 0.001*** |
| MA | 0.628 | 0.001*** |
| FEA | 0.252 | 0.503 |
| pedal TM | 0.477 | 0.001*** |
| skull TM | 0.898 | 0.001*** |

Table S7

K values for individual MA and functional index variables of the extant skull dataset herein. The updated phylogeny used here returned generally lower K values than those reported in [3]. A K value of 1 indicates trait variation distributed as expected under a BM model of evolution; values below 1 indicate closely-related species are less similar than expected by BM, values above 1 that they are more similar. Three asterisks (***) indicates significance at the p = 0.001 level.

| Jaw | Variable | K | p-value |
|---|---|---|---|
| Upper | AMA | 0.5635082 | 0.001*** |
| | PMA | 0.44203 | 0.001*** |
| | OMA | 0.3531941 | 0.001*** |
| | AO | 0.7773561 | 0.001*** |
| | MCH | 0.7805066 | 0.001*** |

| | | | |
|---|---|---|---|
| | ACH | 1.1471418 | 0.001*** |
| Lower | AMA | 0.7289534 | 0.001*** |
| | PMA | 0.4987183 | 0.001*** |
| | OMA | 0.5175259 | 0.001*** |
| | AO | 0.7274349 | 0.001*** |
| | MMH | 0.5001836 | 0.001*** |
| | AMH | 0.6273753 | 0.001*** |

Table S8

Significant differences between extant diet groups, based on phylogenetic HSD of MA and functional indices. The updated phylogeny used here does not change any groups' significant difference from one another from [3]. One asterisk (*) indicates significance at the $p < 0.05$ level, two asterisks (**) indicates significance at the $p < 0.01$ level, and three asterisks (***) indicates significance at the $p = 0.001$ level. Diet abbreviations: GranivoreH, husking granivore; GranivoreS, swallowing granivore; Tetra Hunt, tetrapod hunter.

| | Folivore | Frugivore | Generalist | GranivoreH | GranivoreS | Invertivore | Nectarivore | Piscivore | Scavenger | Tetra Hunt |
|---|---|---|---|---|---|---|---|---|---|---|
| Folivore | | 0.112 | 0.267 | 0.034* | 0.536 | 0.058 | 0.058 | 0.001*** | 0.006** | 0.548 |
| Frugivore | 0.112 | | 0.468 | 0.228 | 0.851 | 0.203 | 0.099 | 0.002** | 0.151 | 0.406 |
| Generalist | 0.267 | 0.468 | | 0.007** | 0.859 | 0.254 | 0.081 | 0.001*** | 0.01* | 0.786 |
| GranivoreH | 0.034* | 0.228 | 0.007** | | 0.148 | 0.001*** | 0.003** | 0.001*** | 0.005** | 0.018* |
| GranivoreS | 0.536 | 0.851 | 0.859 | 0.148 | | 0.658 | 0.164 | 0.014* | 0.154 | 0.739 |
| Invertivore | 0.058 | 0.203 | 0.254 | 0.001*** | 0.658 | | 0.262 | 0.001*** | 0.034* | 0.766 |
| Nectarivore | 0.058 | 0.099 | 0.081 | 0.003** | 0.164 | 0.262 | | 0.489 | 0.49 | 0.392 |
| Piscivore | 0.001*** | 0.002** | 0.001*** | 0.001*** | 0.014* | 0.001*** | 0.489 | | 0.283 | 0.039* |
| Scavenger | 0.006** | 0.151 | 0.01* | 0.005** | 0.154 | 0.034* | 0.49 | 0.283 | | 0.132 |
| Tetra Hunt | 0.548 | 0.406 | 0.786 | 0.018* | 0.739 | 0.766 | 0.392 | 0.039* | 0.132 | |

Table S9

Significant differences between extant diet groups, based on phylogenetic HSD of FEA intervals data. The updated phylogeny used here leads to less significant differences than found in [3]: folivores are no longer significantly different from husking granivores or scavengers; frugivores significantly different from scavengers; or husking granivores significantly different from invertivores. One asterisk (*) indicates significance at the $p < 0.05$ level, two asterisks (**) indicates significance at the $p < 0.01$ level, and three asterisks (***) indicates significance at the $p = 0.001$ level. Diet abbreviations: GranivoreH, husking granivore; GranivoreS, swallowing granivore; Tetra Hunt, tetrapod hunter.

| | Folivore | Frugivore | Generalist | GranivoreH | GranivoreS | Invertivore | Nectarivore | Piscivore | Scavenger | Tetra Hunt |
|---|---|---|---|---|---|---|---|---|---|---|
| Folivore | | 0.405 | 0.291 | 0.08 | 0.088 | 0.036* | 0.097 | 0.377 | 0.061 | 0.212 |
| Frugivore | 0.405 | | 0.981 | 0.531 | 0.464 | 0.522 | 0.363 | 0.94 | 0.068 | 0.676 |
| Generalist | 0.291 | 0.981 | | 0.226 | 0.419 | 0.102 | 0.246 | 0.918 | 0.011* | 0.684 |
| GranivoreH | 0.08 | 0.531 | 0.226 | | 0.311 | 0.228 | 0.315 | 0.52 | 0.035* | 0.388 |
| GranivoreS | 0.088 | 0.464 | 0.419 | 0.311 | | 0.928 | 0.781 | 0.833 | 0.011* | 0.82 |
| Invertivore | 0.036* | 0.522 | 0.102 | 0.228 | 0.928 | | 0.702 | 0.661 | 0.006** | 0.359 |
| Nectarivore | 0.097 | 0.363 | 0.246 | 0.315 | 0.781 | 0.702 | | 0.399 | 0.021* | 0.601 |
| Piscivore | 0.377 | 0.94 | 0.918 | 0.52 | 0.833 | 0.661 | 0.399 | | 0.025* | 0.661 |
| Scavenger | 0.061 | 0.068 | 0.011* | 0.035* | 0.011* | 0.006** | 0.021* | 0.025* | | 0.019* |
| Tetra Hunt | 0.212 | 0.676 | 0.684 | 0.388 | 0.82 | 0.359 | 0.601 | 0.661 | 0.019* | |

Table S10

K values for individual variables used in pedal TM analyses. The updated phylogeny used here returned generally lower K values than those reported in [3]. A K value of 1 indicates trait variation distributed as expected under a BM model of evolution; values below 1 indicate closely-related species are less similar than expected by BM, values above 1 that they are more similar. Two asterisks (**) indicates significance at the $p < 0.01$ level and three asterisks (***) indicates significance at the $p = 0.001$ level.

| Variable | K | p-value |
|---|---|---|
| DI/DIII ratio | 0.7818675 | 0.001*** |
| DII/DIII ratio | 0.6489054 | 0.001*** |
| DIV/DIII ratio | 0.4109582 | 0.001*** |

| | | |
|---|---|---|
| DI angle | 0.643955 | 0.001*** |
| DII angle | 0.6129209 | 0.001*** |
| DIII angle | 0.3919881 | 0.001*** |
| DIV angle | 0.3153363 | 0.010** |

Table S11

Significant differences between extant pedal ecology groups, based on phylogenetic HSD of pedal TM data. The updates to phylogeny and additional extant data used here do not change any groups' significant difference from one another from [3]. One asterisk (*) indicates significance at the $p < 0.05$ level and two asterisks (**) indicates significance at the $p < 0.01$ level. Category abbreviations: Large Raptor, raptor taking prey which does not fit in the foot; Small Raptor, raptor taking prey which can fit in the foot.

| | Ground | Perch | Large Raptor | Small Raptor | Shrike |
|---|---|---|---|---|---|
| Ground | | 0.006** | 0.007** | 0.072 | 0.027* |
| Perch | 0.006** | | 0.995 | 0.414 | 0.433 |
| Large Raptor | 0.007** | 0.995 | | 0.026* | 0.295 |
| Small Raptor | 0.072 | 0.414 | 0.026* | | 0.388 |
| Shrike | 0.027* | 0.433 | 0.295 | 0.388 | |

Table S12

K values for individual variables used in skull TM analyses. Relative Skull Length is defined mathematically in the Methods of the main text. A K value of 1 indicates trait variation distributed as expected under a BM model of evolution; values below 1 indicate closely-related species are less similar than expected by BM, values above 1 that they are more similar. Three asterisks (***) indicates significance at the $p = 0.001$ level.

| Variable | K | p-value |
|---|---|---|
| Skull Length | 0.8925226 | 0.001*** |
| Rostrum Length | 0.9059976 | 0.001*** |
| $LOG_{10}$ Rostrum Length / $LOG_{10}$ Skull Length | 0.6427194 | 0.001*** |
| Relative Skull Length | 1.3161350 | 0.001*** |
